# Immune checkpoint inhibitors amplify type 2 immune mediated repair by pro-regenerative scaffolds

**DOI:** 10.64898/2026.01.31.703034

**Authors:** Jordan Garcia, Anna Ruta, Frank Haoning Yu, Joscelyn C. Mejías, Alexis N. Peña, Natalie Rutkowski, Elise F. Gray-Gaillard, Constance Dubois, Christopher Cherry, Maria Browne, Katlin Stivers, David R. Maestas, Kavita Krishnan, Alexander T.F. Bell, Elana J. Fertig, Carisa M. Cooney, Damon Cooney, Patrick Byrne, Alexander T. Hillel, Kellie N. Smith, Hongkai Ji, Robert Anders, Drew M. Pardoll, Jennifer H. Elisseeff

## Abstract

Extracellular matrix (ECM) scaffolds induce type 2 immunity to promote repair. Here, we show that immune cells recruited to ECM-treated murine muscle injuries and clinical soft tissue defects express immune checkpoints. Specifically, T_H_2 cells and regulatory T cells (Tregs) increase LAG3 expression, while macrophages express PDL2. TCR analysis and a triple-reporter strain for interleukin (IL)-13 and Treg fate-mapping suggest that Tregs in ECM-treated wounds transition into T_H_2-like exTregs that express LAG3. Immune checkpoint inhibition (ICI) significantly stimulated type 2 immunity in ECM-treated wounds, including increased T_H_2 cells, Treg transition to T_H_2-like exTregs, and pro-regenerative macrophages. Moreover, ICI enhanced muscle repair and reduced fibrosis in ECM-treated wounds. Collectively, these findings show Treg/T_H_2 plasticity in wound healing and introduce a novel ICI application to enhance immune-mediated regeneration.

## Main Text

Immune checkpoint inhibition (ICI) has revolutionized the field of oncology, enhancing the activity of tumor-reactive effector T cells and re-activating previously exhausted T cells to combat a growing list of cancer types.^1^ Analogous applications of immune checkpoint-targeting therapies are under development for chronic infections such as malaria, HIV, and tuberculosis.^2^ Conversely, activating immune checkpoint pathways has shown promise as a treatment strategy in autoimmune conditions and to enhance survival of vascularized composite allografts.^3,4^ ICI has even been leveraged to improve the effects of aging in different tissues by targeting age-related inflammation and cellular senescence.^5^

The role of immune checkpoints on regulation of T cell responses in the wound environment and to reconstructive biomaterials remains unknown. The burden of chronic wounds following trauma, infection, and cancer resection persists despite recent advances in wound care.^6,7^ During wound healing, the inflammatory, proliferative, and remodeling phases govern either the recovery of functional tissue or progression to chronic unresolved wounds or fibrosis-induced dysfunction.^8^ Multiple immune cell types contribute to recovery following injury, with T cells emerging as key mediators in the tissue damage response.^9,10^ A myriad of factors such as pre-existing cancer, infection, inflammation, metabolic dysfunction, advanced age, and cellular senescence can influence T cell phenotype and thus outcomes of tissue injury and repair.^11–15^ We previously demonstrated that biological scaffolds derived from the extracellular matrix (ECM) promote T_H_2 CD4^+^ T cells that together with interleukin (IL)-4-secreting eosinophils and M2-like macrophages induce pro-regenerative tissue environments that support wound healing.^16^ Other regenerative therapies leveraging T_H_2 cells utilize soluble helminth egg derivatives which have been tested in muscle, cornea, and cartilage repair models.^17^ Understanding how biomaterials modulate immune pathways to influence effector T cell phenotypes could lead to strategies to augment their beneficial effect on healing.

Here, we show that ECM-based wound therapies promote immune checkpoint expression, in particular LAG3 and PDL2, and increase sensitivity to ICI. We find that the characteristic type 2 immune response to ECM scaffolds develops, in part, by regulatory T cells (Tregs) transitioning to IL13-producing “T_H_2-like exTregs”. We show that combination ICI treatment regimens targeting PD1, CTLA4, and LAG3 further expand IL10^+^ T_H_2 cells, promote Treg transition to T_H_2-like exTregs, and augment the pro-regenerative effect of ECM implants in muscle wounds. These results shed light on Treg-T_H_2 cell dynamics during tissue repair and suggest a novel application of ICI for increasing the pro-regenerative effects of biomaterial-based wound therapies.

### ECM scaffolds promote T cell and macrophage expression of immune checkpoints LAG3 and PDL2

To characterize immune checkpoint expression in response to tissue injury, we performed volumetric muscle loss (VML) injury in murine quadricep muscles and treated the wound sites with saline (injury only control) or pro-regenerative biological ECM scaffolds. Transcriptomic profiling of bulk muscle tissue 1-week post injury using the NanoString nCounter PanCancer Immune Profiling Panel revealed significantly higher expression of immune checkpoint genes in ECM-treated VML injuries including *Pdcd1lg1* (encodes PDL1)*, Pdcd1lg2* (encodes PDL2), and *Ctla4* (**Fig. 1A**). Since T cells are a smaller subset of immune cells in VML wounds, we performed a similar transcriptomic analysis on isolated T cells (Live CD45^+^CD11b^-^CD3^+^) and found that ECM treatment significantly increased T cell expression of *Lag3*, *Tnfrsf18* (encodes GITR), *Tigit*, *Ctla4, Pdcd1lg1*, and *Pdcd1lg2* (**Fig. 1B**). ECM treatment of the VML injury promoted expression of genes related to type 2 immunity and Tregs, which are consistent with the pro-regenerative immune response previously linked to biologically derived materials. Specifically, the expression of type 2-related genes *Arg1* and *Il1rl1* significantly increased in bulk tissue analysis of ECM-treated injuries compared to saline-treated controls while the type 17-related gene *Rorc* decreased (**Fig. S1A**). In the sorted T cells, expression of T_H_2-related (*Arg1*, *Il5, Il13, Il1rl1,* and *Gata3*) and Treg-related (*Il10* and *Foxp3*) genes significantly increased in the ECM-treated condition compared saline-treated controls (**Fig. S1B**).

**Fig. 1.**
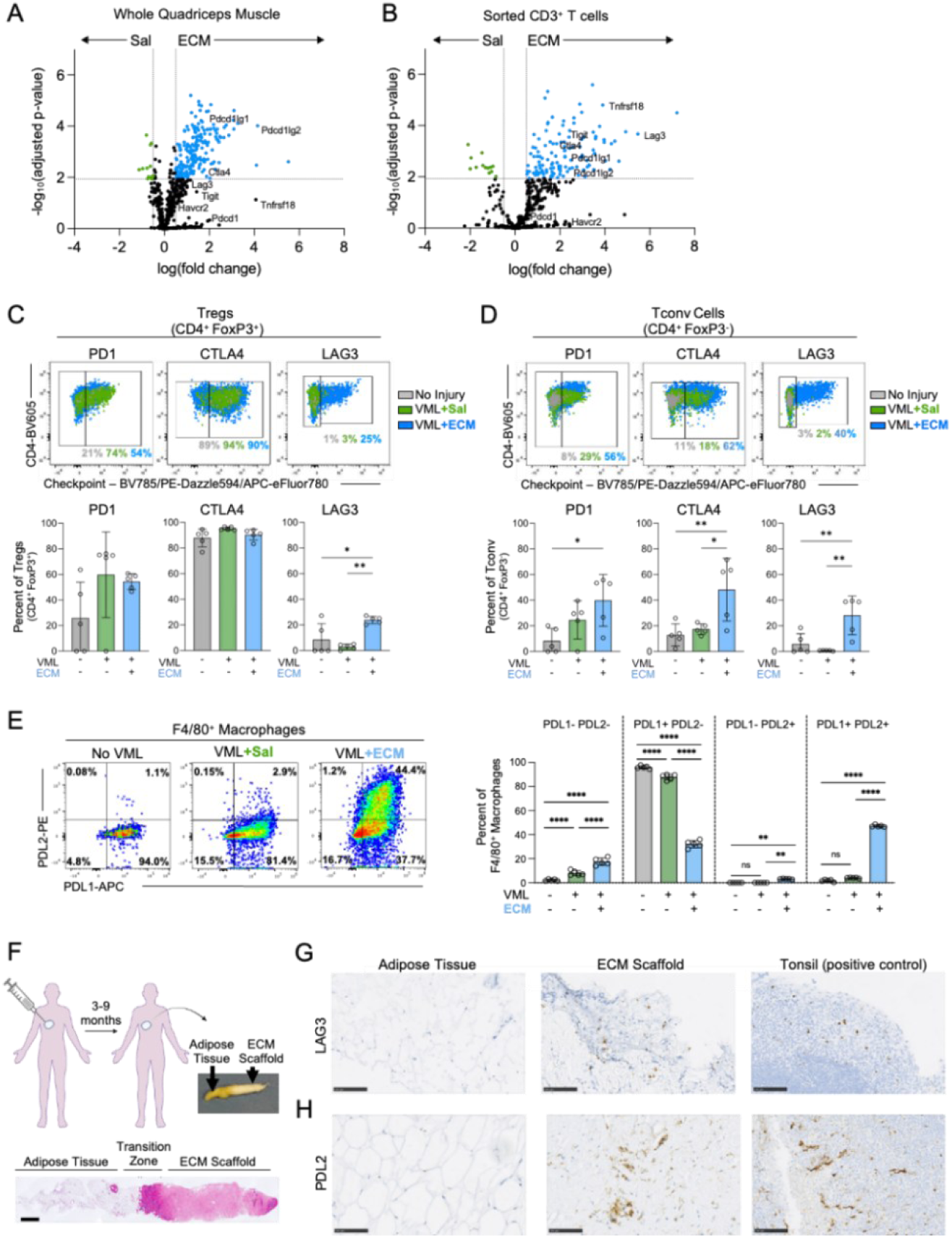
Wound treatment with ECM scaffolds promotes expression of LAG3 and PDL2 in CD4^+^ T cells and macrophages, respectively. Volumetric muscle loss (VML) injury was performed in murine quadriceps muscle and treated with either saline (Sal, injury only control) or extracellular matrix (ECM) scaffold. **(A)** Differential gene expression between Sal-and ECM-treated VML injuries performed using NanoString nCounter PanCancer Immune Profiling Panel on whole quadriceps muscle or **(B)** sorted CD3^+^ T cells at 1-week post injury. Genes encoding immune checkpoints are labeled. **(C)** Representative flow cytometry plots and quantification of PD1, CTLA4, and LAG3 expression by Tregs (CD4^+^FoxP3^+^) or **(D)** conventional CD4^+^ T cells (CD4^+^FoxP3^-^) in uninjured muscle, Sal-treated VML, and ECM-treated VML at 1-week. **(E)** Representative flow cytometry plots and quantification of PDL1 and PDL2 expression by macrophages (CD11b^+^ F4/80^+^) in uninjured muscle, Sal-treated VML, and ECM-treated VML at 1-week. **(F)** Acellular adipose tissue (AAT) ECM implants were biopsied 3-9 months after injection in patients enrolled in clinical trial NCT03544632. Representative gross pathology and H&E staining (scale bar: 1 mm). **(G)** Representative immunohistochemistry staining of LAG3 or **(H)** PDL2 in native adipose tissue (negative control), AAT ECM, and tonsil (positive control) (scale bar:100mm. Data is presented as mean±SD and analyzed using one-way ANOVA with Tukey’s multiple comparisons test (C-E). NS p>0.05; * p<0.05; ** p<0.01; *** p<0.001; **** p<0.0001.

We further investigated immune checkpoint protein expression in the various T cell subsets in response to biomaterial treatment by using high-parameter flow cytometry. Tregs (CD4^+^FoxP3^+^) in uninjured as well as saline-and ECM-treated VML injured muscles ubiquitously expressed CTLA4 and GITR (**Fig. 1C, S2A**). While injury increased Treg expression of PD1, TIGIT, and TIM3 irrespective of treatment, LAG3 was the only analyzed checkpoint uniquely upregulated in ECM-treated wounds. The conventional CD4^+^ T cells (CD4^+^FoxP3^-^) similarly increased expression of PD1, TIM3, and GITR in response to injury, but only upregulated CTLA4, LAG3, and TIGIT expression with ECM treatment (**Fig. 1D, S2B**). CD8^+^ T cells increased expression of PD1, CTLA4, and GITR with injury, while their low expression of LAG3, TIGIT, and TIM3 was unchanged (**Fig. S2C**). Notably, CD8^+^ T cell expression of immune checkpoints did not significantly change with ECM treatment. A time course analysis comparing various post injury time points (3-day, 1-week, and 3-weeks) revealed that LAG3 expression peaks on CD4^+^ T cells 1-week after injury and subsequently decreases while PD1 expression remains elevated (**Fig. S2D**). Previous work by our group demonstrated that both allogeneic and xenogeneic ECM sources elicit robust T_H_2 immune responses.^18^ The ECM scaffolds utilized in these experiments were porcine-derived, so to determine if checkpoint expression is driven by xenogeneic ECM only, VML injuries were treated with an allogeneic ECM composed of murine type I/III collagen. Even with allogeneic ECM, LAG3 expression was induced in CD4^+^ T cells 1-week after injury, albeit to a less extent than with xenogeneic ECM (**Fig. S2E**).

Since T_H_2 cells are known to upregulate PDL2 expression in macrophages^19^, we further evaluated immune checkpoint ligand expression on myeloid cells in the injury and biomaterial microenvironments. We characterized both PD1 ligands, PDL1 and PDL2, given their increased expression with ECM treatment in our NanoString dataset and their growing clinical relevance in ICI response.^20^ First, we analyzed our previously published single-cell RNA-sequencing dataset^21^ that compared saline-and ECM-treated VML injuries and found increased transcript levels of *Pdcd1Ig1* and *Pdcd1lg2* in myeloid clusters with ECM treatment (**Fig. S3A**). Specifically, we found increased expression of *Pdcd1lg2* in the previously defined regenerative “R2” macrophage subset that is enriched in ECM-treated injuries.^22^ We subsequently validated checkpoint ligand expression using flow cytometry which confirmed that ECM-treated muscle injury, but not saline-treated, induced PDL2 expression in F4/80^+^ macrophages (**Fig. 1E**). While monocytes also upregulated PDL2 in response to ECM treatment, eosinophils did not (**Fig. S3B**). Collectively, these findings confirm that biomaterial implants can modulate immune checkpoint expression, suggesting biomaterial immune responses may be sensitive to ICI.

Finally, we investigated if human immune responses to clinical ECM-based scaffolds also increase immune checkpoint expression and mirror our murine ECM-treated injury model. We obtained biopsies from patients who had soft-tissue defects injected with ECM scaffolds derived from decellularized human adipose tissue as part of a phase 2 clinical trial (NCT03544632).^18^ Using an ultrasound-guided core-needle biopsy technique, samples were obtained 3-9 months following ECM injection (**Fig. 1F**). Morphologically, hematoxylin and eosin (H&E) stained tissue biopsies showed eosinophilic changes alongside native adipocytes with a transitional region rich in mononuclear cellular infiltrate. Given LAG3 and PDL2 were immune checkpoints most unique to the ECM-treated murine VML injuries, we performed immunohistochemistry for these checkpoints which showed robust positive staining in all 7 patient biopsies (**Fig. 1G, 1H, S4A**). Positive detection of LAG3 and PDL2 was confined to the ECM region of the biopsy and was not observed in the adjacent native adipose tissue. The observed cellular staining patterns matched those in human tonsil samples (positive controls) and were verified by an experienced histopathologist.

### ECM scaffolds promote LAG3 expression and expansion of Treg and T_H_2 populations

To further characterize the phenotype of the CD4^+^ T cell subsets in the ECM implant response, we isolated T cells (Live CD45^+^ CD3^+^) using fluorescence activated cell sorting (FACS) from saline-and ECM-treated VML wounds at 1-week post-injury and performed 5’ paired single-cell RNA-and T cell receptor (TCR)-sequencing (scRNA/TCRseq) using 10x Genomics. We captured a total of 37,580 high-quality T cells, including 32,502 from ECM-treated mice (n=3) and 5,078 from saline-treated mice (n=14). Clustering generated seven distinct T cell clusters (**Fig. 2A, S5A, S5B**), including clusters that we identified as T_H_2 (*Gata3, Areg,* and *Il4*) and Treg (*Foxp3*, *Il2ra*, and *Ctla4*) based on expression of canonical marker genes (**Fig. S5C**). Following sub-cluster analysis of the Mixed T cell cluster, we identified T_H_1 (*Tbx21* and *Ifng*) and T_H_17 (*Rorc*, *Il17a*, and *Il17f*) sub-populations (**Fig. S6A**).

**Fig. 2.**
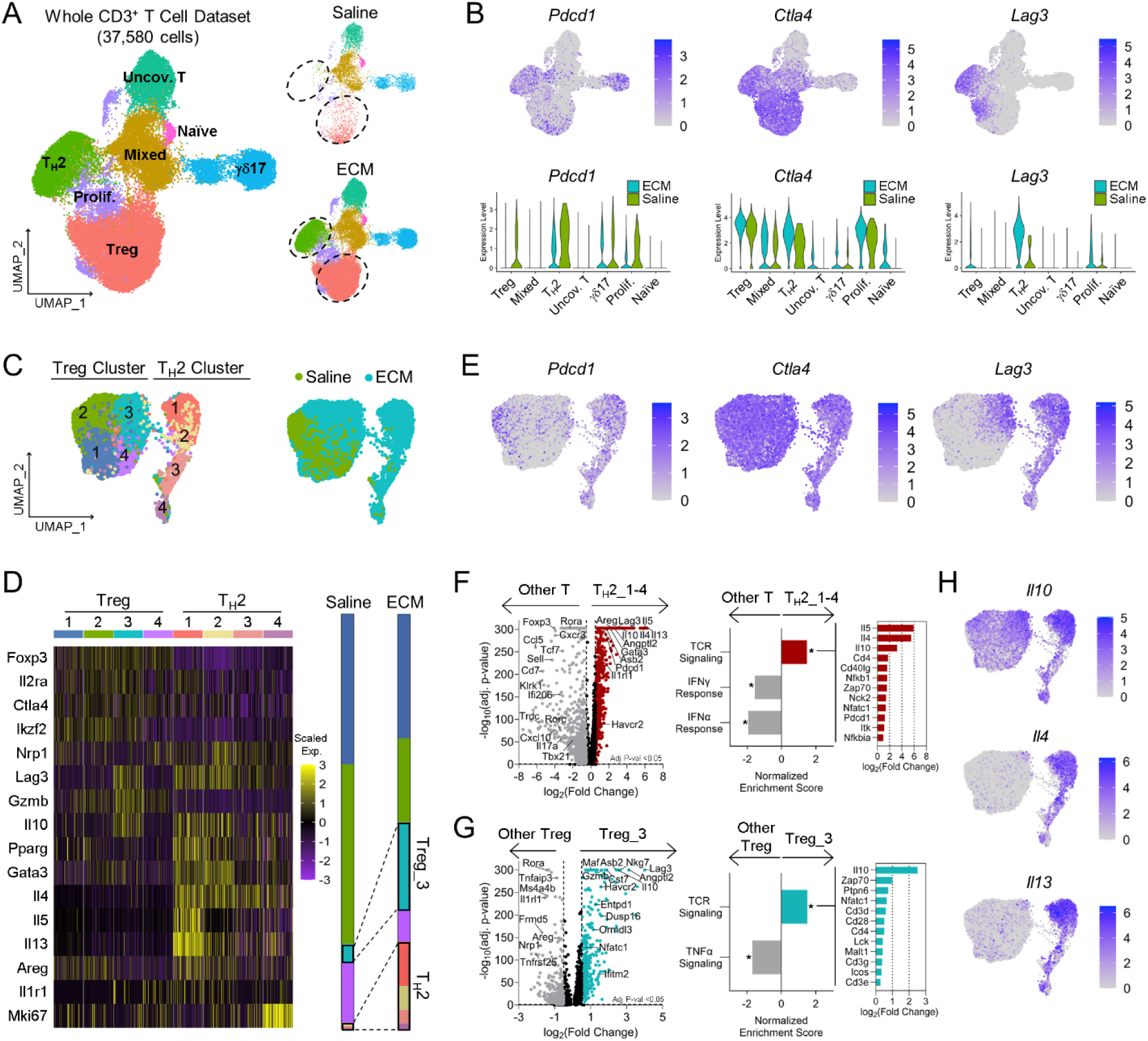
Tregs and T_H_2 cells enriched in ECM-treated wounds express LAG3 and other immune checkpoints. CD3^+^ T cells were FACS isolated from saline-treated (n=14) and ECM-treated (n=3) VML injuries at 1-week for paired single-cell RNA/TCR sequencing. **(A)** Overall UMAP of the main CD3^+^ T cell clusters and their distribution in saline-and ECM-treated VML. Dotted circles label the CD4 and Treg clusters enriched in ECM-treated VML. **(B)** Expression of immune checkpoint genes *Pdcd1*, *Ctla4*, and *Lag3* depicted on feature plots and violin plots split by saline-and ECM-treated VML condition per CD3^+^ T cell cluster. **(C)** UMAP of Treg and T_H_2 sub-clusters and their distribution in saline-(green) and ECM-treated (blue) VML. **(D)** Scaled expression heatmap of canonical Treg and T_H_2 marker genes by sub-clusters. Stacked bar plot comparing the relative abundance of Treg and T_H_2 sub-clusters in saline-and ECM-treated VML. Black outlines label the Treg_3 and T_H_2 cells enriched in ECM-treated VML. **(E)** Feature plots of immune checkpoint (*Pdcd1*, *Ctla4*, *Lag3*) gene expression in Treg and T_H_2 clusters. **(F)** Differential gene expression and gene set enrichment analysis (GSEA) comparing T_H_2 cells to all other T cells or **(G)** Treg_3 to all other Tregs. Top 12 leading edge genes in TCR signaling pathways ranked by relative fold change. **(H)** Ordered feature plots of cytokine (*Il10*, *ll4*, *Il13*) gene expression in Treg and T_H_2 clusters.

The T_H_2 and Treg clusters were enriched in ECM-treated wounds (**Fig. 2A**) and exhibited high expression of *Ctla4* and *Lag3* (**Fig. 2B**). Subcluster analysis identified four main T_H_2 subclusters that all expressed some combination of the canonical T_H_2 cytokines (*Il4, Il5, and Il13*) (**Fig. 2C, 2D, S6B**). All T_H_2 subclusters exhibited notable expression of *Lag3, Pdcd1* (encodes PD1), and *Ctla4* (**Fig. 2E**). Relative to all other T cells, the T_H_2 cells exhibited higher expression of *Areg*, *Il1rl1, Asb2, Dusp16,* and *Ifitm2*, which are shown to be necessary for systemic T_H_2-mediated responses in allergy, helminth infections, and adipose tissue remodeling (**Fig. 2F**).^23^ Based on gene set enrichment analysis (GSEA), the T_H_2 cells were enriched in TCR signaling pathways and decreased in Interferon (IFN)-α and IFNγ response pathways (**Fig. 2F**).

ECM treatment expanded the Treg cluster and sub-cluster analysis revealed a Treg subset unique to the ECM condition (**Fig. 2C, S7A**). Three Treg subclusters (1, 2, and 4) were found in both ECM-and saline-treated conditions, while subcluster 3 (Treg_3) notably expanded in ECM-treated samples. In addition to the canonical Treg genes (*Foxp3*, *Il2ra*, and *Ctla4*) expressed by all Treg subclusters, Treg_2 expressed adipose-associated genes (*Pparg*), whereas Treg_4 expressed higher levels of *Rorc* suggesting a relationship with T_H_17 cells (**Fig. S7A**).^24^ Since Treg_3 was dependent on the presence of ECM implants, we further explored its transcriptomic profile by directly comparing it to all other Tregs (**Fig. 2G**). Treg_3 expressed higher levels of *Il10* and *Gzmb,* consistent with a more activated, immunosuppressive Treg state. Notably, the ECM-specific Treg_3 also had significantly higher expression of *Lag3*. GSEA showed enrichment in TCR signaling and decreased TNFα signaling in Treg_3 (**Fig. 2G**). Additionally, the expression of *Nrp1* and *Ikzf2* (encodes Helios) has been proposed to distinguish thymic-derived Tregs (tTregs) from Tregs derived from conventional T cells, termed ‘peripheral’ Tregs (pTregs). While the relative expression levels of *Nrp1* and *Ikzf2* in tTregs and pTregs is still debated, low *Nrp1* and *Ikzf2* expression is often associated with pTregs.^25–29^ The ECM-specific Treg_3 had lower expression of *Nrp1* and *Ikzf2* (log_2_(Fold Change):-1.79 and-0.46, respectively) relative to the other Tregs, thereby suggesting peripheral rather than thymic origin (**Fig. 2G, S7B**). Altogether, these results suggest that ECM scaffolds recruit Treg subsets of both peripheral and thymic origin.

We recognized that Treg_3 significantly increased expression of genes associated with T_H_2 cell differentiation and propagation (*Asb2, Dusp16, Ifitm2, Ormdl3, Maf, and Gata3*) (**Fig. 2G, S7C**), potentially indicating phenotypic plasticity of Tregs.^30–35^ Conversely, Treg_3 decreased expression of genes associated with suppression of T_H_2 immune responses and activation of Tregs (*Tnfrsf25, Frmd5, Ms4a4b, Rora, and Tnfaip3*).^36–41^ Feature plots for these T_H_2-related and Treg-related genes demonstrate inverse spatial distributions over the Treg subclusters (**Fig. S7D**). Further, feature plots of the canonical T_H_2 cytokines *Il4* and *Il13* reveal expression in Treg subclusters, including the ECM-specific Treg_3 (**Fig. 2H**). Notably, the T_H_2 subsets and Treg_3 subcluster exhibited the highest level of *Il10* expression.

### ECM scaffolds promote Treg plasticity and Treg transition towards T_H_2-like exTregs

Given the transcriptional similarities between Treg_3 and T_H_2 cells, we hypothesized that ECM treatment promotes Treg phenotypic plasticity and the transition to T_H_2-like exTregs. Prior studies show that exTregs express higher levels of *Cst7* and *Nkg7,* which are both significantly increased in Treg_3 (**Fig. S8A**).^42,43^ Additionally, pTregs are more likely to become exTregs, thus Treg_3 (*Nrp1*^low^*Ikzf2*^low^) may be of peripheral origin.^27^ Given these observations, we performed TCR analysis to look for potential TCR clonal sharing between the T_H_2 and Treg subclusters in the ECM-treated condition. The paired nature of the scRNA/TCRseq dataset allows each TCR sequence to be attributed to an individual cell with its associated transcriptome and cluster identity. Complete TCR clones were strictly defined based on cells with matching paired α and β chains, identified by their V(D)J gene usage and CDR3 nucleotide sequences. In cases where only one chain (α or β) was recovered, the sequences were retained as partial TCR clones. ECM treatment resulted in expansion of the TCR repertoire of Tregs as well as T_H_2 cells (**Fig. S8B**). We detected 21 overlapping TCR sequences (10 complete and 11 partial) in Tregs or T_H_2 cells across the independent ECM-treated mice, however none of these shared sequences were present in all three of the ECM-treated mice.

Based on transcriptional clustering, we identified Tregs that shared TCR sequences with cells in the T_H_1, T_H_2, and T_H_17 subclusters. We found that Treg_3 and T_H_2 shared the greatest number of clones, both in terms of unique TCR counts and cell counts, in all three ECM-treated biological replicates (**Fig. S8C**). While there was minimal clonal overlap between Treg_1 or Treg_2 with the T_H_1, T_H_2, and T_H_17 subclusters, Treg_4 (exhibited higher *Rorc* expression relative to other Treg subclusters) shared a significant number of clones with T_H_17. Further analysis considering T_H_2 subclusters demonstrated Treg_3 shared the most clones with T_H_2_1, although there was clonal sharing with each of the T_H_2 subclusters (**Fig. 3A, S8D**).

**Fig. 3.**
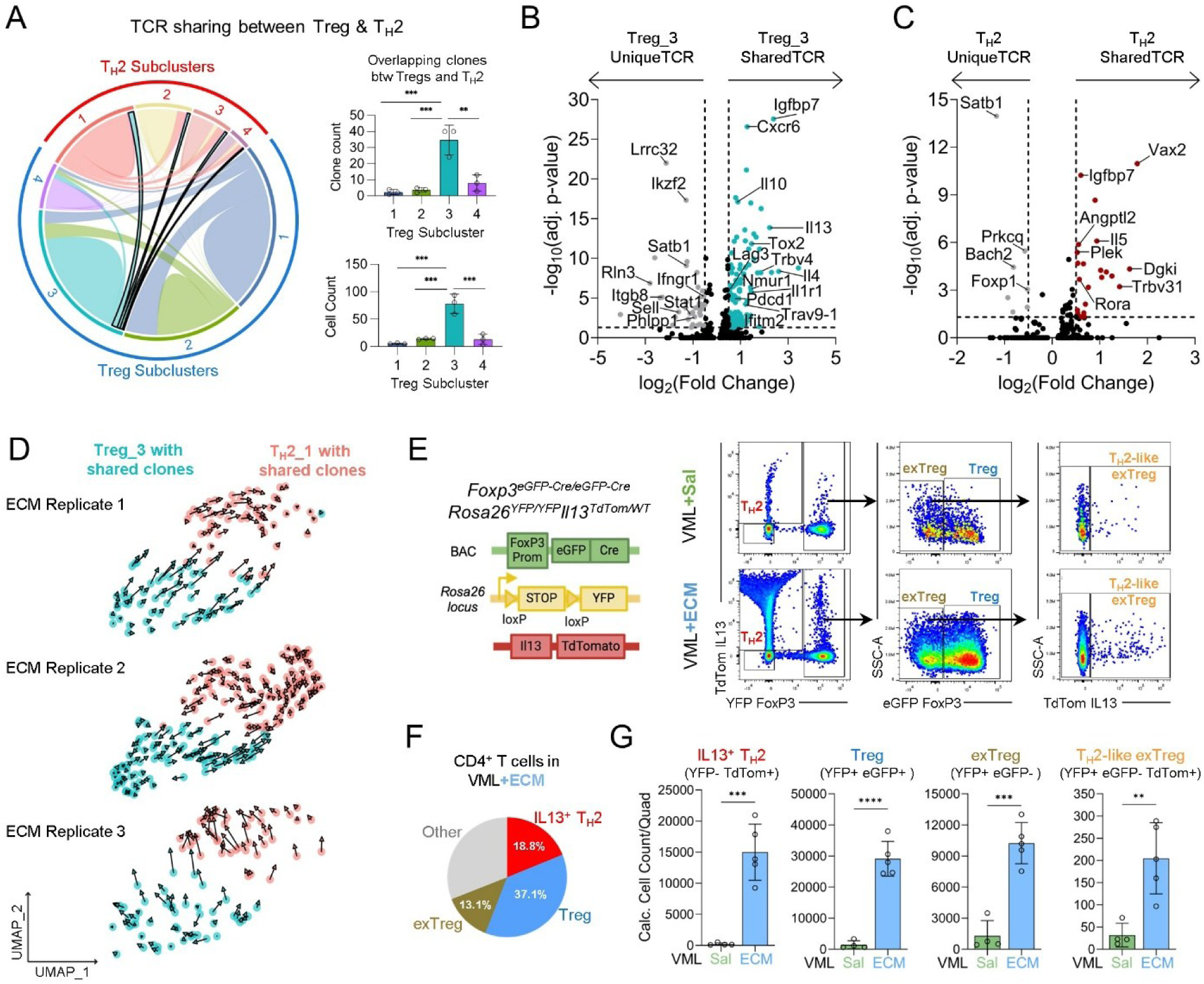
Treg plasticity and transition to T_H_2-like exTregs in ECM-treated wounds. Paired single-cell RNA/TCR sequencing enabled detection of abTCR sequences for cells in the different T cell clusters. Clones were strictly defined as cells that shared an identical abTCR based on CDR3 nucleotide sequences. **(A)** Chord plot of TCR sharing within Treg and T_H_2 sub-clusters in ECM-treated VML. Black outline highlights Treg_3 TCRs shared with T_H_2 sub-clusters. Quantification of unique shared clones (abTCR sequences) and total number of Tregs that share TCRs with T_H_2 cells by Treg sub-cluster. **(B)** Differential gene expression of Treg_3 cells that share TCR sequences with T_H_2 cells compared to Treg_3 cells that don’t or **(C)** of T_H_2 cells that share TCR sequences with Treg_3 cells compared to T_H_2 cells that don’t. **(D)** RNA velocity analysis of Treg_3 cells that share abTCR sequences with T_H_2_1 cells in ECM-treated VML by independent biological replicate. **(E)** Genetic construct of triple-reporter mouse strain and representative flow cytometry gating scheme to identify IL13-producing T_H_2 (CD4^+^YFP^-^TdTom^+^), Tregs (CD4^+^YFP^+^eGFP^+^), exTregs (CD4^+^eGFP^-^YFP^+^) and IL13-producing “T_H_2-like exTregs” (CD4^+^YFP^+^eGFP^-^TdTom^+^). **(F)** Composition of CD4^+^ T cells in ECM-treated VML injuries. **(G)** Quantification of IL13^+^ T_H_2, Treg, exTreg, and T_H_2-like exTreg cell counts/quad in saline-and ECM-treated VML injuries at 1-week. Data is presented as mean±SD and analyzed using one-way ANOVA with Tukey’s multiple comparisons test (A) or two-tailed unpaired student T test (G). NS p>0.05; * p<0.05; ** p<0.01; *** p<0.001; **** p<0.0001.

Next, we compared the transcriptional profile of Treg_3 cells that share TCRs with T_H_2 cells (Treg_3_sharedTCR) to Treg_3 cells that do not share TCRs with T_H_2 cells (Treg_3_uniqueTCR) (**Fig. 3B**). First, Treg_3_sharedTCR had similar *Nrp1,* but decreased *Ikzf2* (-1.29 fold change) expression relative to Treg_3_uniqueTCR (**Fig. S8E**). This greater decrease in relative *Ikzf2* expression by Treg_3_sharedTCR further supports their being pTregs and potential to become exTregs. Additionally, Treg_3_sharedTCR exhibited significantly greater expression of T_H_2-related genes (*Il4, Il13, Il10, Nmur1, Cxcr6, Ifitm2,* and *Ifitm3*)^32,44–46^ than Treg_3_uniqueTCR (**Fig. 3B**). Further, they had elevated expression of T cell activation genes (*Il1r1, Crtam, Pdcd1,* and *Il1r2*)^47,48^ and decreased expression of genes related to maintaining Treg phenotype (*Lrrc32, Itgb8, Satb1,* and *Phlpp1*).^49–52^ The Treg_3_sharedTCR had over-representation of the αTCR subunit *Trav9-1* and βTCR subunit *Trbv4* (**Fig. 3B**).

Next, we compared the transcriptional profile of T_H_2 cells that share TCRs with Treg_3 cells (T_H_2_sharedTCR) to T_H_2 cells that do not share TCRs with Treg_3 cells (T_H_2_uniqueTCR) (**Fig. 3C**). The T_H_2_sharedTCR cells demonstrated increased expression of the T_H_2-related genes *Il5* and *Angptl2*. Notably, T_H_2_sharedTCR cells also demonstrated increased expression of the Treg-associated genes *Rora* and *Dgki*.^46,53^ As a complementary analysis, RNA velocity was performed exclusively on cells in Treg_3 and T_H_2_1 with shared TCRs (**Fig. 3D**). The resulting trajectory suggested a transition from Treg_3 to T_H_2_1, which aligned with our prediction that Treg_3 cells may transition to a future T_H_2-like exTreg state.^54^ This putative Treg phenotype transition was consistently observed across ECM-treated biological replicates and using multiple dimensional reduction visualization techniques (**Fig. S7F**).

To study dynamics between Tregs and T_H_2 cells in ECM-treated injuries, we utilized a triple reporter strain that was originally constructed by crossing FoxP3 fate-mapping mice with IL13 TdTomato reporter mice (**Fig. 3E, S9A, S9B**).^55,56^ The resulting *Foxp3*^eGFP-Cre/eGFP-Cre^ *Rosa26^YFP/YFP^ IL13^TdTomato/WT^*(referred to as FoxP3^eGFP/YFP^IL13^TdTomato^) enabled us to simultaneously track Tregs (YFP^+^eGFP^+^, currently express FoxP3) and IL13-producing T_H_2 cells (YFP^-^TdTom^+^). Additionally, this strain allowed us to identify exTregs (YFP^+^eGFP^-^), which are cells that previously expressed FoxP3 but no longer do so. Using these mice, we validated that both IL13^+^ T_H_2 cells and FoxP3^+^ Tregs are significantly increased in 1-week ECM-treated wounds relative to saline-treated controls, both as a percentage of CD4^+^ T cells and absolute cell counts per quad (**Fig. 3F, 3G, S9C**). A sub-population of FoxP3^+^ Tregs that expressed the canonical T_H_2 cytokine IL13 also significantly increased with ECM wound treatment (**Fig. S9D**).

To validate the computationally putative Treg-to-T_H_2 transition, we first confirmed that exTregs were present in both saline-and ECM-treated wounds, albeit at significantly higher numbers with ECM treatment (**Fig. 3E-G, S9E**). In ECM-treated VML injuries, exTregs comprised 13.1±1.7% of CD4^+^ T cells, suggesting considerable potential for Treg plasticity (**Fig. 3F**). Next, we identified a “T_H_2-like exTreg” population that produced IL13 (YFP^+^eGFP^-^TdTom^+^). While only a small proportion of exTregs in the ECM-treated injuries produced IL13 (1.94±0.50%), this represented nearly an 8-fold increase in the number of T_H_2-like exTregs (**Fig. 3G, S9F**), suggesting that T_H_2-like exTregs contribute to the overall T_H_2 cell expansion observed in ECM-treated muscle wounds. Lastly, a significantly greater proportion of the two T_H_2-associated cells (IL13^+^ T_H_2 and IL13^+^ exTreg) expressed LAG3 compared to Tregs and IL13^-^ exTregs, potentially indicating that these T_H_2-associated cells may be more sensitive to ICI (**Fig. S9G**).

### Transcriptomic analysis predicts ICI responsiveness in mouse and human ECM scaffolds

Given the increased expression of immune checkpoint expression in ECM-treated wound environments, we explored whether ICI could enhance the ECM-induced pro-regenerative type 2 immune response. In our murine scRNA/TCRseq dataset, examining transcriptomic profiles of Treg_3 and T_H_2 clusters yielded a host of genes associated with ICI-responsiveness (**Fig. 4A**). Relative to other Tregs, the Treg_3 exhibited increased gene expression of human cancer ICI-responsiveness markers including *Podnl1* (glioma); *Lag3*, *Cxcr6*, and *Ccr5* (HCC: hepatocellular carcinoma and NSCLC: non-small cell lung cancer); *Cst7* (CRC: colorectal cancer); and *Il1r2* (pancreatic cancer^).57–62^ Additionally, Treg_3 had higher expression of *Angptl2*, *Entpd1,* and *Cxcr6*, which are markers of ICI-responsiveness in murine tumor models.^63–65^ The T_H_2 cells also upregulated markers of ICI-responsiveness relative to other T cells, including *Ctsd* and *Lpcat3* (melanoma); *Ramp1* and *Jup1* (gastric cancer); *Lag3* and *Elovl1* (HCC); *Il1r2* (pancreatic cancer); *Cst7* (CRC); *Fgl2* (bladder cancer); and *Cd74* (multiple tumor types).^66–73^ The T_H_2 cells also expressed markers of murine ICI-responsiveness included *Angptl2*, *Cxcr6*, and *Bhlhe40*.^74^ Notably, these subsets had increased expression of several ICI-responsive genes present in the ‘PredictIO’ gene set (*Lag3*, *Nkg7*, *Tbx21*, *Cst7, Gzmb, Cxcr6, Ikzf3, Prkch, Gbp5, Rab20, Rab11fip4,* and *Eaf1)*.^75^

**Fig. 4.**
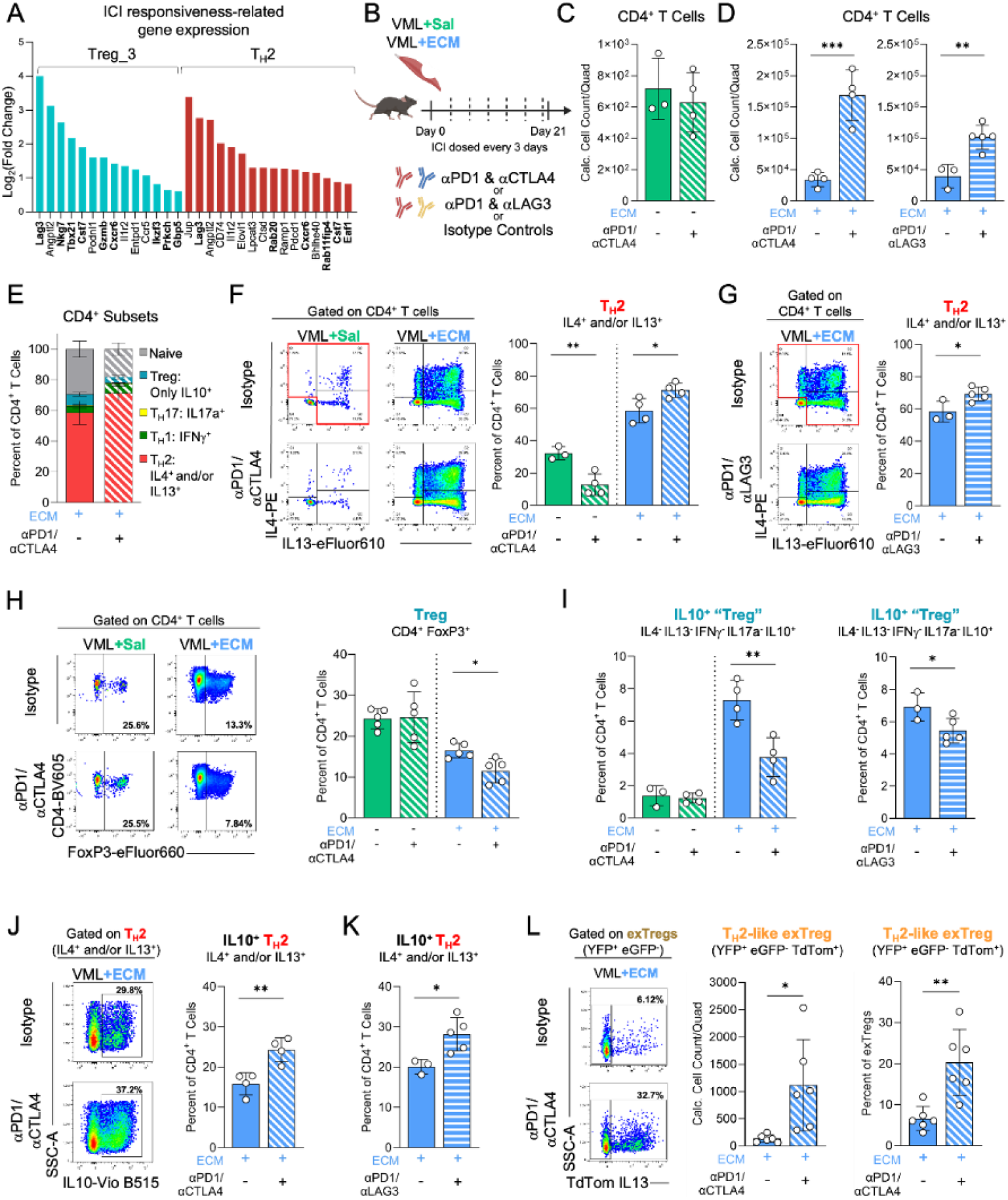
Immune checkpoint inhibition (ICI) expands T_H_2 and T_H_2-like exTregs in ECM-treated wounds. **(A)** Differential expression of genes associated with ICI responsiveness in Treg_3 and T_H_2 cells relative to all other Tregs or all other T cells, respectively. Bolded genes are those utilized by PredictIO.^74^ **(B)** Experiment schematic for 3-week ICI dosing regimen consisting of either combination aPD1/aCTLA4 (5mg/kg each) or aPD1/aLAG3 (5 mg/kg and 20 mg/kg, respectively) following saline-or ECM-treated VML. **(C)** Quantification of CD4^+^ T cell counts/quad in saline-or **(D)** ECM-treated VML following ICI. **(E)** Stacked bar plot comparing relative abundance of CD4^+^ T cell subsets (T_H_1, T_H_2, T_H_17, Treg, Naïve) identified based on cytokine expression profile in ECM-treated VML following ICI. **(F)** Representative flow plots and frequency of T_H_2 cells (CD4^+^, IL13^+^ and/or IL4^+^) in saline-or ECM-treated VML following aPD1/aCTLA4 or **(G)** aPD1/aLAG3. **(H)** Representative flow plots and frequency of Tregs (CD4^+^ FoxP3^+^) in saline-or ECM-treated VML following aPD1/aCTLA4. **(I)** Frequency of IL10^+^ “Tregs” (CD4^+^, IL4^-^ IL13^-^ IFNg^-^ IL17a^-^ IL10^+^) in saline-or ECM-treated VML following aPD1/aCTLA4 or aPD1/aLAG3. **(J)** Representative flow plots and frequency of IL10^+^ T_H_2 cells in ECM-treated VML following aPD1/aCTLA4 or **(K)** aPD1/aLAG3. **(L)** Representative flow plots, cell count/quad, and frequency of T_H_2-like exTregs (CD4^+^ YFP^+^ eGFP^-^ TdTom^+^) in ECM-treated VML following aPD1/aCTLA4. Data is presented as mean±SD and analyzed using two-tailed unpaired student T test (C, D, F-L). NS p>0.05; * p<0.05; ** p<0.01; *** p<0.001; **** p<0.0001.

To explore ICI-responsive gene signatures in our clinical ECM implant biopsies, we performed spatial transcriptomic analysis using the NanoString GeoMx Digital Spatial Profiler. We selected regions of interest (ROIs) based on H&E staining morphology as well as endothelial cell (CD31) and nuclear (SYTO13) markers. In total, 24 CD31^-^ segmented ECM ROIs were compared with 13 native adipose tissue ROIs. The ECM ROIs had significantly increased expressed of 46 genes, multiple of which were associated with ICI-responsiveness in human solid tumors including *ITGB2, FCER1G,* and *TPM4* (glioma); *SIRPA* (melanoma); *TFRC* (gastric cancer); *CD74* and *LY6E* (multiple tumor types) (**Fig. S10**).^70,76–82^ Additionally, several other increased genes are implicated in the regulation of immune checkpoint expression including *NR4A1, PKM,* and *CMTM6*.^83–85^ *CD74* expression was also increased by ECM-specific murine T_H_2 cells and is a known regulator of Treg and conventional CD4^+^ T cell activity.^86,87^ The only ‘PredictIO’ gene that was upregulated in this dataset was *B2M*.^74^ Collectively, the protein expression of immune checkpoints and ICI-related transcriptomic profiles suggests mouse and human ECM implant immune responses may be sensitive to and modulated by ICI.

### ICI potentiates T_H_2 immune response in ECM-treated wounds

To assess the effect of ICI on T cell responses to tissue injury and ECM implants, we administered combination anti-PD1/anti-CTLA4 (5mg/kg each, RMP1-14/9H10), combination anti-PD1/anti-LAG3 (5mg/kg and 20mg/kg respectively, RMP1-14/C9B7W), or isotype controls to wild-type C57BL/6 mice with saline-or ECM-treated VML injuries over a 3-week period (**Fig. 4B**). Flow cytometric analysis revealed that both combination ICI regimens significantly increased CD4^+^ T cells in ECM-treated wounds (absolute cell counts and percentage of CD3^+^ T cells), but did not affect CD4^+^ T cells in saline-treated wounds (**Fig. 4C, 4D, S11A**). Combination anti-PD1/anti-CTLA4, also increased the absolute cell counts of other lymphoid populations including CD8^+^ T cells, γδ T cells, NKT cells, NK cells, and B cells in ECM-treated wounds, but significantly decreased CD8^+^ T cell and NKT cell percentage out of CD3^+^ T cells (**Fig. S11B-I**).

Using an intracellular cytokine staining panel to characterize CD4^+^ T cell subsets, we assessed the impact of ICI on T_H_1 (IFNγ), T_H_2 (IL4 and IL13), T_H_17 (IL17a), and Tregs (exclusively IL10) following *ex vivo* stimulation (**Fig. 4E**). Both combination ICI regimens significantly expanded T_H_2 cells (IL4^+^ and/or IL13^+^) in ECM-treated wounds relative to isotype controls, both in terms of absolute cell counts and percentage of CD4^+^ T cells (**Fig. 4F, 4G, S12A**). The ICI regimens did not significantly affect the percentage of T_H_1 or T_H_17 cells in ECM-treated wounds, but increased the absolute cell count of T_H_1 cells (**Fig. 4E, S12C**). Interestingly, combination ICI decreased the T_H_2 compartment in saline-treated wounds, suggesting an ECM scaffold-specific response (**Fig. 4F, S12D**). The ICI-induced expansion of T_H_2 cells was contingent on the use of combination ICI therapy and not observed with anti-PD1, anti-CTLA4, or anti-LAG3 monotherapy regimens (**Fig. S13**). Intranuclear staining for the Treg transcription factor FoxP3 demonstrated that combination ICI significantly decreased Treg percentage of CD4^+^ T cells in ECM-treated wounds (**Fig. 4H**). Intracellular cytokine staining for canonical Treg cytokine (exclusively IL10^+^) corroborated these results, showing that while both combination ICI regimens increased the absolute cell count of IL10^+^ “Tregs”, they significantly decreased IL10^+^ “Tregs” percentage of CD4^+^ T cells in ECM-treated wounds (**Fig. 4I, S12B**).

### ICI expands IL10^+^ T_H_2 cells and drives Treg transition to T_H_2-like exTregs in ECM-treated wounds

Since we observed that combination ICI shifts Treg and T_H_2 proportions in opposing directions in ECM-treated wounds, we further explored the effects of combination ICI on Treg and T_H_2 cell dynamics. First, both combination ICI regimens stimulated expansion of a T_H_2 (IL4^+^ and/or IL13^+^) subset that also expressed the canonical Treg cytokine IL10 in ECM-treated injuries, both in terms of absolute cell count and percentage of CD4^+^ T cells (**Fig. 4J, 4K, S14A**). Conversely, the IL10^-^T_H_2 subset only increased in absolute cell counts and did not change as percentage of CD4 ^+^ T cells (**Fig. S14B, S14C**). Using the FoxP3^eGFP/YFP^IL13^TdTomato^ reporter mice, we found that combination anti-PD1/anti-CTLA4 decreased the Treg (YFP^+^eGFP^+^) percentage of CD4^+^ T cells, but increased the exTregs (YFP^+^eGFP^-^) percentage of YFP^+^ cells in ECM-treated wounds (**Fig. S14D-F**). Further, we observed that anti-PD1/anti-CTLA4 significantly increased the frequency and absolute cell count of T_H_2-like exTregs (YFP^+^eGFP^-^TdTom^+^) in ECM-treated wounds (**Fig. 4L**), demonstrating that ICI drives Treg transition to T_H_2-like exTregs. Collectively, we hypothesize that ECM treatment of VML injuries promotes ICI-responsive LAG3^+^ CD4^+^ T cells which, upon ICI treatment, preferentially expand towards a T_H_2 phenotype.

### ICI increases pro-regenerative myeloid cells, promotes muscle regeneration, and reduces fibrosis in ECM-treated injuries

Since we observed that combination anti-PD1/anti-CTLA4 stimulated a larger CD4^+^ T cell and T_H_2 response, we examined whether this ICI regimen also modulates ECM-enriched pro-regenerative myeloid cells after VML injury. ICI did not affect F4/80^+^ macrophage frequency or absolute count and did not affect the presence of canonical M1 (CD86^+^ CD206^-^) and M2 (CD86^-^ CD206^+^) macrophages in either saline-or ECM-treated wounds (**Fig. S15A-C**). Previously our group has characterized novel macrophage subsets associated with pro-regenerative biological and pro-fibrotic synthetic materials.^87^ The pro-regenerative ECM implants promote macrophage populations that we designated “R1” and “R2”, which are defined by their CD301b, CD9, CD206, and MHCII expression profiles (**Fig. 5A, S20**). ICI polarized macrophages towards the R2 phenotype while diminishing the presence of R1 macrophages in ECM-treated wounds (**Fig. 5A**). The percentages and absolute cell counts of fibrosis-associated macrophage subsets did not change with ICI treatment in either wound condition (**Fig. S15D, S15E**). Other myeloid cell types important for type 2 immune responses include eosinophils, basophils, and mast cells, which functionally modulate muscle injury healing.^88–90^ While eosinophils significantly increased as a proportion of CD45^+^ immune cells in ECM-treated wounds relative to saline-treated controls, they were not affected by ICI (**Fig. S15F**). Mast cells exhibited an increase in cell count but not as a percentage of CD45^+^ cells in response to injury treatment or ICI (**Fig. S15G**). Basophils, while a small percentage of the CD45^+^ immune cell population, doubled in response to ICI in ECM-treated wounds (**Fig. S15H**). Lastly, monocytes and neutrophils remained unaffected by ICI treatment in both wound conditions (**Fig. S15I, S15J**).

**Fig. 5.**
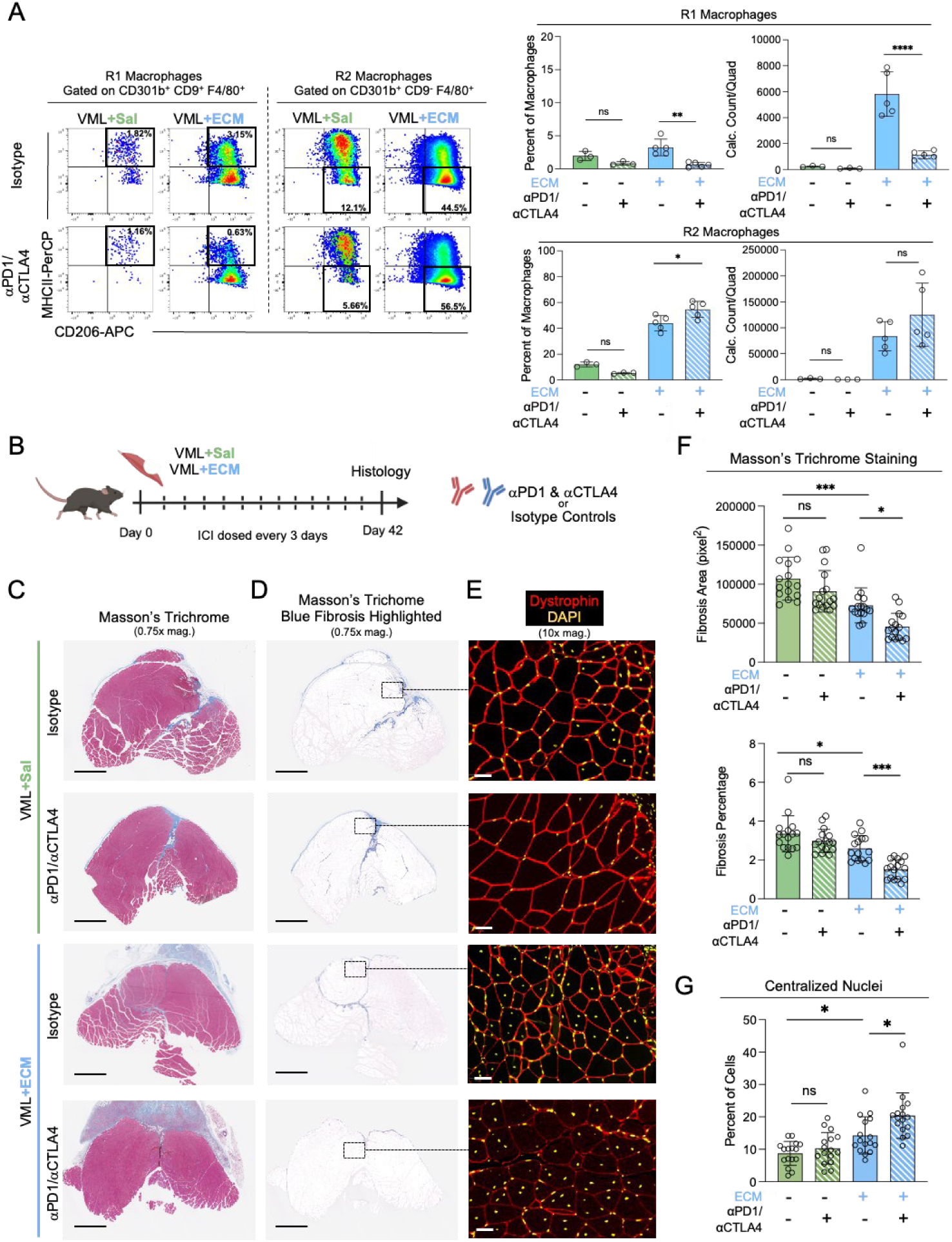
ICI expands R2 macrophages and promotes muscle regeneration in ECM-treated wounds. **(A)** Representative flow plots, frequency, and absolute counts of regenerative R1 (CD301b^+^ CD9^+^ MHCII^+^ CD206^+^) and R2 (CD301b^+^ CD9^-^ MHCII^-^ CD206^+^) macrophages (CD45^+^ CD11b^+^ SiglecF^-^ Ly6C^lo/mid^ F4/80^+^) in saline-or ECM-treated VML following aPD1/aCTLA4 at 3-weeks. **(B)** Experiment schematic for 6-week ICI dosing regimen consisting of aPD1/aCTLA4 following saline-or ECM-treated VML to histologically assess muscle healing and fibrosis. **(C)** Axial tissue sections of saline-or ECM-treated VML injured muscle following 6-weeks of aPD1/aCTLA4. Representative Masson’s Trichrome staining (scale bar: 1 mm) or **(D)** isolation of Blue-only fibrosis from Masson’s Trichome staining (scale bar: 1 mm) or **(E)** immunofluorescence staining of Dystrophin (outlines myocytes, red) and DAPI (nuclei, yellow) (scale bar: 50 μm). **(F)** Quantification of muscle fibrosis as pixel area and percentage of total tissue area based on Blue-only Masson’s Trichrome staining. **(G)** Quantification of myocytes with centrally located nuclei. Data is presented as mean±SD and analyzed using two-way ANOVA with Tukey’s multiple comparisons test (A, F, G). NS p>0.05; * p<0.05; ** p<0.01; *** p<0.001; **** p<0.0001.

Lastly, we investigated whether ICI impacts muscle fibrosis and regeneration following VML injury. We administered anti-PD1/anti-CTLA4 (5mg/kg each, RMP1-14/9H10) or isotype controls to wild-type C57BL/6 mice with saline-or ECM-treated VML injuries over an extended 6-week period prior to muscle harvest for histological analysis (**Fig. 5B**). First, Masson’s Trichrome staining used for visualization of tissue fibrosis showed more pervasive fibrosis in saline-treated injuries compared to ECM-treated injuries (**Fig. 5C, 5D**). Within the ECM-treated condition, ICI significantly decreased quantified fibrosis (surface area and percentage) relative to isotype controls (**Fig. 5F**). No effect of ICI was observed in saline-treated injury. To assess muscle regeneration, we performed dystrophin immunofluorescence staining and quantified myocytes with centralized nuclei (marked by DAPI staining) along the injured edge of the muscle tissue (**Fig. 5E**). ECM-treated muscle injuries had a significantly increased proportion of myocytes with centralized nuclei compared to saline-treated muscle injuries (**Fig. 5G**). ICI further increased the proportion of myocytes with centralized nuclei in ECM-treated wounds, suggesting synergy with the biomaterial scaffold to enhance muscle regeneration.

## Discussion

Our findings demonstrate a novel application of ICI to augment the pro-regenerative effects of biologic scaffolds for tissue repair and wound healing. Previous work focuses on modifying biomaterials to enhance their pro-regenerative effect via biomaterial functionalization, our results shift the focus to modulating the ECM-specific immune response using ICI to improve biomaterial-mediated muscle healing.^91,92^ While the results in this study coincide with earlier observations that T_H_2 cell-mediated immunity is beneficial for muscle regeneration, our work points to a novel relationship between Tregs and T_H_2 cells in the response to biomaterial implants.^93^ Additional studies are needed to fully understand the roles of different cell types responding to ECM implants, with exciting findings pointing outside of the T cell and macrophage compartments to B cells, dendritic cells, and mast cells.^94–96^ Here, we used ICI as a means to understand the role of immune checkpoint molecules in regulating ECM-driven immune responses and to demonstrate that systemic ICI treatment is capable of expanding pro-regenerative T_H_2 cells and macrophage subsets. As the fields of bioengineering and immunology increasingly intersect, there will likely be additional opportunities to leverage immune-modulating therapies for augmenting the effects of biomaterials in different disease contexts.

While the contribution of Tregs to ECM immune response is not surprising given their known importance in untreated muscle injury repair^23^, the observation that Tregs can switch phenotypes and become exTregs is novel in the context of wound healing and biomaterials. exTregs are derived from pTregs after exposure to T_H_1-, T_H_2-, or T_H_17-related cytokines and are well-documented in other settings like atherosclerosis, asthma and allergy, and autoimmunity.^41^ Here we show that a population of Tregs recruited to ECM-treated muscle injury expresses low levels of *Nrp1* and *Ikfz2,* consistent with murine pTregs.^27^ While pTregs have been shown to recognize self-antigens and subsequently contribute to autoimmunity after transitioning to exTreg-states, our model suggests that clonally expanded pTregs may contribute to the pro-regenerative immune response in their transition to T_H_2-like exTregs.^28^ Although we did not identify a specific αβTCR sequence enriched in the ECM condition across biological replicates, we did find that clone-sharing Tregs and T_H_2 cells were more likely to express the αTCR subunit *Trav9-1* and βTCR subunit *Trbv4*, suggesting a potential proclivity for these subunit alleles in recognizing a limited repertoire of ECM-derived epitopes. Like in other settings where Tregs become T_H_2-like exTregs, such as airway inflammation, the immune microenvironment of ECM-treated muscle injuries is rich in T_H_2-related cytokines.^41^ The presence of IL10^+^ T_H_2 cells, and their heightened responsiveness to ICI, begs the question of whether these T_H_2 cells are derived, at least in part, from T_H_2-like exTregs detected in our fate-mouse experiments. Future studies can confirm this as well as the secreted factors that modulate the Treg-exTreg transition in our model. Lastly, we report for the first time a role for ICI in the induction of exTregs. Our study identified high *Lag3* expression in our ECM-recruited pTreg population (Treg_3 subcluster), a finding consistent with prior work identifying a role for LAG3 in exerting the immunosuppressive effects of pTregs in graft-versus-host disease.^97^ In another study, pTreg-deficient mice exhibit reduced T cell exhaustion and enhanced T cell infiltration into tumors.^98^ Our results suggest a direct effect of ICI on pTregs and the induction of exTregs, implicating their potential role in anti-tumor immunity or in the development of immune-related adverse events (irAEs) which have not yet been explored.

ICI, which employs monoclonal antibodies to disrupt canonical regulation of T cell and antigen presenting cell activation, has revolutionized the treatment of many cancer types.^1^ With an expanding list of therapeutic options capable of inhibiting various immune checkpoint molecules, the use of ICI has spread beyond oncology and into other areas of medicine and pathophysiology. Prior to this study, the role of immune checkpoints in wound healing was poorly described. Clinically, there appears to be little evidence that perioperative ICI effects wound healing in various contexts including abdominal surgery, melanoma surgery, and Mohs surgery reconstruction.^99–101^ Similarly in our study, we found no effect of ICI on muscle wound healing in the absence of ECM implants. Despite the expression of immune checkpoints (PD1, CTLA4, and PDL1) in saline-treated muscle injury, ICI did not impact T cell and macrophage frequency or phenotype and had no effect on muscle fibrosis or regeneration. Conversely, ECM implants increased LAG3 expression in Tregs; LAG3, CTLA4, and TIGIT expression in conventional CD4^+^ T cells; and PDL2 expression in macrophages. These cells appeared to be ICI-responsive, displaying changes in cell count, frequency, and phenotype with ICI, ultimately leading to improved muscle healing. Human ECM samples similarly expressed LAG3 and PDL2, suggesting the human immune response to ECM-based implants could similarly be augmented by ICI. These differences in immune checkpoint expression and overall ICI-responsiveness may be explained by the potential xenogeneic and allogeneic epitopes present in ECM implants. Meanwhile, the self-antigens released during saline-treated muscle injury alone potentially fail to stimulate similar responses due to positive and negative TCR selection during T cell development. Nonetheless, the findings presented here point to a novel application of ICI and provide additional understanding of the immune response to biomaterial implants. While we delivered ICI systemically in these proof-of-concept studies, engineering strategies for localized and sustained ICI delivery at the ECM-treated injury site should be explored to minimize frequent high dosing and irAE risk. This therapeutic combination may be particularly beneficial in aged contexts, where biomaterial wound healing responses are impaired and immune checkpoint expression is elevated.^102^

In conclusion, this study demonstrates that ECM treatment of muscle injuries promotes LAG3 expression on Tregs and conventional CD4^+^ T cells, increases PDL2 expression on macrophages, encourages Treg transition to T_H_2-like exTregs, and heightens overall ICI-responsiveness. We find analogous markers of ICI-responsiveness, including expression of immune checkpoints LAG3 and PDL2, in clinical biopsies of human ECM implants. We demonstrate that ICI increases the type 2 immune response by stimulating T_H_2 expansion, in part by promoting T_H_2-like exTregs, and macrophage polarization towards a pro-regenerative phenotype, that ultimately augments muscle healing. Overall, these findings suggest a novel mechanism by which the pro-regenerative effects of biomaterial implants can be optimized for improved tissue regeneration using ICI.

### Limitations of the study

Although we found correlating results between the murine and human immune responses to ECM-based biomaterials, additional studies will be needed to further verify our murine findings in humans. An inherent limitation to our analysis is the discrepancy between the timing of tissue analysis in our murine studies (1-3 weeks) versus human studies (3-9 months). The timing of human biopsies was due to a previously determined clinical trial protocol and samples were processed in a retrospective manner. Future studies could consider earlier biopsy timepoints and process samples prospectively to accommodate high-throughput methodologies like scRNAseq and spectral flow cytometry. The signal of LAG3 and PDL2 expression may be brighter at earlier timepoints as they are in mice, and future studies could also explore the role of exTregs in the response to biomaterials. Lastly, the markers for tTregs and pTregs (*Nrp1* and *Ikzf2*) are not reliable in humans, so were not explored in this study.^27,28^ As such, markers of tTregs and pTregs should be developed if Treg origin predicts the likelihood of becoming an exTregs.

## Acknowledgements

We would like to thank the Johns Hopkins University (JHU) animal facilities and caretakers, JHU SKCCC High Parameter Flow core facility for performing FACS, JHU Transcriptomics & Deep Sequencing core facility for performing single-cell RNA sequencing, UPMC Hillman Cancer Center for performing the spatial transcriptomics using NanoString Digital Spatial Profiler, JHU Oncology Tissue Services SKCCC core facility for histology sectioning, Dawn Newcomb for sharing breeder pairs for the FoxP3^eGFP/YFP^IL13^TdTomato^ reporter strains, Jessica Gucwa from Cytek for flow cytometry input, and ACell Integra LifeSciences for providing ECM materials. We thank the Departments of Otolaryngology-Head and Neck Surgery and Plastic and Reconstructive Surgery at Johns Hopkins Hospital for accommodating our clinical trial patients in their clinics. Finally, we would also like to express our sincerest gratitude for the patients who participated in the clinical trial (NCT03544632) through which tissue biopsies could not otherwise be obtained.

## Funding

NIH Pioneer Award DP1AR076959 (J.H.E.)

NIH T32 Postdoctoral Research Training Program 3T32DC000027-30S1 (J.G.) NSF Graduate Research Fellowship Program DGE1746891(A.R.) K99AG081564 (J.C.M)

Mark Foundation for Cancer Research (K.N.S., D.M.P.) Bloomberg∼Kimmel Institute for Cancer Immunotherapy (K.N.S., D.M.P)

## Author Contributions

J.G., A.R., D.M.P., and J.H.E. conceptualized and designed the studies.

J.G., A.R., J.C.M., and D.R.M. performed experiments and interpreted results. C.D., N.R., E.F.G.G., M.B., and K.S. assisted with experimental work.

F.H.Y. performed computational analyses and interpreted results.

K.K., C.C., E.J.F., K.N.S., and H.J. assisted with and advised computational analysis.

A.N.P., A.B., and E.J.F. performed and advised digital spatial profiling of human clinical samples. J.G., C.M.C., D.C., A.T.H., and P.B. supported or conducted clinical trial workflow and biopsies.

R.A. performed histology staining of human clinical samples and interpreted results. J.G., A.R., and J.H.E. wrote the manuscript.

D.M.P. and J.H.E. supervised the study.

## Competing Interests

C.C. is the owner of C M Cherry Consulting. E.J.F. was on the scientific advisory board of Resistance Bio and a consultant for Mestag Therapeutics. P.B. is chief medical officer for Aegeria Inc. A.T.H. is a consultant for Airiver, Inc. K.N.S. holds founders’ equity in Clasp Therapeutics.

D.M.P. is a consultant at Aduro Biotech, Amgen, Astra Zeneca, Bayer, Compugen, DNAtrix, Dynavax Technologies Corporation, Ervaxx, FLX Bio, Immunomic, Janssen, Merck, and Rock Springs Capital. D.M.P. holds equity in Aduro Biotech, DNAtrix, Ervaxx, Five Prime therapeutics, Immunomic, Potenza, Trieza Therapeutics. D.M.P. is a member of the scientific advisory board for Bristol Myers Squibb, Camden Nexus II, Five Prime Therapeutics, and WindMil. D.M.P. is a member of the board of directors in Dracen Pharmaceuticals. J.H.E. holds equity in Unity Biotechnology and Aegeria Soft Tissue and is a consultant for Tessara.

## Data and Materials Availability

Raw FASTQ files and aligned count files for both RNA and TCR sequences will be made available for download through NCBI GEO under the accession number **GSE308460**. Processed objects will be available upon reasonable request. Scripts used for data analysis and visualization in this work were written in R and Python. Scripts for all analysis of scRNA/TCRseq are publicly accessible in the GitHub repository hosted at https://github.com/Elisseeff-Lab/T-Cells-in-ECM-Modulated-Wounds. All other data are available in the main text or the supplementary materials.

## Supplementary Materials

Materials and Methods

Figs. S1 to S20

Tables S1 to S8

## Materials and Methods

### Mice

All murine procedures were approved by the Johns Hopkins University Institutional Animal Care and Use Committee (protocols MO21M80/MO24M66). Wild-type C57BL/6J mice were obtained directly from Jackson Laboratory (stock #00064). The triple reporter FoxP3^eGFP/YFP^IL13^TdTomato^ strain was generated by crossing the parent strains *Foxp3*^eGFP-Cre/eGFP-Cre^*Rosa26^YFP/YFP^*with *Foxp3*^eGFP-Cre/eGFP-Cre^*Rosa26^YFP/YFP^Il13^TdTomato/TdTomato^*, both gifted by Dawn Newcomb (originally from Talal Chatila and Andrew McKenzie).^55,56,103^ Mice were genotyped to confirm strain identities. Wild-type C57BL/6J mice were maintained as Helicobacter negative, while FoxP3^eGFP/YFP^IL13^TdTomato^ reporter mice and their parent strains were received as Helicobacter positive. Only female mice at 6-7 weeks of age were used.

### Volumetric Muscle Loss (VML) Injury Model

Mice were randomized prior to start of experiments between uninjured, untreated VML injury (saline) or biomaterial-treated VML injury groups. Mice were anesthetized and surgical sites thoroughly disinfected. Pain management was provided with 5 mg/kg subcutaneous injections of carprofen (Rimadyl, Zoetis). Bilateral VML injury was performed as a survival surgery where a partial thickness defect is made in the quadriceps femoris muscles. A longitudinal incision is made through skin and fascia to expose 10mm of the muscle. A critical-sized portion of muscle (approximately 3 mm x 4 mm x 4 mm) was resected from each hindlimb using sterilized dissection scissors. The wound beds were each treated with either 75 μL of 1x sterile Dulbecco’s Phosphate Buffered Saline (dPBS) or extracellular matrix (ECM) implant material (200 mg/mL in dPBS, see below). Immediately after treatment, the skin was closed with suture. Mice were closely monitored for the duration of the experiment (3 days, 1 week, 3 weeks, or 6 weeks) before euthanasia.

### Extracellular Matrix (ECM) Scaffolds

ECM biomaterial utilized in this study was primarily porcine-derived urinary bladder matrix (UBM) (ACell, Integra LifeSciences), which was reconstituted at 200 mg/mL in sterile dPBS prior to applying to VML muscle injury sites. For allogenic ECM source, mouse collagen type I/III from tail tendon (Bio Rad) was reconstituted at 200 mg/mL in sterile dPBS.

### Immune Checkpoint Inhibitor (ICI) Treatment

Mice were randomized between isotype control or ICI groups prior to treatment. Mice were treated with ICI or matched isotype controls via intraperitoneal injection every 3 days starting on day 3 post VML injury. The ICI regimens trialed in this study included anti-PD1 (RMP1-14, BioXCell), anti-CTLA4 (9H10, BioXCell), and anti-LAG3 (C9B7W, Leinco Technologies). Appropriate isotypes control antibodies used in this study included Rat IgG2a Isotype (2A3, BioXCell), Syrian Hamster Isotype (polyclonal, BioXCell), and Rat IgG1 Isotype (R1379, Leinco Technologies), respectively. Anti-PD1 and anti-CTLA4 were each dosed at 5 mg/kg. The anti-LAG3 dose varied (5 mg/kg, 10 mg/kg, 20 mg/kg), but is clearly indicated in figure labels and captions. For the combination anti-PD1/anti-LAG3 experiment presented in **Figure 4**, the first ICI dose at day 3 post injury was only anti-PD1, while all subsequent doses were combination anti-PD1/anti-LAG3 (5 mg/kg and 20 mg/kg, respectively).

### Quadriceps Muscle Tissue Processing for Flow Cytometry

Quadriceps muscles and any remaining ECM scaffolds were harvested at pre-defined endpoints (3 days, 1 week, or 3 weeks after injury), with bilateral muscle pairs from the same mouse pooled together. The muscle was mechanically diced and digested at 37°C with 1.67 Wünsch U/mL (5 mg/mL) of Liberase TL (Roche Diagnostics, Sigma Aldrich) and 0.2 mg/mL DNase I (Roche Diagnostics, Sigma Aldrich) in RPMI-1640 media supplemented with L-Glutamine and 15 mM HEPES (Gibco) for 45 minutes. The digested tissue was passed through a 70 μm cell strainer (ThermoFisher Scientific) and washed twice with 1x dPBS. Samples being processed for exclusively T cell profiling (**Suppl Tables 3-7**) were further processed for leukocyte enrichment using a Percoll (GE Healthcare) density gradient (80%, 40%, and 20% layers) and centrifugation (2100 g for 30 minutes at room temperature, lowest acceleration and no breaks).

### Flow Cytometry Staining

For samples undergoing intracellular cytokine staining (**Suppl Tables 4,5**), cells were first counted and plated at 2 million cells/well, followed by *ex vivo* stimulation with Cell Stimulation Cocktail Plus Protein Transport Inhibitors (eBioscience) in complete cell culture media (RPMI-1640 supplemented with 10% FBS, 15 mM HEPES, and 5 mM Sodium Pyruvate) for 4 hours at 37°C in a cell culture incubator. All samples were plated in 96-well U-bottom plate, followed by viability staining for 30 minutes. Samples were washed with FACS buffer (dPBS with 1% w/v BSA) prior to surface staining (**Suppl Tables 1-7**) in the presence of TruStain Fcx (1:20) (BioLegend), Monocyte Blocker (1:50) (BioLegend), and Super Bright Complete staining buffer (1:50) (eBioscience) for 45 minutes on ice in the dark, followed by washed with FACS buffer. Samples from triple reporter mice (**Suppl Table 6**) or for FACS were acquired fresh, whereas other samples with only surface marker staining (**Suppl Table 1,2**) were fixed with FluoroFix buffer (BioLegend) for 15 minutes. Samples with intracellular cytokine staining were fixed and permeabilized using Cytofix/Cytoperm kit (BD Biosciences) (**Suppl Table 5**) or Cyto-Fast Fix/Perm buffer set (BioLegend) (**Suppl Table 4**) as per manufacturer protocol. Samples with transcription factor staining (**Suppl Table 3**) were fixed and permeabilized using True-Nuclear Transcription Factor buffer set (BioLegend) as per manufacturer protocol. Flow cytometry was performed using either a 4-laser Attune NxT Flow Cytometer (ThermoFisher Scientific) or 4-laser spectral Aurora (Cytek). For experiments using the triple reporter FoxP3^eGFP/YFP^IL13^TdTomato^, the voltage for blue (detector channels 4-14) and all yellow/green (detector channels 1-10) was decreased by 60% to get the bright TdTomato signal on-scale.

### Flow Cytometry Analysis

Spectral unmixing of data collected on the Cytek Aurora was performed using SpectroFlo software, with tissue autofluorescence signatures extracted from unstained samples. Flow cytometry datasets were analyzed using FlowJo software (v10.7-10.8, Tree Star), with manual gate placement based on fluorophores-minus-one (FMO) or isotype (for intracellular cytokines) controls. All reported populations are from events on-scale, singlets (using diagonal gating of FSC-H vs. FSC-A), live (negative for dead cell staining), CD45^+^ events. For intracellular cytokine staining studies, only samples within a pre-determined total cell count cut-off (< 1 million total collected cells) were included in downstream analyses to minimize technical artifact of under-stimulation or-staining. Gating schemes provided in **Figures S16-S20**.

### NanoString nCounter Gene Expression Analysis

Transcript profiling was performed to compare saline-treated (injury only controls) and ECM-treated muscles at 1-week post VML injury. Whole quadriceps muscle was harvested and placed in RNALater for at least 24 hours at 4°C and then transferred into TRIzol reagent (Thermo Fisher Scientific) and flash frozen for storage. Samples were then thawed and homogenized using a Bead Ruptor 12 (OMNI International) with 2.8 mm ceramic beads (OMNI International). RNA was isolated using TRIzol reagent and chloroform extraction. RNA was purified using the RNeasy Plus kits (Qiagen) and gDNA eliminator columns. RNA was quantified using a Qubit RNA fluorometric assay high-sensitivity kit (Thermo Fisher Scientific).

For isolated T cells, samples were processed and stained for flow cytometry as described above (**Suppl Table 8**). Fluorescence activated cell sorting (FACS) was performed using a 4-laser FACSAria Fusion Cell Sorter (BD) at the JHU SKCCC High Parameter Flow Core to isolate T cells (Singlets Live CD45^+^ CD11b^-^CD3^+^). T cells were sorted directly into RLT buffer containing 2-mercaptoethanol for cell lysis. RNA was purified using an RNeasy Plus Micro kit (Qiagen). RNA was quantified using a Qubit RNA fluorometric assay high-sensitivity kit (Thermo Fisher Scientific).

For NanoString, 100 ng of RNA per sample (3 replicates per condition) was added to the barcoded probe set for the nCounter Mouse PanCancer Immune Panel (NanoString Technologies). Samples were left to hybridize for 20 hours at 65°C. Samples were processed by the nCounter Digital Analyzer and NanoString Prep Station under high-sensitivity mode (NanoString Technologies). Data was analyzed using nSolver Software (v4.0).

### Single-Cell RNA/TCR Sequencing

Sample Acquisition: scRNA/TCRseq was performed on sorted CD3⁺ T cells from saline-treated (injury only controls) and ECM-treated muscles at 1-week post VML injury in wild-type C57BL/6J mice. For the saline-treated group (n = 14), cells from each independent biological sample were labeled with TotalSeq C oligo-tagged hashing antibodies (BioLegend) prior to pooling (4-5 samples together) for sorting and sequencing. For the ECM-treated group (n = 3), each independent biological sample was also labeled hashing antibodies, but due to high T cell yield were sorted separately and sequenced on individual lanes. Tissue samples were harvested, processed (including Percoll density gradient), and stained (viability and surface markers) as described above. For hashing, TotalSeq-C antibodies (**Suppl Table 7**) were added during the final 30 minutes of the surface staining protocol. FACS was performed using a 4-laser FACSAria Fusion Cell Sorter (BD) at the JHU SKCCC High Parameter Flow Core to isolate T cells (Singlets Live CD45^+^ CD3^+^ SCC^lo^). T cells were sorted into dPBS with 1% BSA, then centrifuged and re-suspended in dPBS with 0.01% BSA. Cells were counted using the Countess Automated Cell Counter (Thermo Fisher Scientific) and processed for 5′ paired scRNA /TCRseq with feature barcoding at the Johns Hopkins Transcriptomics & Deep Sequencing Core. Libraries for gene expression, surface protein feature barcodes, and paired αβ TCR sequences were prepared using the Chromium Next GEM Single Cell 5′ Library & Gel Bead Kit v1.1 (R2-only HT configuration) (10x Genomics).

Data Pre-Processing: FASTQ files were processed using Cell Ranger (10x Genomics, v7.2.0) with default parameters. Gene expression reads were aligned to 10x Genomics mouse reference transcriptome mm10-2020-A, which is based on GRCm38 genome assembly and GENCODE vM23 annotation^104^. To demultiplex the hashed CD3⁺ T cell samples, the Cell Ranger multi pipeline was used with hashtag oligo (HTO) barcode sequences to assign individual cells to their respective samples. Following demultiplexing, bamtofastq was used to generate sample-specific FASTQ files. For TCR analysis, the Cell Ranger vdj function was applied to align and assemble paired αβ TCR sequences from the corresponding V(D)J libraries.

Quality Control: High-quality cells were retained by applying the following thresholds: UMI count > 500, feature count > 400, and mitochondrial gene content < 3.5%. These criteria were used to exclude low-quality cells and potential empty droplets. Additionally, only genes expressed in at least 0.1% of cells were retained for downstream analysis to minimize noise from rare transcripts.

Data Processing: Seurat (v5.0.3) was used for normalization, identification of highly variable genes, scaling (including regressing out the percentage of mitochondrial genes, feature count and total UMI count), and principal component analysis (PCA)^105–107^. The number of principal components used for downstream analyses was guided by inspection of an elbow plot.

Dimensionality reduction was performed using Uniform Manifold Approximation and Projection (UMAP)^108^.

Clustering: Clustering was conducted using the Louvain algorithm for graph-based community detection^109^. Cluster resolution parameters were selected based on multiple criteria: silhouette scores, canonical marker gene expression, cluster stability assessed using Clustree (v0.5.1), and differential expression analysis performed using the Wilcoxon rank-sum test implemented in Presto (v1.0.0)^110,111^ testing genes expressed in more than 10% of cells in either comparison group. Cluster differential expression analysis was performed by comparing each cluster against all remaining cells. We manually removed the myeloid–T cell doublets and low-quality clusters after initial clustering. Seven major T cell clusters were manually annotated based on known marker genes (**Figure S4**).

Subclustering: We applied the same clustering pipeline described above to perform subclustering of select T cell populations. Within the “Mixed” cluster, subclustering revealed that subcluster 0 consisted of T_H_17 cells, while subclusters 1, 5, and 6 represented T_H_1 cells. We separated these T_H_17 and T_H_1 cells from the remaining cells in the “ Mixed” cluster, which comprised a heterogeneous mix of CD4⁺, CD8⁺, and γδ T cells (**Figure S6A**). Subclustering of the “T_H_2” cell cluster identified subclusters 0, 1, 4, and 6 as true T_H_2 cells based on canonical type 2 marker genes, which we manually separated from the remaining irrelevant populations (**Figure S6B**). Subclustering of the “Treg” cluster resulted in four distinct Treg subclusters (**Figure S7**). Resulting sub-clusters from T_H_2 and Treg were combined for downstream analysis and visualization.

Differential Expression Analysis: Differential expression analysis between any two groups of interest was performed using the Wilcoxon rank-sum test, testing genes expressed in more than 10% of cells in either comparison group. Genes with an adjusted p-value <0.05 and |log_2_(fold change)| >0.5 were considered significantly differentially expressed.

Gene Set Enrichment Analysis (GSEA): Ranked GSEA was performed using the fgsea package (v1.31.2) with 10,000 permutations based on Wilcoxon rank-sum test results^112^. Genes were ranked by the statistic:-log₁₀(adjusted p-value) × sign(log fold change). Gene sets were obtained from the KEGG and Hallmark collections within MSigDB^113,114^. Pathways with an adjusted p-value <0.05 were considered significantly enriched.

TCRseq Analysis: Paired scTCRseq data were integrated with gene expression data using scRepertoire (v1.12.0)^115^. TCR clones were defined based on cells with matching paired α and β chains, identified by their V(D)J gene usage and CDR3 nucleotide sequences. Clonal expansion was quantified per animal, enabling sample-specific assessments of T cell clonality.

Each unique combination of α (TRA) and β (TRB) chains, defined by V(D)J gene usage and CDR3 nucleotide sequences, was considered a distinct TCR clone. Complete TCR clones were identified based on cells sharing the same TRA–TRB pair. If only one chain (α or β) was recovered, these partial sequences were grouped only with cells that had the same available chain matched in V(D)J usage and CDR3 nucleotide sequence. Cells with both chains missing were excluded. Clones present in two or more cells were considered clonal, and the number of cells per clone was used to measure the extent of clonal expansion. For clonal analysis, we focused on T_H_2 and Treg cells from the ECM-treated condition. To categorize levels of expansion, we stratified clone sizes using the 75th, 90th, 99th, and 99.99th percentiles of clone sizes. These thresholds were adjusted within scRepertoire to fit the distribution of clone sizes. Based on this distribution, clones were classified as hyperexpanded (more than 40 cells), large (10 to 40 cells), medium (3 to 9 cells), small (2 cells), or single (1 cell) (**Figure S8B**).

To explore clonal relationships between T_H_2 and Treg subclusters, we counted both the number of shared clones and the number of contributing cells between each T_H_2 subcluster–Treg subcluster pair. These overlaps were visualized using chord diagrams generated with the circlize package and heatmaps created with the pheatmap package (**Figure 3A, S8C**)^116,117^. We also quantified clones shared between the T_H_2 cluster and each of the four Treg subclusters in ECM-treated animals and presented these results as bar plots (**Figure 3A**). In addition, we quantified clones shared between each of the four Treg subclusters and the T_H_1, T_H_2, and T_H_17 clusters and presented these results as bar plots (**Figure S8D**).

To compare transcriptomic profiles associated with TCR sharing, we carried out differential gene expression analysis using the Wilcoxon rank-sum test. Specifically, we compared Treg_3 cells that share TCR sequences with T_H_2 cells to Treg_3 cells that do not, and similarly, T_H_2 cells that share TCRs with Treg_3 cells to those that do not (**Figure 3B, 3C, S8E**).

RNA Velocity: Spliced and unspliced transcript matrices were generated for each ECM-treated sample using the Velocyto command-line tool.^54^ To infer transcriptional dynamics between T_H_2 and Treg_3 cells with shared TCR clonotypes, we extracted the relevant cell barcodes, along with their UMAP, t-SNE, PCA embeddings, and cluster annotations, in R and imported them into Python. For each sample, loom files were read and filtered in Python to retain only cells matching the extracted barcodes. RNA velocity was estimated using the dynamical model implemented in scVelo^118^. Velocity fields were visualized by projecting RNA velocities onto PCA, t-SNE, and UMAP embeddings using scVelo (**Figure 3D, S8F**).

ICI Responsiveness Related Gene Expression: Assessment of ICI responsiveness was determined by performing differential gene expression analysis between Treg_3 cells and the other Treg subclusters (Treg_1, Treg_2, and Treg_4) and between T_H_2 cell and all other T cells in the dataset. Differentially expressed genes (adjusted p-value <0.05) were manually reviewed and any with documented importance in predicting responsiveness to ICI in human or murine settings are highlighted.

Public CD45^+^ Enriched scRNAseq Dataset Analysis: A previously published dataset utilizing a Drop-Seq scRNAseq pipeline was utilized to determine *Pdcd1lg1* (*Cd274*) and *Pdcd1lg2* expression in macrophage subsets of uninjured, saline-treated VML injured, and ECM-treated VML injured muscles at 1-week.^21^ Differential gene expression was performed using the Wilcoxon rank-sum test.

### Quadriceps Muscle Histopathology

Quadriceps muscles were harvested at 6 weeks post VML injury and fixed in 10% neutral buffered formalin for 48 hours followed by dehydration in increasing concentrations of EtOH. Tissues were transferred to xylene prior to embedding in paraffin. Tissues were sectioned as 6 μm slices and stained for either Masson’s Trichrome or immunofluorescence. Dystrophin (clone EPR21189, Abcam) was stained at 1:1000 dilution using tyramide signal amplification method with Opal-650 (Akoya). After blocking with bovine serum albumin, the primary antibody was incubated at room temperature for 30 minutes, followed by 30 minutes of incubation with horseradish peroxidase polymer-conjugated secondary antibody, and 10 minutes with Opal-650. Slides were counterstained with 4′,6-diamidino-2-phenylindole (DAPI) for 5 minutes prior to mounting with DAKO mounting medium. Image acquisition was obtained using a Zeiss Axio Imager A2 and Zeiss AxioVision software (v4.2). Masson’s Trichrome was analyzed using ImageJ software. Briefly, intramuscular collagen deposition was quantified by creating a mask excluding all extra-muscular collagen and ECM implants. Collagen staining was then selected by excluding all color besides blue tones. The raw pixel count and percentage of muscle area occupied by blue collagen was quantified. Centralized nuclei were quantified using the QuPath software (0.51-x64). Regions of interest (ROIs) were created on each muscle specimen approximating the superficial 500 μm of muscle tissue. Total myocytes and myocytes with centralized nuclei were counted in each ROI.

### Phase 2 Clinical Trial Tissue Sample Collection

Human acellular adipose tissue (AAT) implant biopsies were previously obtained during a phase II, dose-escalation, open-label study (NCT03544632) in which patients with soft tissue defects received 1-2 injections of AAT material. This study was conducted at Johns Hopkins Hospital with IRB and HRPO approval (IRB00155003 and HRPO Log No. Eo14221a). Briefly, patients were enrolled in a 12-month, prospective, dose-escalation study to assess safety and efficacy of AAT injections ranging from 5-40 cc of AAT material. Patients underwent ultrasound-guided core needle biopsy anytime between 3-9 months after injection using a 14-gauge Max-Core disposable core-needle biopsy device and an Esoate MyLab50 XVision ultrasound machine. Samples for fixed in 10% formalin and embedded in paraffin prior to staining. 4 μm slices were sectioned on Leica Bond Plus and dried overnight at 37°C. Samples were stored in vacuum-sealed container at 4°C until further processing.

### Human Tissue Immunohistochemistry

Immunohistochemistry was performed on 7 patient biopsy samples. Automated LAG3 staining was performed on the Leica Bond RX (Leica Biosystems). Slides were baked and dewaxed online followed by antigen retrieval. Endogenous peroxidase was blocked then anti-LAG3 clone 17B4 (LS-C344932-100, LSBio, Shirley, MA, RRID: AB_3076336) was applied for 60 min at a concentration of 0.025 µg/mL at room temperature. Detection was performed by combining the PowerVision (Leica) and Tyramide Signal Amplification (Akoya) systems. Slides were counterstained, dried, and coverslipped using Ecomount (Biocare Medical). Automated PD-L2 staining was performed on the Leica Bond RX (Leica Biosystems). Slides were baked and dewaxed online followed by antigen retrieval. Endogenous peroxidase was blocked then anti-PD-L2 clone D7U8C (82723S, Cell Signaling Technologies, RRID: AB_2162081) was applied for 180 min at a concentration of 0.158 µg/mL at room temperature. Detection was performed by combining the PowerVision (Leica) and Tyramide Signal Amplification (Akoya) systems. Slides were counterstained, dried, and coverslipped using Ecomount (Biocare Medical).

### Digital Spatial RNA Profiling

The GEOMx Digital Spatial Profiler (RRID:SCR_021660) methodology was utilized to process samples.^119^ AAT samples were initially stained with CD45, CD31, CD68, and CYTO13 and RNA detection probes with UV-photocleavable oligonucleotides using standard immunohistochemistry protocols. ROIs were identified based on hematoxylin and eosin morphology staining. Two ROI types were identified: native adipose tissue and acellular adipose tissue implant. Next ROI segmentation was performed based on CD31^+^ staining, and samples were prepped for the GEOMx Digital Spatial Profiler. 192 circular ROIs were collected by photocleaving the bound oligonucleotides and indexing them based on ROI. The 1800+ gene probe set, Cancer Transcriptome Atlas (NanoString), was utilized and Next Generation Sequencing (Illumina) was used for profiling gene expression. This analysis was performed by the Cytometry Facility at the UPMC Hillman Cancer Center. Data analysis was performed in R and GraphPad Prism (v9.2). Biological probe quality control parameters were applied based on the manufacturer’s protocols. Probes were filtered from the data if the ratio of the geomean probe in all segments was less than or equal to 0.1 or if probes fail Grubbs outlier test where probes should not be an outlier in greater than or equal to 20% of the segments. The limit of quantification filtration was employed for genes and segments, and data were normalized using the variance stabilizing transformation (vst) function from DESeq2.^120^ Differential gene expression analysis was carried out using a linear mixed model.^121^ Patient identity was set as a random effect, and significance thresholds of p_adj_ < 0.05 and 0.58 ≤ Log Fold Change or ≤-0.58 were applied to determine differentially expressed genes. Spatial cell deconvolution was performed using the SpatialDecon R package^122^ with the ImmuneTumor_safeTME normalized cell matrix (NanoString).

### Statistics

Statistical analyses were performed using GraphPad Prism (v9.2-10.5). All data is presented at mean±SD and analyzed using unpaired two-tailed t-test, one-way ANOVA, or two-way ANOVA with Tukey’s multiple comparison’s test (clearly specified in figure captions). Statistical significance was designated at p < 0.05. NanoString nCounter results were analyzed using the nSolver Advanced Analysis Software (v4.0.70, NanoString Technologies). For NanoString analysis, False Discovery Rate-adjusted p-values were determined for each gene by applying the Benjamini-Yekutieli method.

## Supplementary Figures

**Fig S1.**
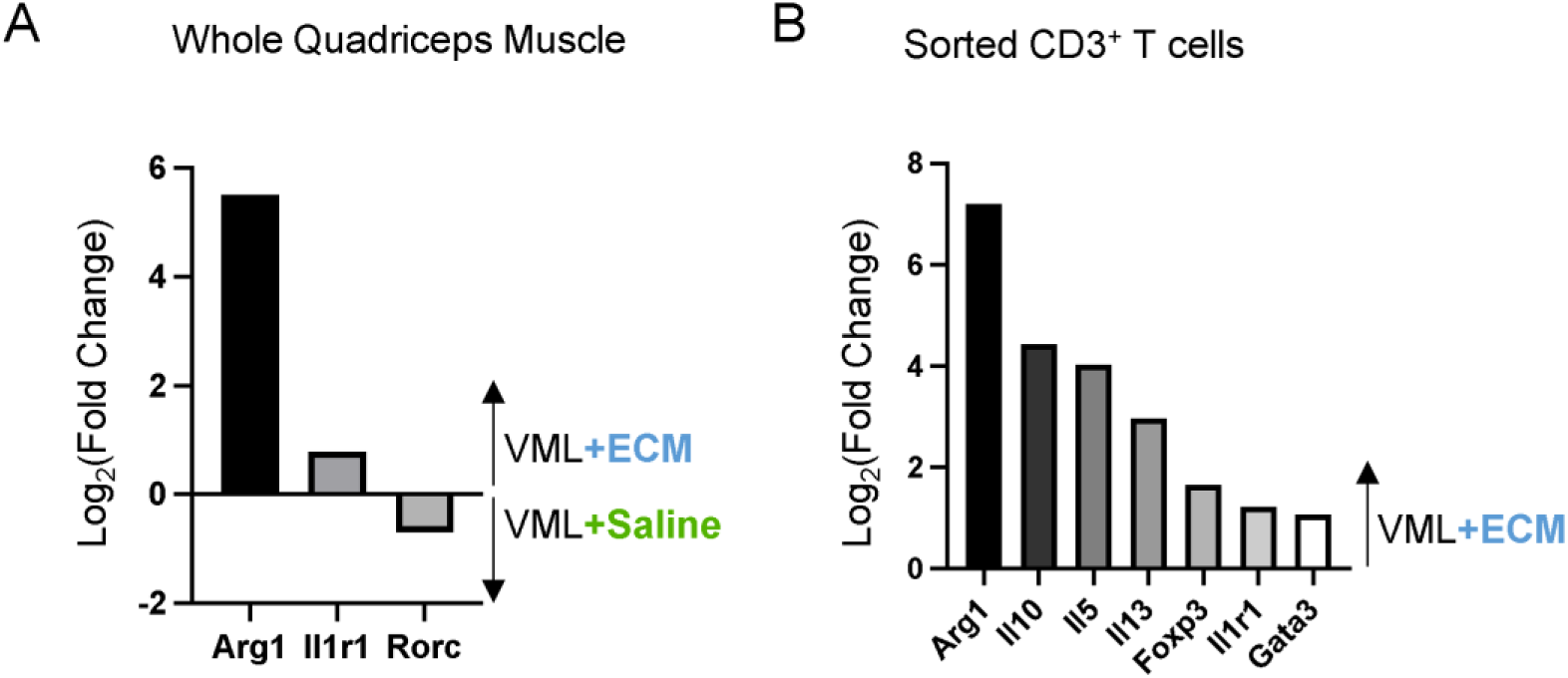
Type 2 immune-related transcriptional signature in 1-week ECM-treated wounds using NanoString PanCancer Immune Profiling panel. (A) Expression of type 2 immune-related genes (*Arg1, Il1r1*) in bulk muscle tissue of ECM-treated VML injuries relative to saline-treated controls. (B) Expression of T_H_2-related (*Arg1, Il5, Il13, Il1r1, Gata3*) and Treg-related (*ll10, Foxp3*) in sorted CD3^+^ T cells from ECM-treated VML injuries relative to saline-treated controls. Data presented as log_2_(fold change) and only depicting significant differentially expressed genes (adj. p<0.05 and |log_2_(FC)|>0.5).

**Fig S2.**
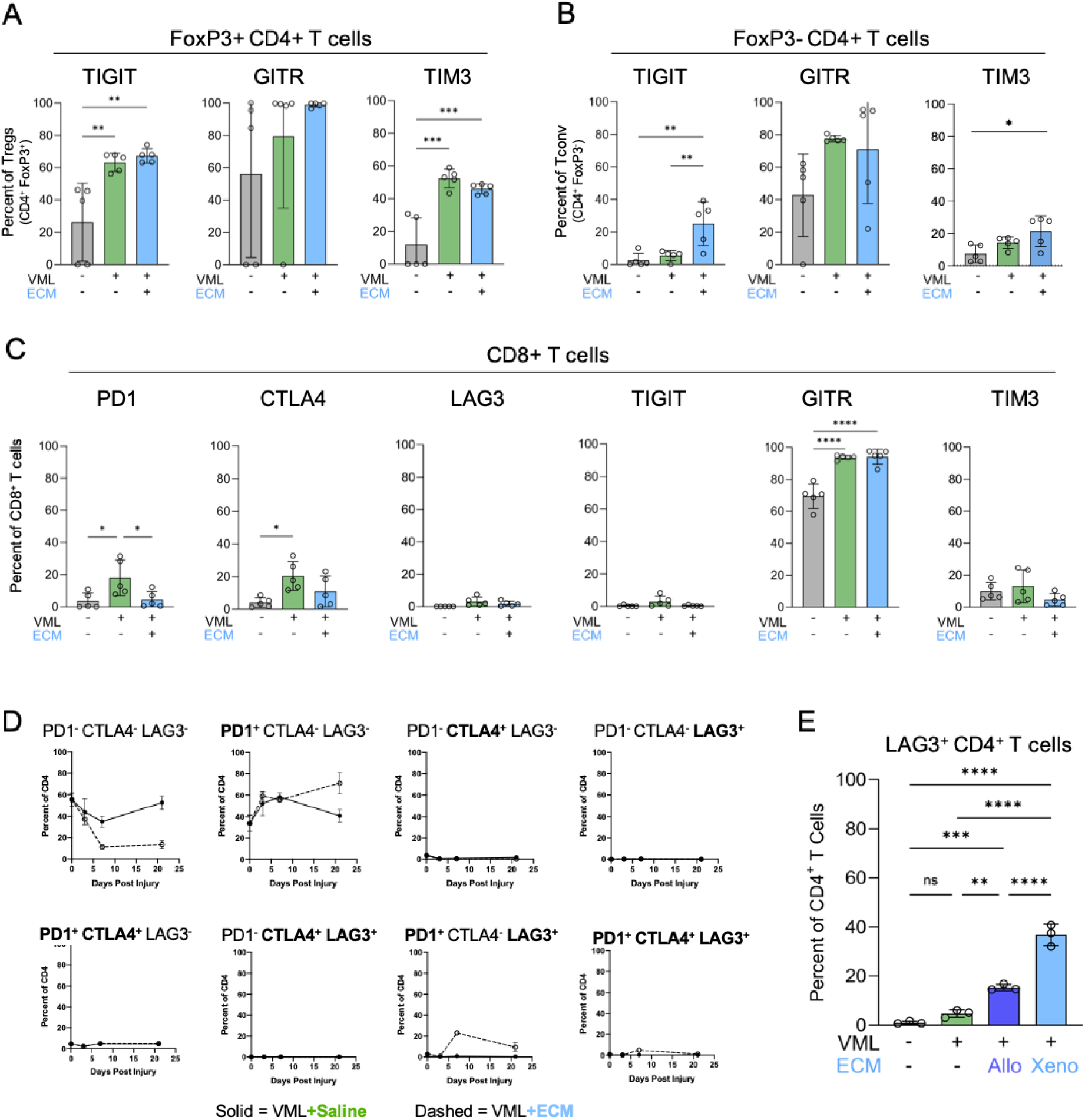
CD4^+^ T cells expressed increased levels of immune checkpoints over a 3-week period following injury with both allogeneic and xenogeneic ECM implants. **(A)** Frequency of TIGIT, GITR, TIM3 protein expression in FoxP3^+^ Tregs in uninjured, saline-treated injuries, or ECM-treated injuries at 1-week. **(B)** Frequency of TIGIT, GITR, TIM3 protein expression in Foxp3^-^ CD4^+^ T cells in uninjured, saline-treated injuries, or ECM-treated injuries at 1-week. **(C)** Frequency of immune checkpoint protein expression in CD8^+^ T cells in uninjured, saline-treated injuries, or ECM-treated injuries at 1-week. **(D)** Time course of PD1, CTLA4, and LAG3 expression in CD4^+^ T cells at 3-days, 1-week, and 3-weeks after saline-treated (solid) or ECM-treated (dashed) injury. **(E)** Frequency of LAG3 expression in CD4^+^ T cells in xenogeneic vs allogeneic ECM-treated injuries at 1-week. Data presented as mean±SD and analyzed using one-way ANOVA with Tukey’s multiple comparisons test (A-C, E). NS p>0.05; * p<0.05; ** p<0.01; *** p<0.001; **** p<0.0001.

**Fig S3.**
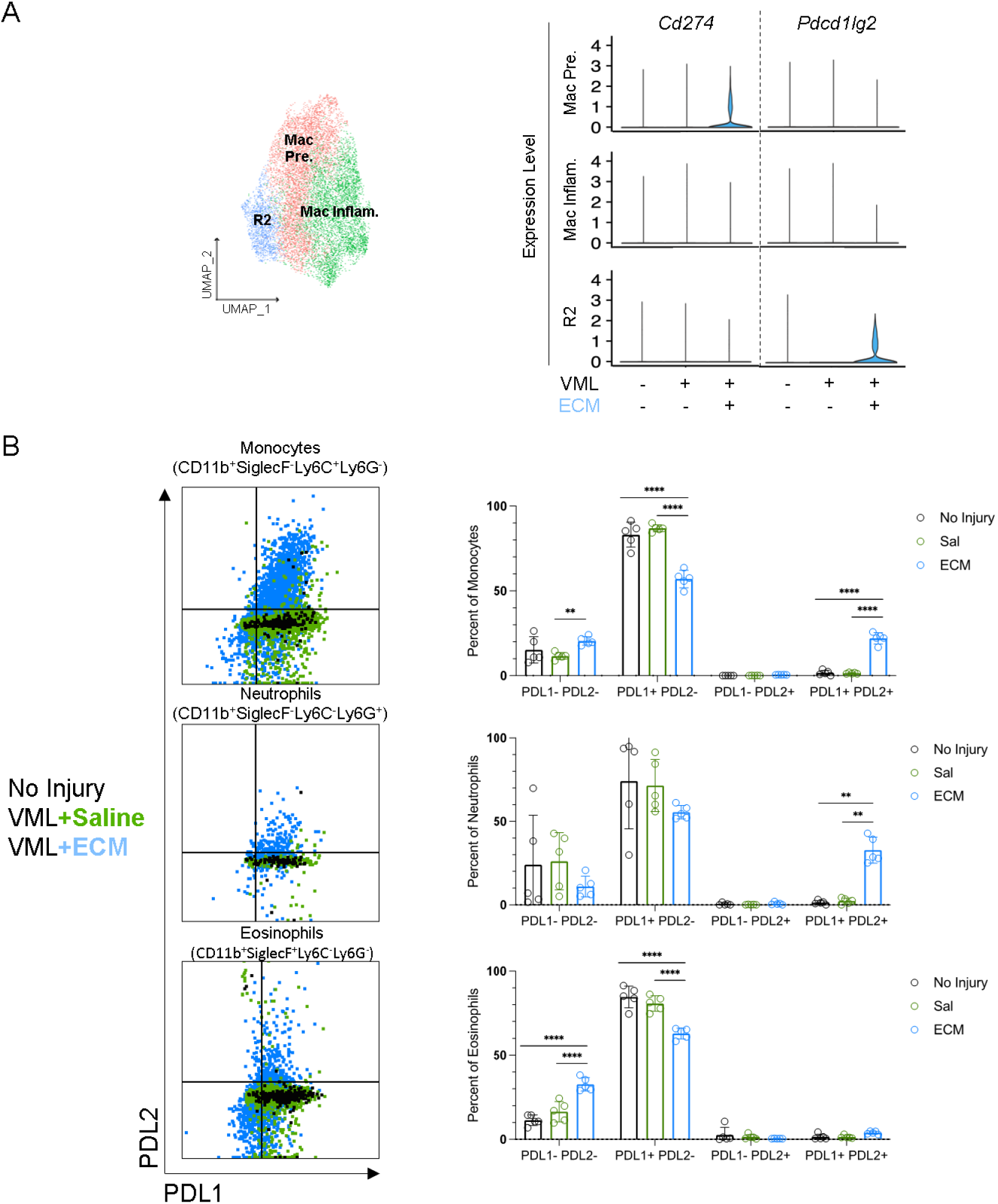
PDL1 and PDL2 expression is present in ECM-treated wounds in some but not all myeloid cell types. **(A)** UMAP of macrophage subclusters from CD45^+^ enriched scRNA-seq dataset collected from uninjured, saline-treated VML, and ECM-treated VML at 1-week. Violin plots for relative expression of *Cd274* (*Pdcd1lg1)* and *Pdcd1lg2*. **(B)** PDL1 and PDL2 protein expression on monocytes, neutrophils, and eosinophils in saline-treated and ECM-treated wounds 1-week after injury. Data presented as mean±SD and analyzed using two-way ANOVA with Tukey’s multiple comparisons test (B). NS p>0.05; * p<0.05; ** p<0.01; *** p<0.001; **** p<0.0001.

**Fig S4.**
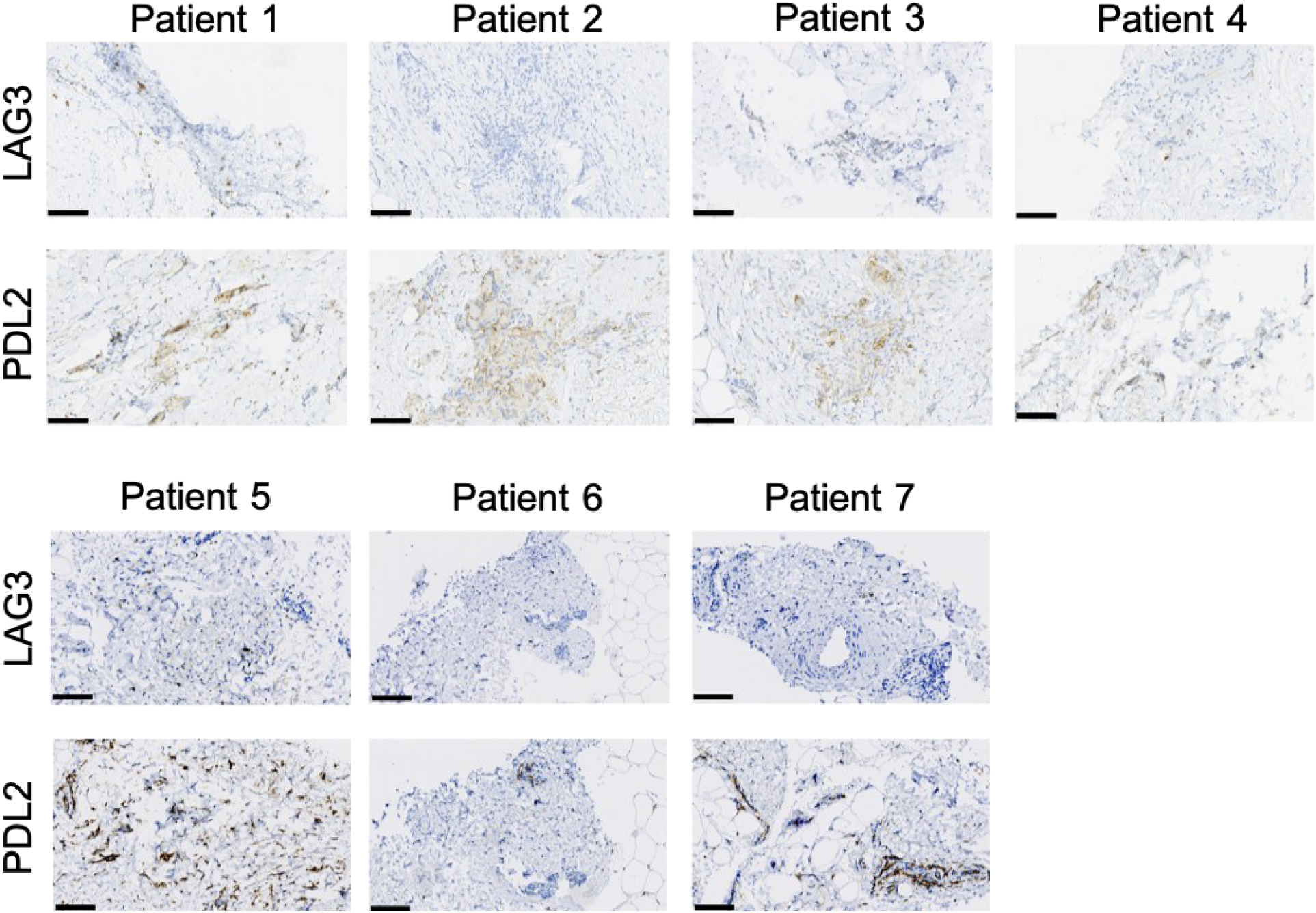
Human ECM implants composed of acellular adipose tissue (AAT) demonstrate LAG3 and PDL2 staining on immunohistochemistry. **(A)** LAG3 and PDL2 immunohistochemistry in seven patients receiving acellular adipose tissue implants for soft tissue reconstruction. Scale bar: 100μm

**Fig S5.**
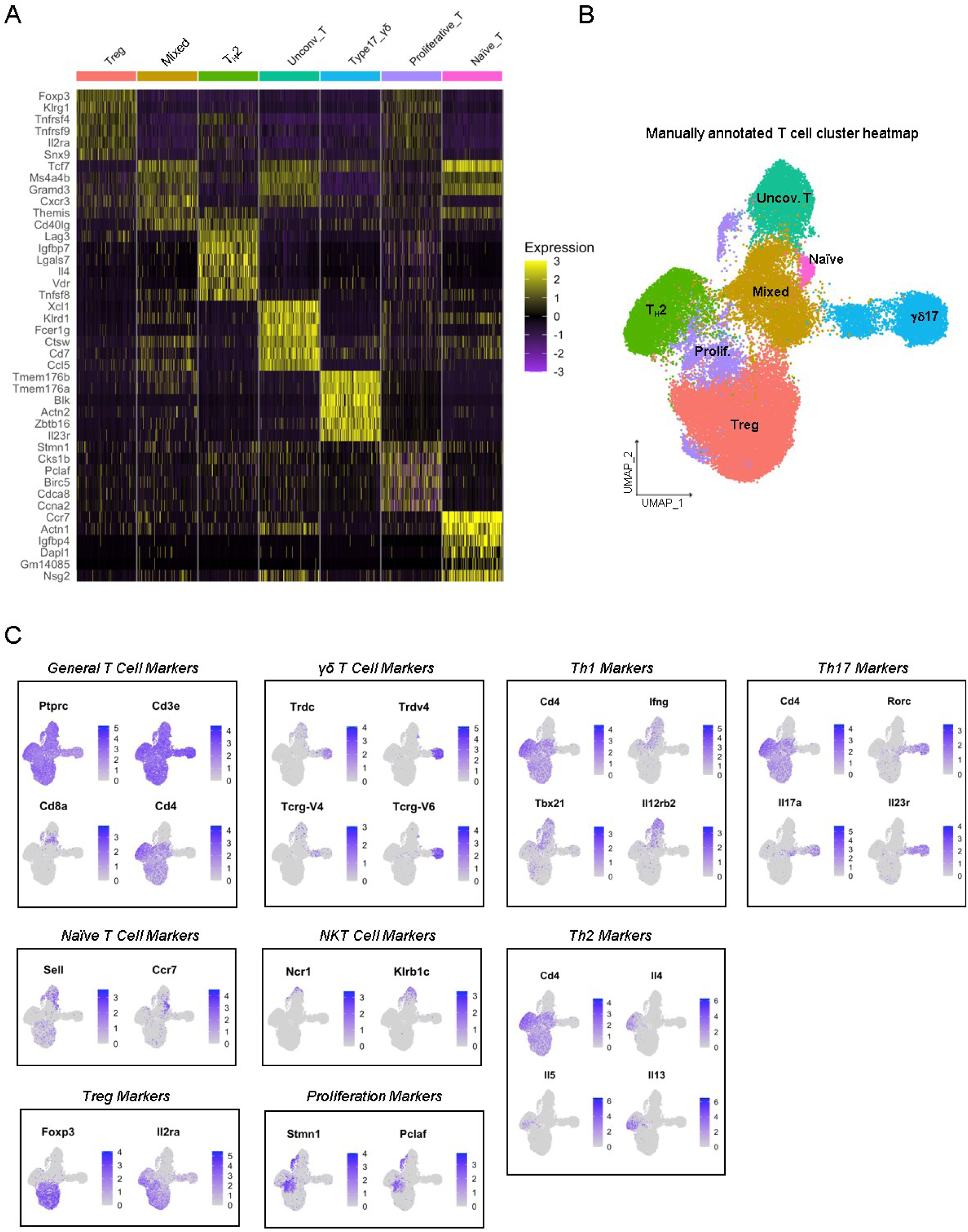
Distinct T cell clusters identified in CD3^+^ T cell scRNA-seq dataset of saline-and ECM-treated VML injuries at 1-week. **(A)** Heatmap of top differentially expressed genes across the seven T cell clusters. **(B)** Representative UMAP projection of the seven T cell clusters identified in the dataset. **(C)** Feature plots of canonical T cell subset markers to validate T cell clusters annotation.

**Fig S6.**
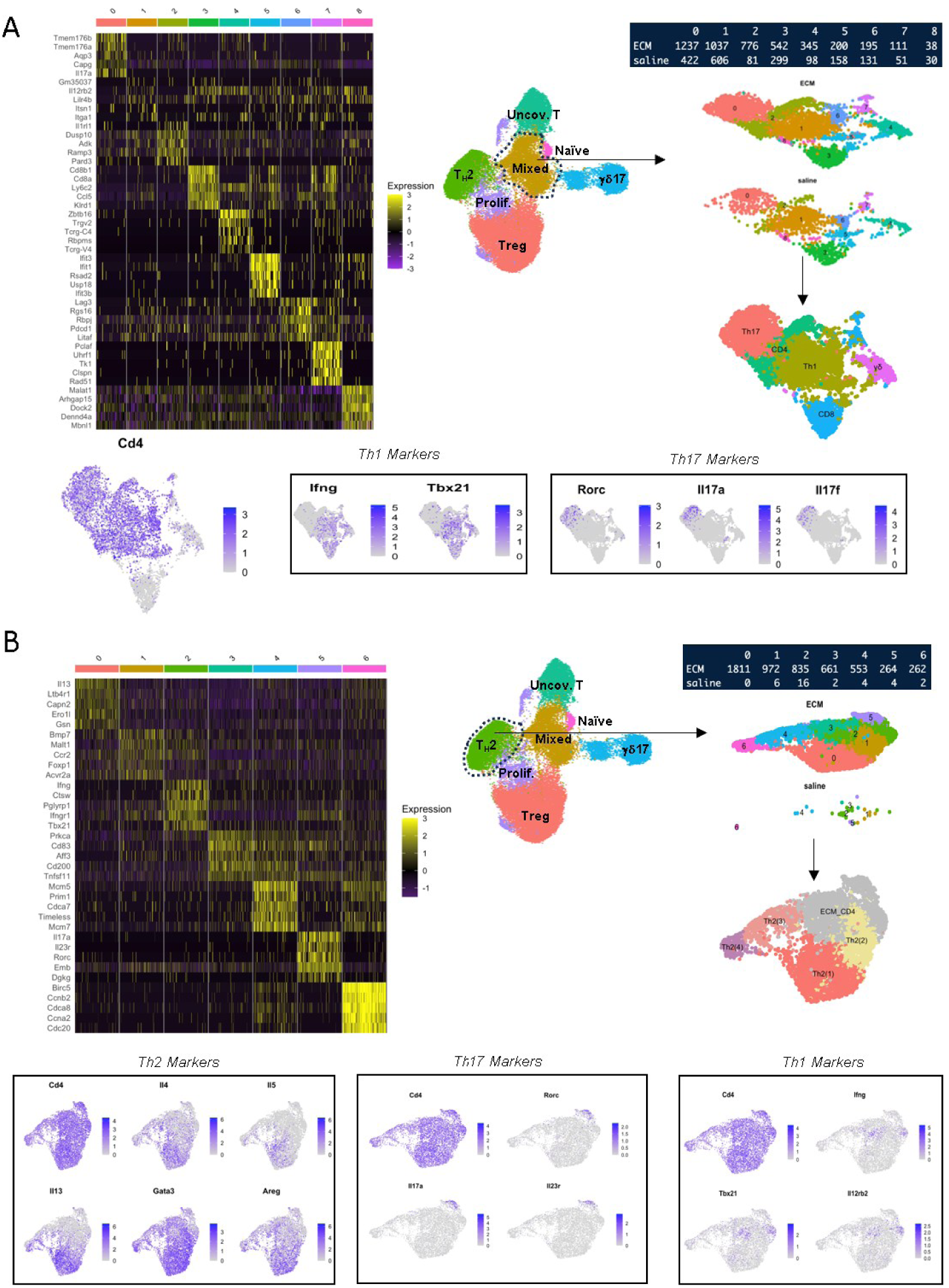
Subcluster analysis shows distinct subclusters consistent with T_H_1, T_H_2, and T_H_17 phenotypes. **(A)** Subcluster analysis was performed on the Mixed T cell cluster. Heat map of top differentially expressed genes per sub-cluster, sub-cluster UMAP, and sub-cluster cell counts by saline and ECM treatment condition. Feature plots of *Cd4*, canonical T_H_1 (*Ifng, Tbx21*), and canonical T_H_17 (*Rorc, Il17a, Il17f*) gene expression. **(B)** Subcluster analysis was performed on the T_H_2 cluster. Heat map of top differentially expressed genes per sub-cluster, sub-cluster UMAP, and sub-cluster cell counts by saline and ECM treatment condition. Feature plots for canonical CD4 helper subset markers clarify subcluster identity including T_H_2 (*Il4, Il5, Il13, Gata3, Areg*), T_H_17 (*Rorc, Il17a, Il23r*), and T_H_1 (*Ifng, Tbx21, Il12rb2*) gene expression. For subsequent data analysis, only sub-clusters that strongly expressed T_H_2 gene signature (0, 1, 4, and 6) were retained and relabeled Th2(1-4). Sub-clusters 2, 3, and 5 were excluded given their relatively high expression of T_H_17 and T_H_1 markers.

**Fig S7.**
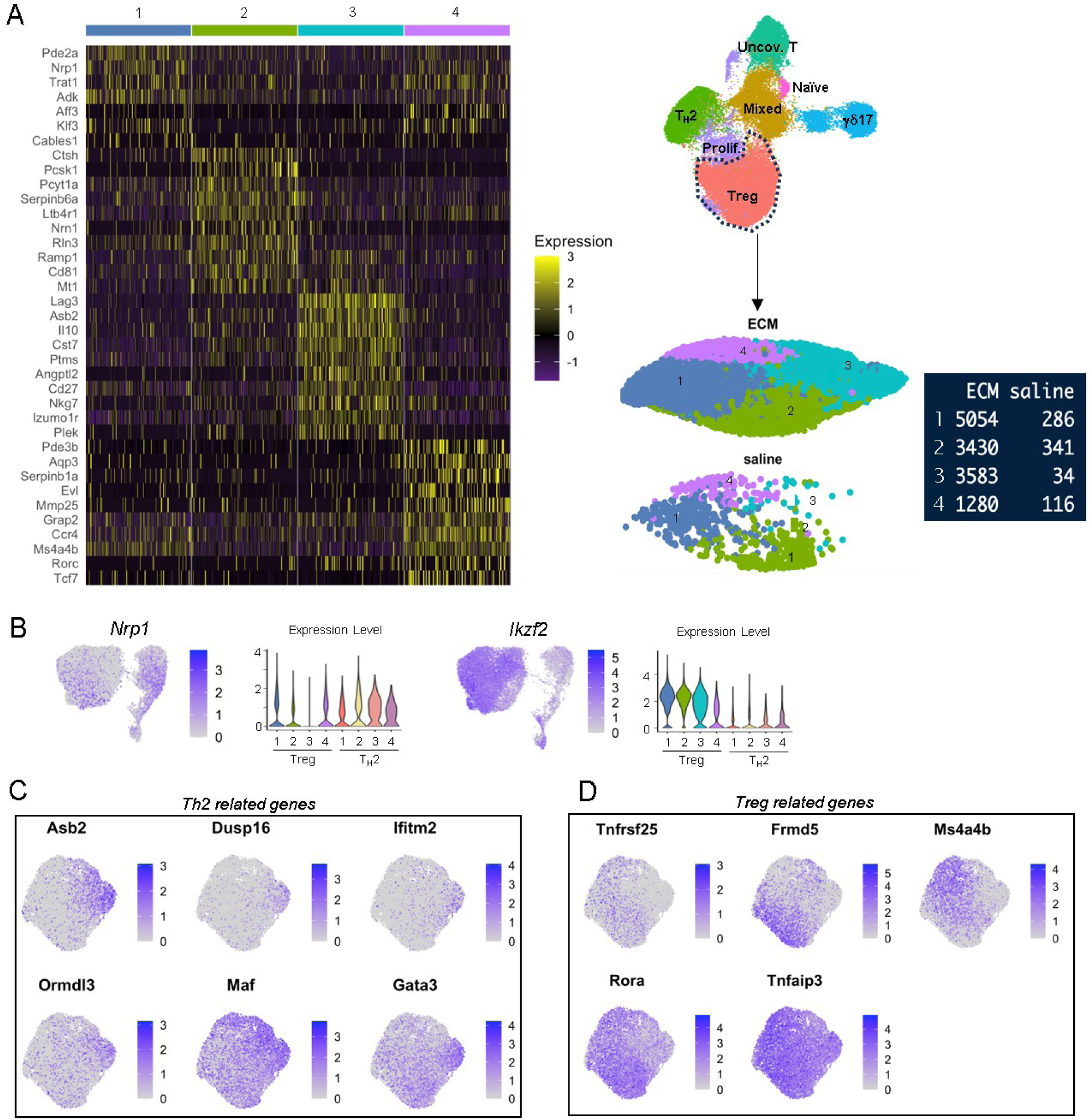
Subcluster analysis shows distinct Treg subclusters with Treg_3 being most present in ECM-treated wounds. (A) Subcluster analysis was performed on the Treg cluster. Heat map of top differentially expressed genes per sub-cluster, sub-cluster UMAP, and sub-cluster cell counts by saline and ECM treatment condition. (B) Feature plots for *Nrp1* and *Ikzf2* on UMAP from combined T_H_2 and Treg populations. Diminished *Nrp1* expression and increased *Ikzf2* expression suggest Treg_3 to be peripherally derived Tregs. (C) T_H_2-related gene feature plots demonstrate overlap in expression on Treg_3 subcluster. (D) Treg-related gene feature plots demonstrate overlap in expression on non-Treg_3 subclusters.

**Fig S8.**
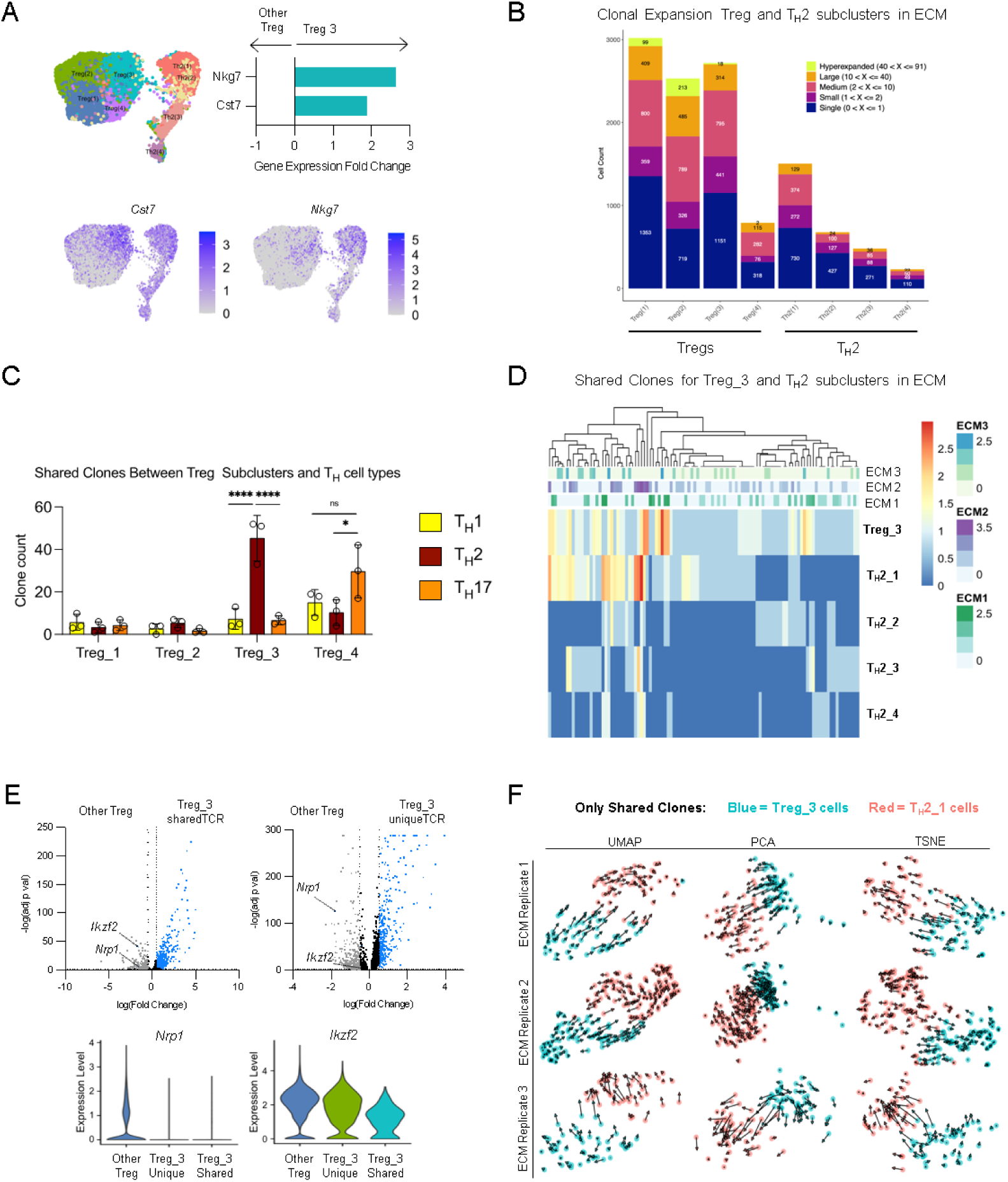
Treg_3 cells share clones (TCR sequences) with T_H_2 cells. **(A)** Expression of *Cst7* and *Nkg7* in Treg_3 relative to other Treg subclusters and T_H_2 cells. **(B)** Clonal expansion of Treg and T_H_2 subclusters in ECM-treated wounds. **(C)** Number of shared TCR clones between each Treg subcluster and T_H_1, T_H_2, and T_H_17 cells. Treg_3 shares the most clones with T_H_2 cells and Treg_4 shares the most clones with T_H_17 cells. **(D)** Heatmap of shared clones between Treg_3 and any T_H_2 subcluster. **(E)** Relative gene expression of Treg_3 cells with shared TCR sequences with T_H_2 cells (Treg_3_sharedTCR) compared to other Treg subclusters. Relative gene expression of Treg_3 cells without shared TCR sequences with T_H_2 cells (Treg_3_uniqueTCR) compared to other Treg subclusters. Violin plots for *Ikzf2* and *Nrp1*. **(F)** RNA velocity diagrams of Treg_3 cells and T_H_2_1 cells with shared TCR sequences by ECM replicate using various visual projection methods (UMAP, PCA, and TSNE) Data presented as mean±SD and analyzed using two-way ANOVA with Tukey’s multiple comparisons test (C). NS p>0.05; * p<0.05; ** p<0.01; *** p<0.001; **** p<0.0001.

**Fig S9.**
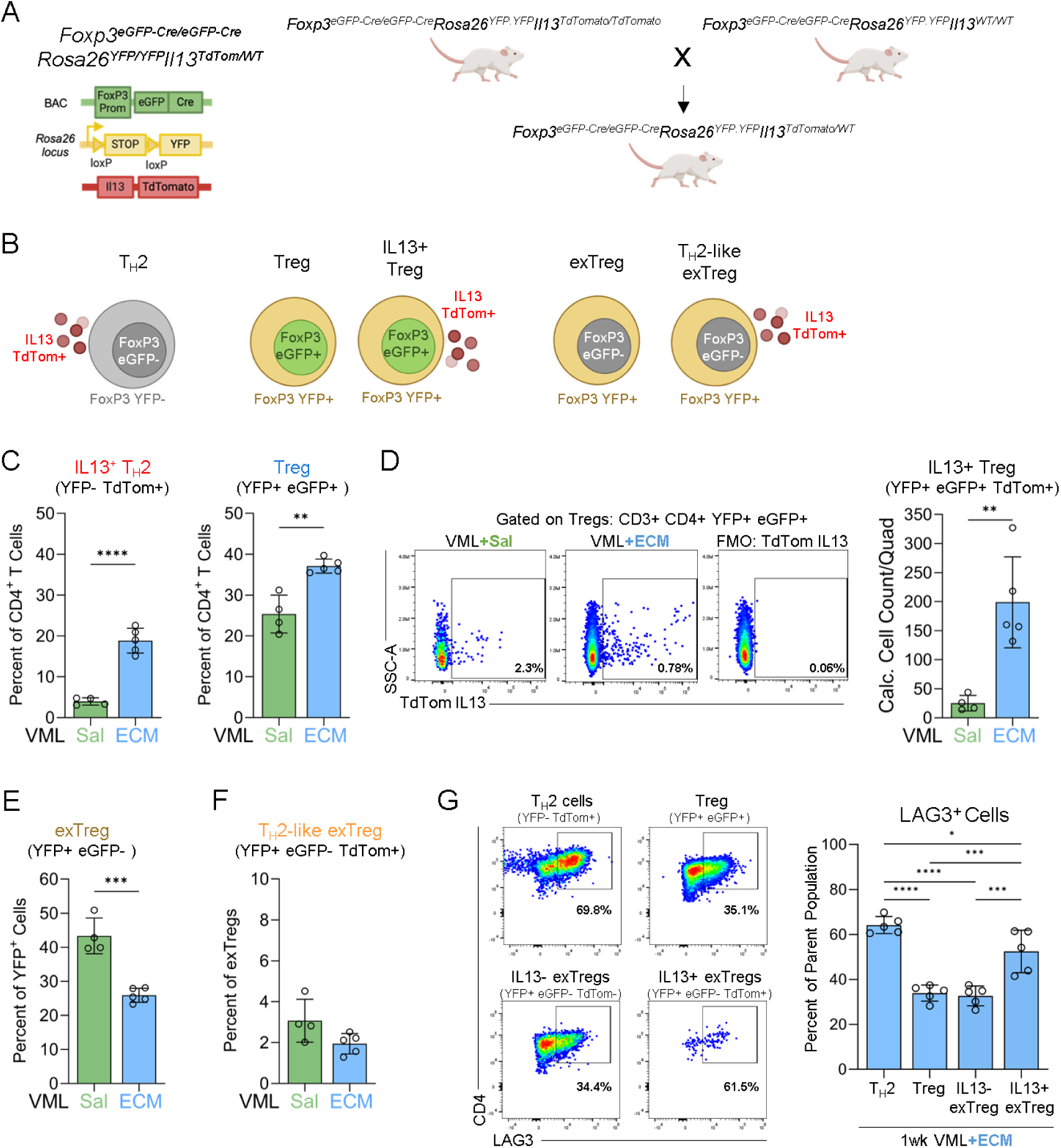
FoxP3^eGFP/YFP^IL13^TdTomato^ reporter mice demonstrate a dynamic response between Tregs and T_H_2 cells in ECM-treated wounds at 1-week. (A) Breeding scheme to generate FoxP3^eGFP/YFP^IL13^TdTomato^ reporter mice from available parent strains. (B) Graphical representation of main CD4^+^ T cell subsets that can be identified with this reporter strain, including IL13^+^ T_H_2 (YFP^-^ TdTom^+^), Tregs (YFP^+^ eGFP^+^), exTregs (YFP^+^ eGFP^-^), and T_H_2-like exTregs (YFP^+^ eGFP^-^ TdTom^+^). (C) Frequency of T_H_2 cells and Tregs out of CD4^+^ T cells in saline-and ECM-treated injuries. (D) Representative flow plots and quantification of IL13^+^ Treg counts/quad in saline-and ECM-treated injuries. (E) Frequency of exTregs out of CD4^+^YFP^+^ T cells in saline-and ECM-treated injuries. (F) Frequency of T_H_2-like exTregs out of exTregs in saline-and ECM-treated injuries. (G) Representative flow plots and quantification of LAG3^+^ expression in various T cell subsets (T_H_2, Tregs, IL13^-^ exTregs, and IL13^+^ exTregs) in saline-and ECM-treated injuries. Data presented as mean±SD and analyzed using unpaired two-tailed T-test (C-F) or one-way ANOVA with Tukey’s multiple comparisons test (G). NS p>0.05; * p<0.05; ** p<0.01; *** p<0.001; **** p<0.0001.

**Fig S10.**
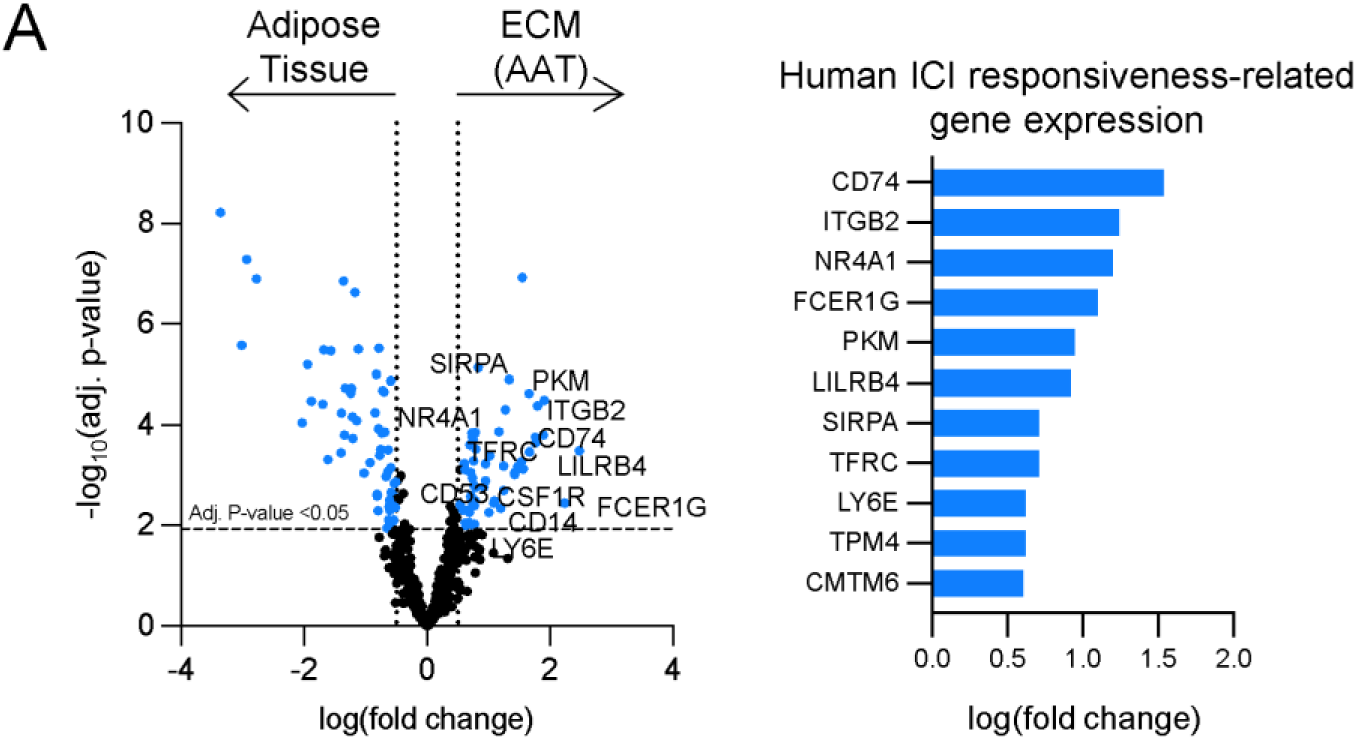
Spatial transcriptomics comparing ICI responsiveness-related genes expression between ECM implants and native adipose tissue in clinical human samples. **(A)** Spatial transcriptomic analysis of differentially expressed genes between ECM implant regions and native adipose tissue in human biopsy samples. Listed genes have been implicated as markers of ICI responsiveness in clinical trials establishing the efficacy of ICI in different tumor settings.

**Fig S11.**
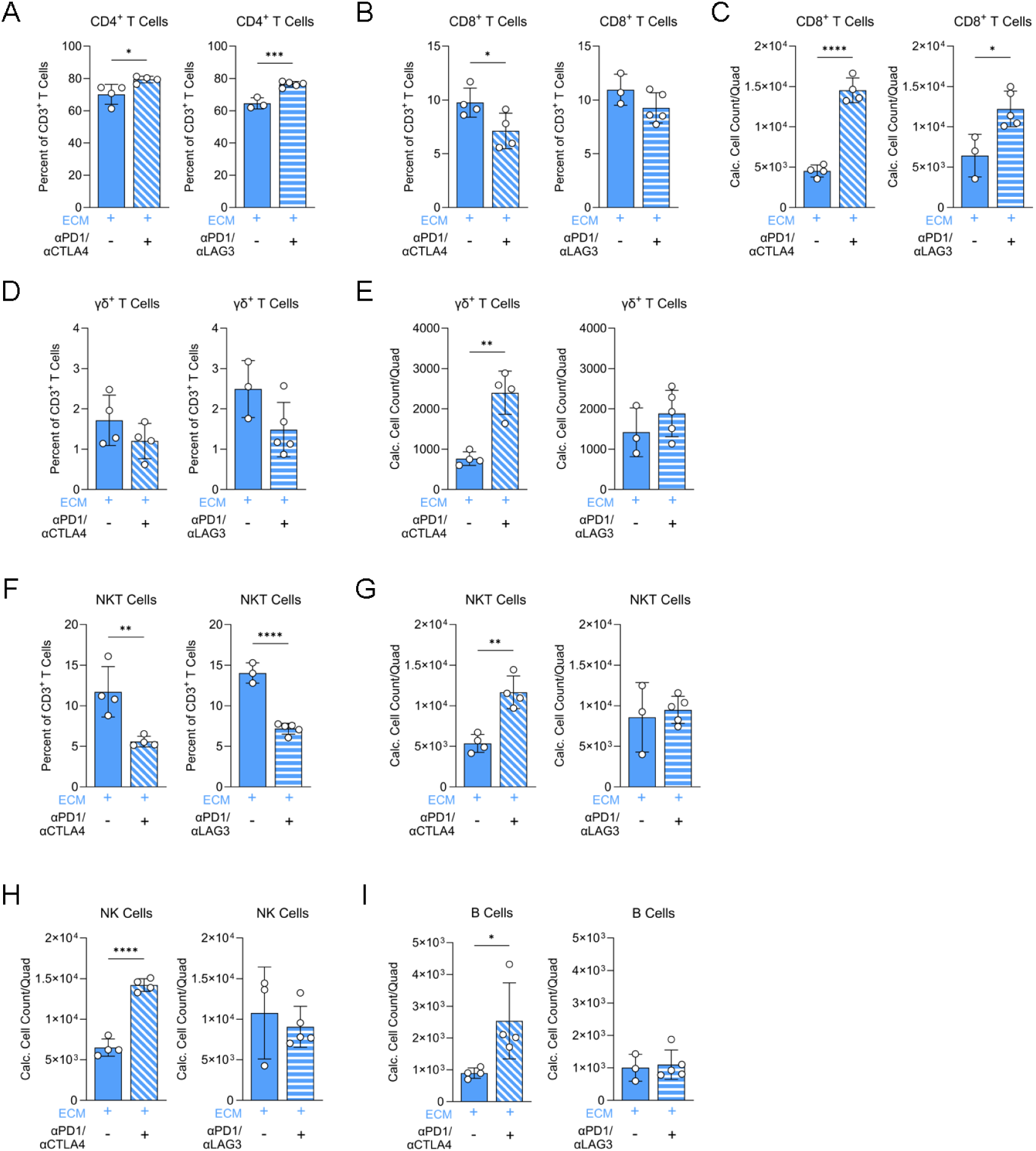
Impact of combination anti-PD1/anti-CTLA4 or anti-PD1/anti-LAG3 treatment on various lymphocyte subsets in ECM-treated injuries at 3-weeks. **(A)** Frequency of CD4^+^ T cells out of CD3^+^ T cells with combination immune checkpoint inhibitors (ICI) in ECM-treated injuries. **(B)** Frequency of CD8^+^ T cells out of CD3^+^ T cells with ICI in ECM-treated injuries. **(C)** Quantification of CD8^+^ T cell counts/quads with ICI in ECM-treated injuries. **(D)** Frequency of γδ T cells out of CD3^+^ T cells with ICI in ECM-treated injuries. **(E)** Quantification of γδ T cell counts/quads with ICI in ECM-treated injuries. **(F)** Frequency of NKT cells out of CD3^+^ T cells with ICI in ECM-treated injuries. **(G)** Quantification of NKT cell counts/quads with ICI in ECM-treated injuries. **(H)** Quantification of NK cell counts/quads with ICI in ECM-treated injuries. **(I)** Quantification of B220^+^ B cell counts/quads with ICI in ECM-treated injuries. Data presented as mean±SD and analyzed using unpaired two-tailed T-test (A-I). NS p>0.05; * p<0.05; ** p<0.01; *** p<0.001; **** p<0.0001.

**Fig S12.**
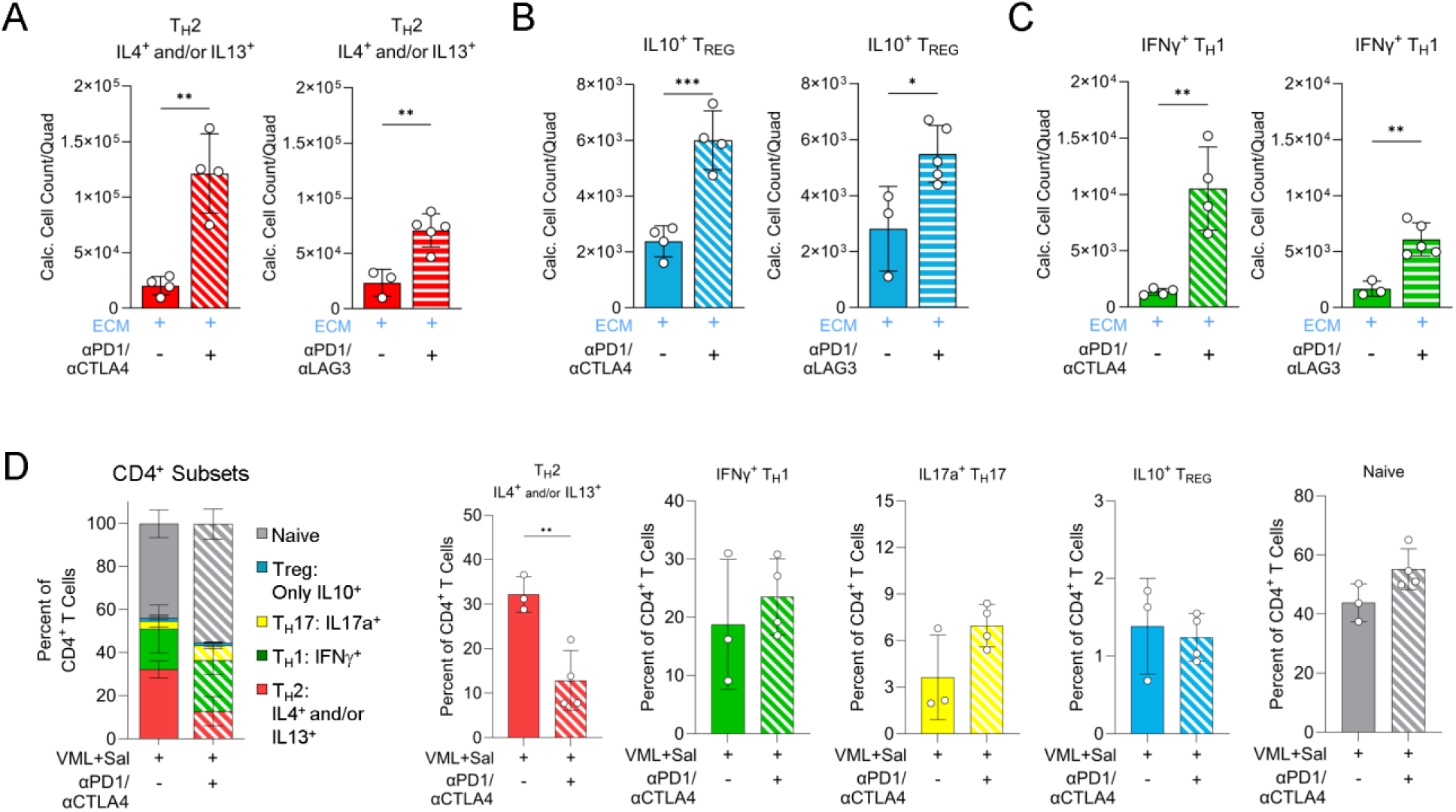
Impact of combination anti-PD1/anti-CTLA4 or anti-PD1/anti-LAG3 treatment on CD4^+^ T cell cytokine expression in saline-or ECM-treated injuries at 3-weeks. (A) Quantification of T_H_2 (lL4^+^ and/or IL13^+^) cell counts/quads with combination immune checkpoint inhibitors (ICI) in ECM-treated injuries. (B) Quantification of IL10^+^ “Treg” (lL4^-^ IL13^-^ IFNγ^-^ IL17a^-^ IL10^+^) counts/quads with ICI in ECM-treated injuries. (C) Quantification of T_H_1 (IFNγ^+^) cell counts/quads with ICI in ECM-treated injuries. (D) Frequency of CD4^+^ T cell subsets (T_H_1, T_H_2, T_H_17, Treg, Naive) with anti-PD1/anti-CLTA4 in saline-treated injuries. Data presented as mean±SD and analyzed using unpaired two-tailed T-test (A-D). NS p>0.05; * p<0.05; ** p<0.01; *** p<0.001; **** p<0.0001.

**Fig S13.**
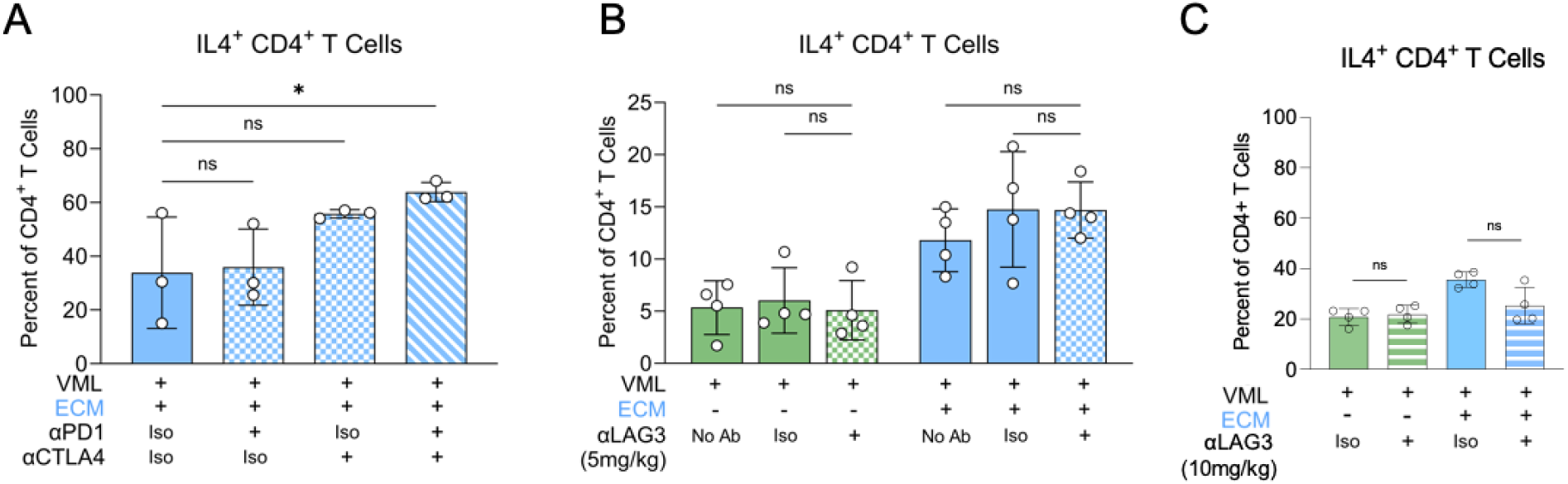
Impact of monotherapy immune checkpoint inhibitors on IL4^+^ T_H_2 responses in saline-or ECM-treated injuries at 3-weeks. **(A)** Frequency of IL4^+^ T_H_2 out of CD4^+^ T cells with isotype, monotherapy anti-PD1, monotherapy anti-CTLA4, or combination anti-PD1/anti-CTLA4 in ECM-treated injuries. **(B)** Frequency of IL4^+^ T_H_2 out of CD4^+^ T cells with no treatment, isotype, or monotherapy anti-LAG3 (standard dose, 5 mg/kg) in saline-and ECM-treated injuries. **(C)** Frequency of IL4^+^ T_H_2 out of CD4^+^ T cells with isotype or monotherapy anti-LAG3 (increased dose, 10 mg/kg) in saline-and ECM-treated injuries. Data presented as mean±SD and analyzed using one-way ANOVA with Dunnett’s multiple comparison relative to isotype control (A) or two-way ANOVA with Tukey’s multiple comparisons test (B-C). NS p>0.05; * p<0.05; ** p<0.01; *** p<0.001; **** p<0.0001.

**Fig S14.**
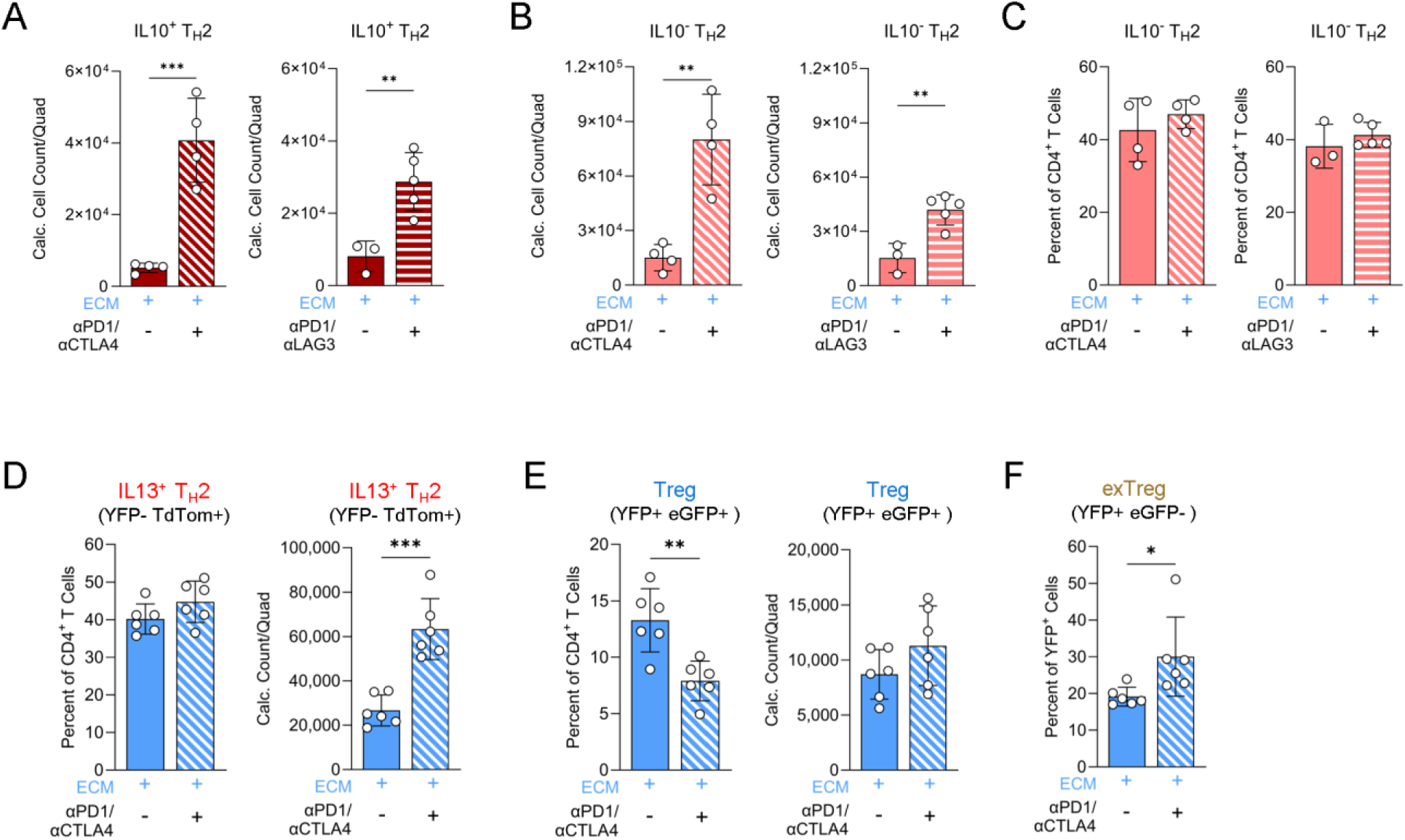
Impact of combination anti-PD1/anti-CTLA4 or anti-PD1/anti-LAG3 treatment on IL10 production by T_H_2 cells and Treg transition to T_H_2-like exTregs in ECM-treated injuries at 3-weeks. (A) Quantification of IL10^+^ T_H_2 (lL4^+^ and/or IL13^+^) cell counts/quads with combination immune checkpoint inhibitors (ICI) in ECM-treated injuries. (B) Quantification of IL10^-^ T_H_2 (lL4^+^ and/or IL13^+^) cell counts/quads with ICI in ECM-treated injuries. (C) Frequency of IL10^+^ T_H_2 out of CD4^+^ T cells with ICI in ECM-treated injuries. (D) Frequency and quantification of IL13^+^ T_H_2 (YFP^-^ TdTom^+^) cell counts/quad with anti-PD1/anti-CTLA4 in ECM-treated injuries using FoxP3^eGFP/YFP^IL13^TdTomato^ reporter mice. (E) Frequency and quantification of Treg (YFP^+^ eGFP^+^) cell counts/quad with anti-PD1/anti-CTLA4 in ECM-treated injuries using FoxP3^eGFP/YFP^IL13^TdTomato^ reporter mice. (F) Frequency of exTregs (YFP^+^ eGFP^-^) out of YFP^+^ cells with anti-PD1/anti-CTLA4 in ECM-treated injuries using FoxP3^eGFP/YFP^IL13^TdTomato^ reporter mice. Data presented as mean±SD and analyzed using unpaired two-tailed T-test (A-F). NS p>0.05; * p<0.05; ** p<0.01; *** p<0.001; **** p<0.0001.

**Fig S15.**
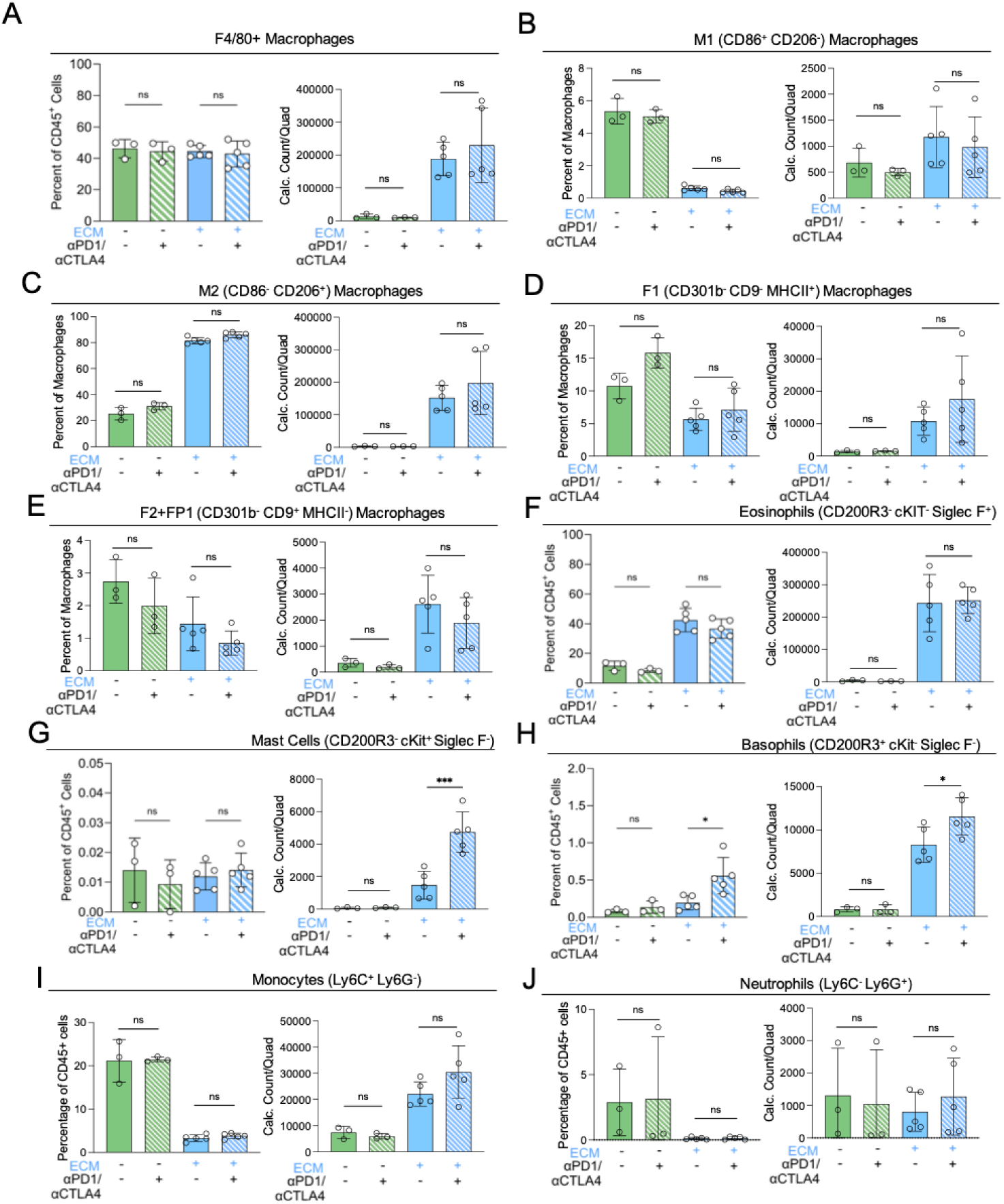
Impact of combination anti-PD1/anti-CTLA4 on macrophage polarization and myeloid subsets in saline-or ECM-treated injuries at 3-weeks. **(A)** Frequency and count of F4/80^+^ macrophages out of CD45^+^ with anti-PD1/anti-CTLA4 (ICI) in saline-and ECM-treated injuries. **(B)** Frequency and count of M1 (CD86^+^CD206^-^) macrophages with ICI in saline-and ECM-treated injuries. **(C)** Frequency and count of M2 (CD86^-^CD206^+^) macrophages with ICI in saline-and ECM-treated injuries. **(D)** Frequency and count of F1 (CD301b^-^ CD9^-^ MHCII^+^) macrophages with ICI in saline-and ECM-treated injuries. **(E)** Frequency and count of F2+FP1 (CD301b^-^ CD9^+^ MHCII^-^) macrophages with ICI in saline-and ECM-treated injuries. **(F)** Frequency and count of SiglecF^+^ eosinophils with ICI in saline-and ECM-treated injuries. **(G)** Frequency and count of CD200R3^-^ cKit^+^ mast cells with ICI in saline-and ECM-treated injuries. **(H)** Frequency and count of CD200R3^+^ cKit^-^ basophils with ICI in saline-and ECM-treated injuries. **(I)** Frequency and count of Ly6C^+^ Ly6G^-^ monocytes with ICI in saline-and ECM-treated injuries. **(J)** Frequency and count of Ly6C^-^ Ly6G^+^ neutrophils with ICI in saline-and ECM-treated injuries. Data presented as mean±SD and analyzed using two-way ANOVA with Tukey’s multiple comparisons test (A-J). NS p>0.05; * p<0.05; ** p<0.01; *** p<0.001; **** p<0.0001.

**Fig S16.**
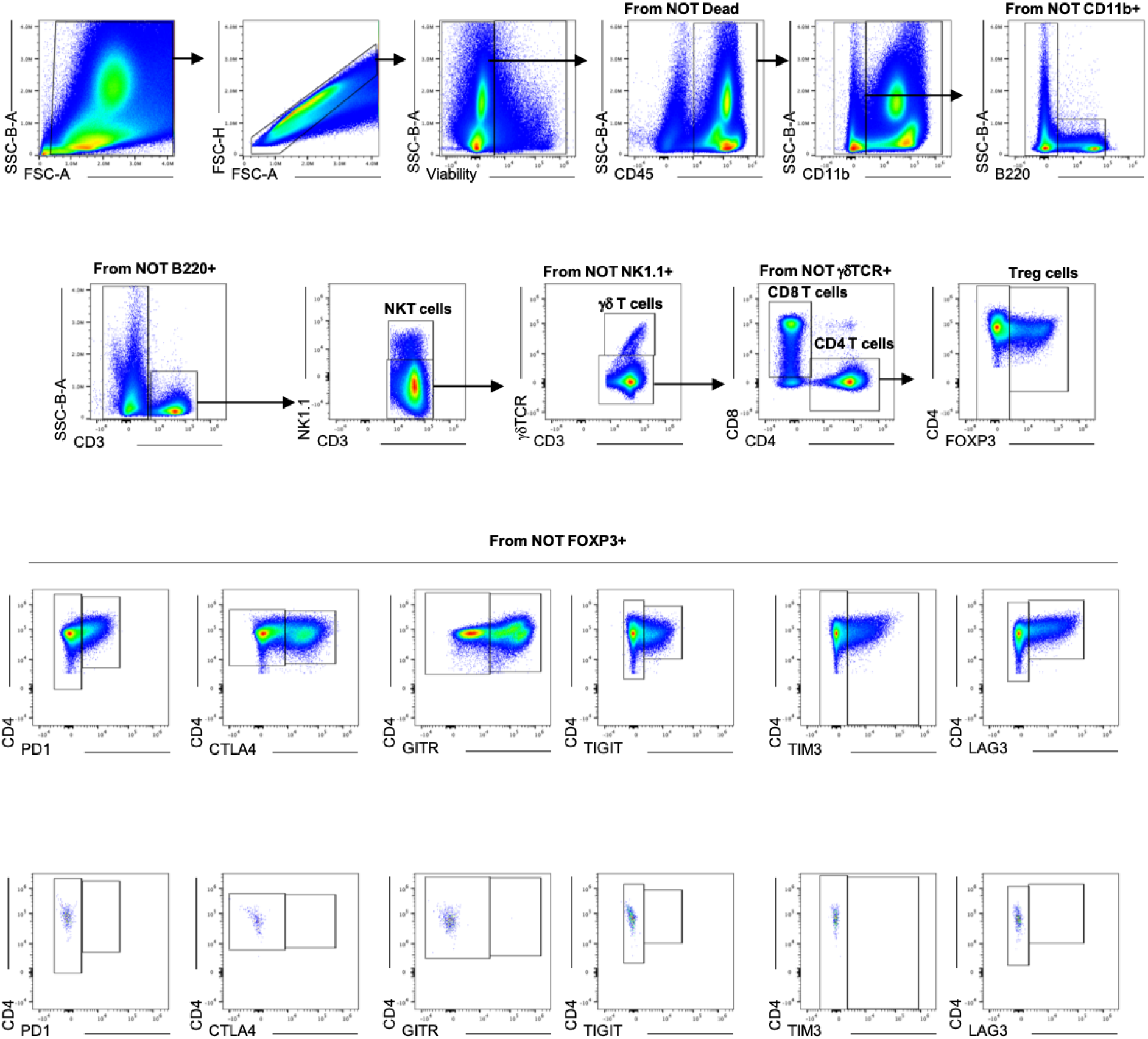
Gating scheme for immune checkpoint expression panel with representative expression of immune checkpoints on conventional CD4^+^ T cells and associated fluorophore-minus-one controls.

**Fig S17.**
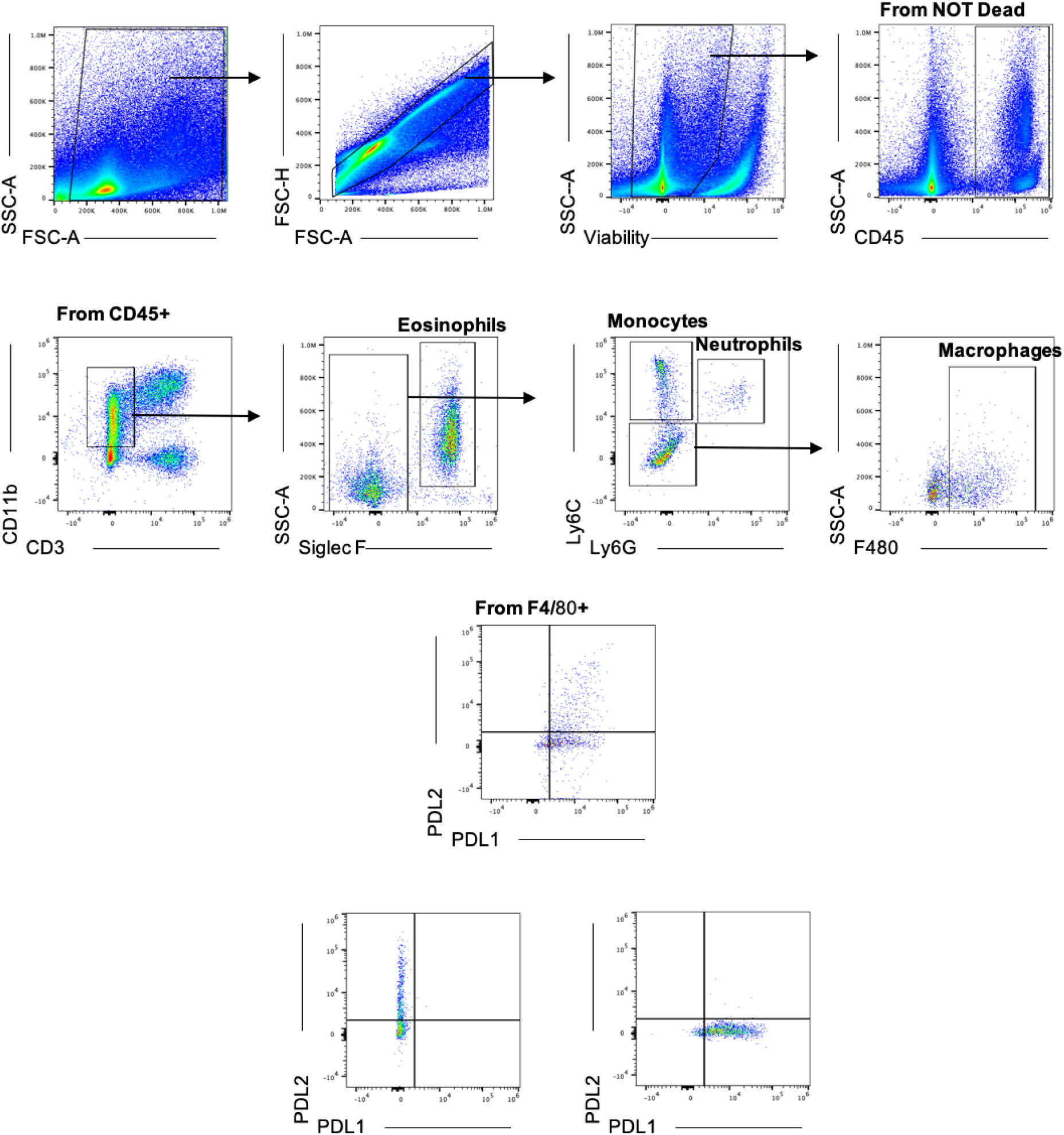
Gating scheme for PDL1/PDL2 checkpoint expression panel with representative expression on conventional F4/80^+^ macrophages and associated fluorophore-minus-one controls.

**Fig S18.**
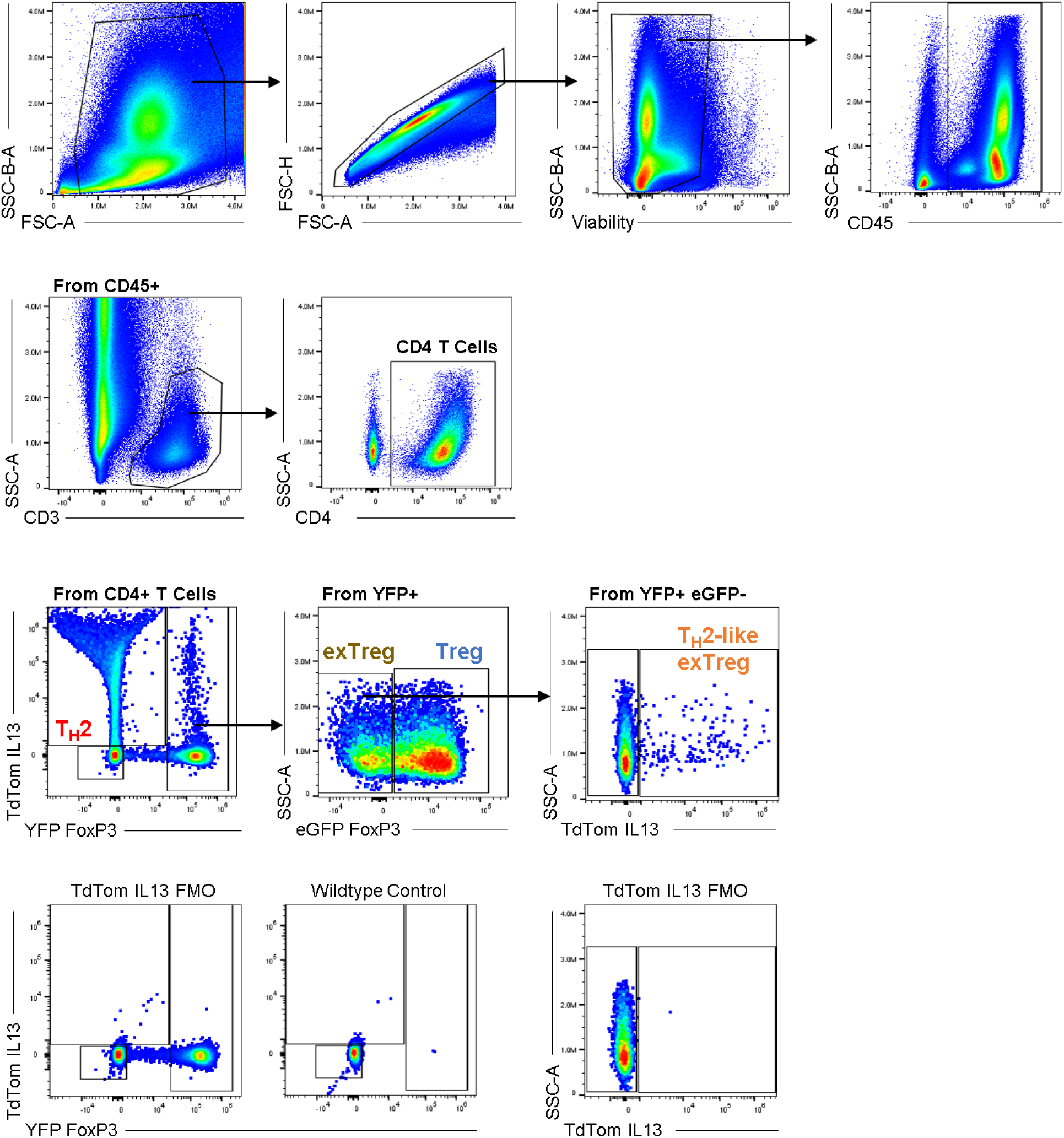
Gating scheme for FoxP3^eGFP/YFP^IL13^TdTomato^ reporter strain panel to identify T_H_2, Treg, exTreg, and T_H_2-like exTreg populations and associated controls.

**Fig S19.**
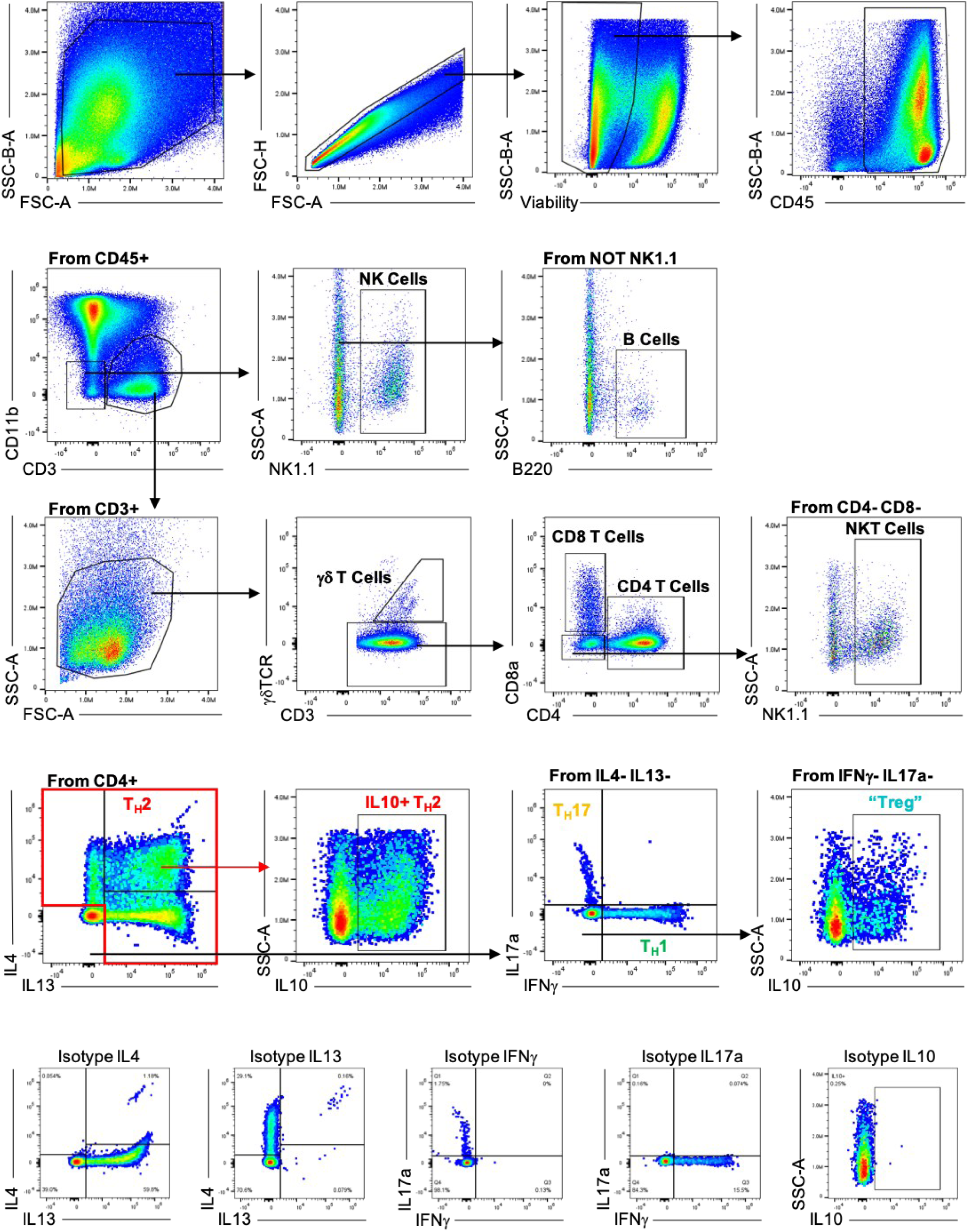
Gating scheme for intracellular cytokine expression panel with representative expression of intracellular cytokines on conventional CD4^+^ T cells and associated isotype controls.

**Fig S20.**
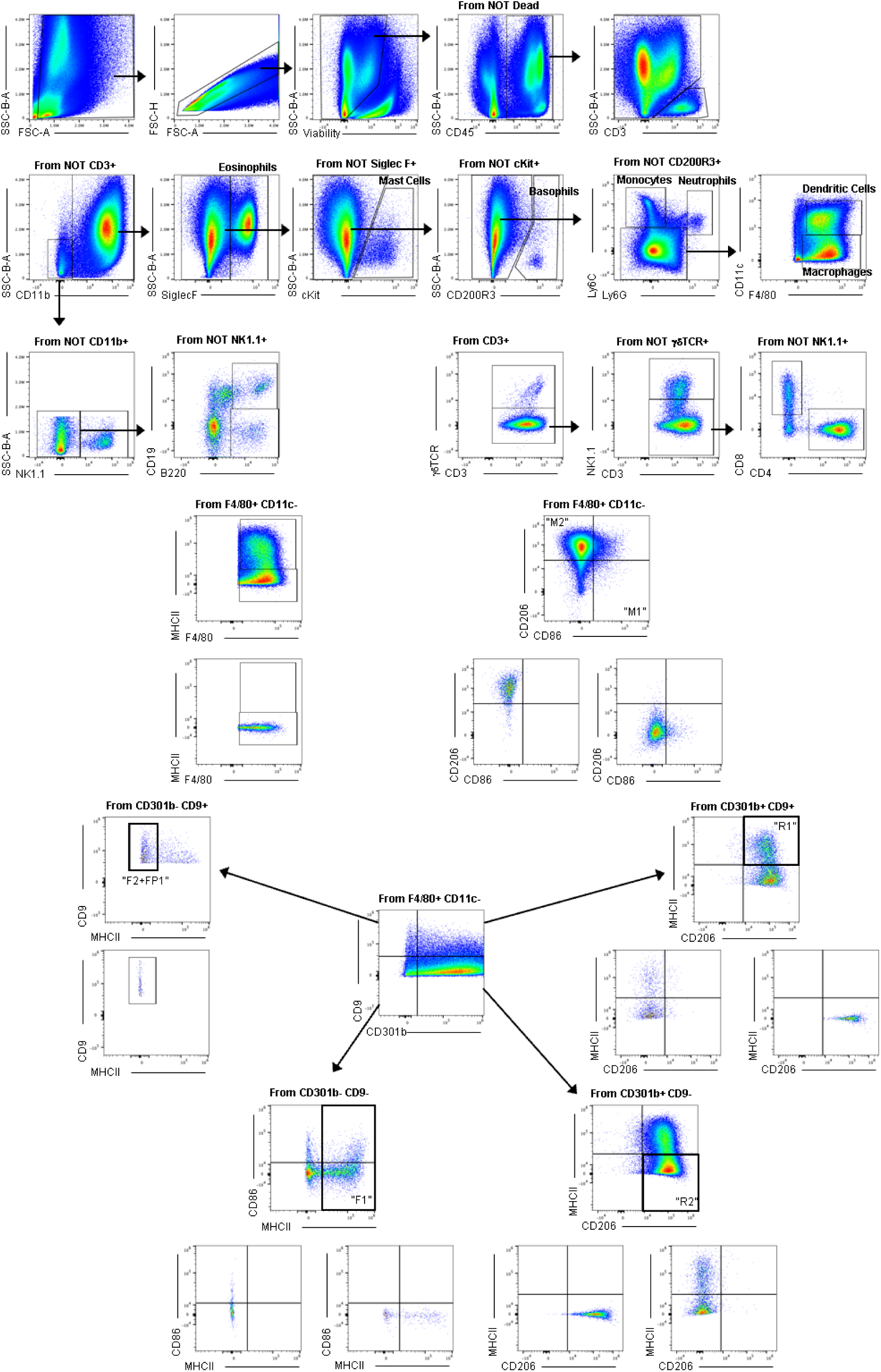
Gating scheme for Pan-Immune panel with representative gating of macrophage subsets and associated fluorophore-minus-one controls.

## Material Tables

**Suppl Table 1:** PDL1/PDL2 Macrophage Profiling Panel (Attune NxT)

| Marker | Fluorophore | Dilution | Clone | Vendor | Catalog | RRID |
| --- | --- | --- | --- | --- | --- | --- |
| Ly6G | Pacific Blue | 1:250 | 1A8 | BioLegend | 127612 | AB_2251161 |
| Ly6C | BV510 | 1:250 | HK1.4 | BioLegend | 128033 | AB_2562351 |
| CD45 | BV605 | 1:150 | 30-F11 | BioLegend | 103140 | AB_2562342 |
| CD4 | BV711 | 1:200 | GK1.5 | BioLegend | 100557 | AB_2562607 |
| CD3 | AF488 | 1:200 | 17A2 | BioLegend | 100210 | AB_389301 |
| PDL2 | PE | 1:200 | TY25 | BioLegend | 107206 | AB_2162011 |
| SiglecF | PE-CF594 | 1:200 | E50-2440 | BD | 562757 | AB_2687994 |
| F4/80 | PE-Cy7 | 1:200 | BM8 | BioLegend | 123114 | AB_893478 |
| PDL1 | APC | 1:200 | 10F.9G2 | BioLegend | 124312 | AB_10612741 |
| CD11b | AF700 | 1:250 | M1/70 | BioLegend | 101222 | AB_493705 |
| Viability | eFluor780 | 1:1000 | N/A | Thermo | 65-0865-18 | N/A |

**Suppl Table 2:**
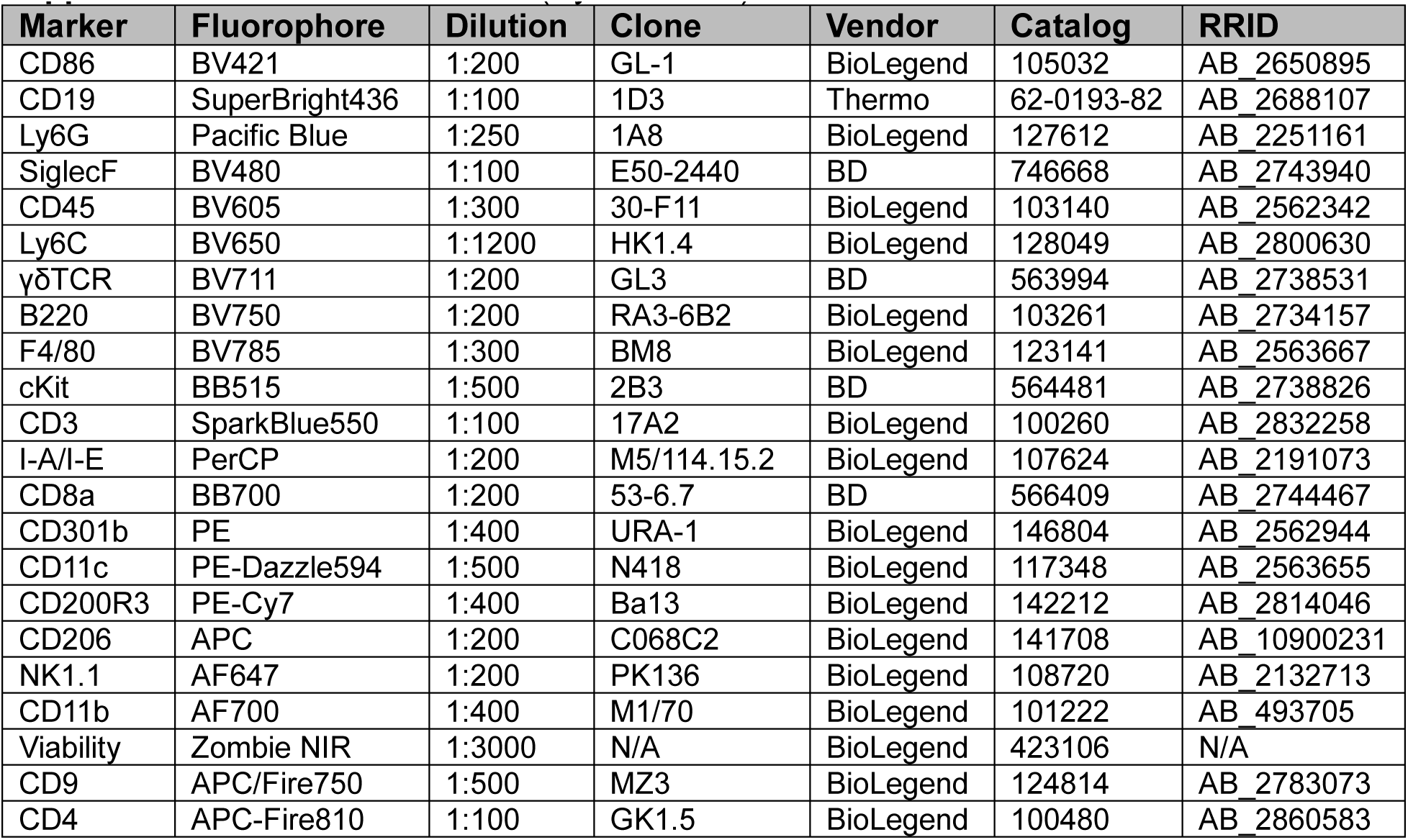
Pan-Immune Panel (Cytek Aurora)

| Marker | Fluorophore | Dilution | Clone | Vendor | Catalog | RRID |
| --- | --- | --- | --- | --- | --- | --- |
| CD86 | BV421 | 1:200 | GL-1 | BioLegend | 105032 | AB_2650895 |
| CD19 | SuperBright436 | 1:100 | 1D3 | Thermo | 62-0193-82 | AB_2688107 |
| Ly6G | Pacific Blue | 1:250 | 1A8 | BioLegend | 127612 | AB_2251161 |
| SiglecF | BV480 | 1:100 | E50-2440 | BD | 746668 | AB_2743940 |
| CD45 | BV605 | 1:300 | 30-F11 | BioLegend | 103140 | AB_2562342 |
| Ly6C | BV650 | 1:1200 | HK1.4 | BioLegend | 128049 | AB_2800630 |
| γδTCR | BV711 | 1:200 | GL3 | BD | 563994 | AB_2738531 |
| B220 | BV750 | 1:200 | RA3-6B2 | BioLegend | 103261 | AB_2734157 |
| F4/80 | BV785 | 1:300 | BM8 | BioLegend | 123141 | AB_2563667 |
| cKit | BB515 | 1:500 | 2B3 | BD | 564481 | AB_2738826 |
| CD3 | SparkBlue550 | 1:100 | 17A2 | BioLegend | 100260 | AB_2832258 |
| I-A/I-E | PerCP | 1:200 | M5/114.15.2 | BioLegend | 107624 | AB_2191073 |
| CD8a | BB700 | 1:200 | 53-6.7 | BD | 566409 | AB_2744467 |
| CD301b | PE | 1:400 | URA-1 | BioLegend | 146804 | AB_2562944 |
| CD11c | PE-Dazzle594 | 1:500 | N418 | BioLegend | 117348 | AB_2563655 |
| CD200R3 | PE-Cy7 | 1:400 | Ba13 | BioLegend | 142212 | AB_2814046 |
| CD206 | APC | 1:200 | C068C2 | BioLegend | 141708 | AB_10900231 |
| NK1.1 | AF647 | 1:200 | PK136 | BioLegend | 108720 | AB_2132713 |
| CD11b | AF700 | 1:400 | M1/70 | BioLegend | 101222 | AB_493705 |
| Viability | Zombie NIR | 1:3000 | N/A | BioLegend | 423106 | N/A |
| CD9 | APC/Fire750 | 1:500 | MZ3 | BioLegend | 124814 | AB_2783073 |
| CD4 | APC-Fire810 | 1:100 | GK1.5 | BioLegend | 100480 | AB_2860583 |

**Suppl Table 3:** T Cell Immune Checkpoint Panel (Cytek Aurora)

| Marker | Fluorophore | Dilution | Clone | Vendor | Catalog | RRID |
| --- | --- | --- | --- | --- | --- | --- |
| NK1.1 | BV421 | 1:50 | PK136 | BioLegend | 108732 | AB_2562218 |
| GITR | BV480 | 1:200 | DTA-1 | BD | 746745 | AB_2744008 |
| CD45 | BV510 | 1:200 | 30-F11 | BioLegend | 103138 | AB_2563061 |
| CD4 | BV605 | 1:300 | GK1.5 | BioLegend | 100451 | AB_2564591 |
| γδTCR | BV711 | 1:200 | GL3 | BD | 563994 | AB_2738531 |
| B220 | BV750 | 1:200 | RA3-6B2 | BioLegend | 103261 | AB_2734157 |
| PD1 | BV785 | 1:200 | RMP1-30 | BD | 748264 | AB_2872692 |
| TCF1/7 | AF488 | 1:100 | C63D9 | Cell Signaling | 6444S | AB_2797627 |
| CD3 | SparkBlue550 | 1:100 | 17A2 | BioLegend | 100260 | AB_2832258 |
| CD8a | BB700 | 1:200 | 53-6.7 | BD | 566409 | AB_2744467 |
| TIGIT | PerCP-eFluor710 | 1:100 | GIGD7 | Thermo | 46-9501-82 | AB_11150967 |
| CD25 | PE | 1:150 | PC61 | BioLegend | 102008 | AB_312857 |
| CTLA4 | PE-Dazzle594 | 1:200 | UC10-4B9 | BioLegend | 106318 | AB_2564496 |
| TIM3 | PE-Cy7 | 1:150 | RMT3-23 | BioLegend | 119716 | AB_2571933 |
| TOX | APC | 1:150 | REA473 | Miltenyi Biotec | 130-118-335 | AB_2751485 |
| FoxP3 | eFluor660 | 1:100 | FJK-16s | Thermo | 50-5773-82 | AB_11218868 |
| CD11b | AF700 | 1:400 | M1/70 | BioLegend | 101222 | AB_493705 |
| Viability | ZombieNIR | 1:5000 | N/A | BioLegend | 423106 | N/A |
| LAG3 | APC-eFluor780 | 1:100 | C9B7W | Thermo | 47-2231-82 | AB_2637323 |

Markers shown in blue are stained as intracellular

**Suppl Table 4:** Intracellular Cytokine Panel (Cytek Aurora)

| Marker | Fluorophore | Dilution | Clone | Vendor | Catalog | RRID |
| --- | --- | --- | --- | --- | --- | --- |
| IL5 | BV421 | 1:100 | TRFK5 | BioLegend | 504311 | AB_2563161 |
| CD45 | BV510 | 1:200 | 30-F11 | BioLegend | 103138 | AB_2563061 |
| CD4 | BV605 | 1:300 | GK1.5 | BioLegend | 100451 | AB_2564591 |
| CD11b | BV650 | 1:500 | M1/70 | BioLegend | 101259 | AB_2566568 |
| γδTCR | BV711 | 1:200 | GL3 | BD | 563994 | AB_2738531 |
| B220 | BV750 | 1:200 | RA3-6B2 | BioLegend | 103261 | AB_2734157 |
| IL10 | Vio B515 | 1:100 | REA1008 | Miltenyi Biotec | 130-116-848 | AB_2727720 |
| CD3 | SparkBlue550 | 1:100 | 17A2 | BioLegend | 100260 | AB_2832258 |
| CD8a | BB700 | 1:100 | 53-6.7 | BD | 566409 | AB_2744467 |
| IL4 | PE | 1:100 | 11B11 | BioLegend | 504104 | AB_315318 |
| IL13 | PE-eFluor610 | 1:250 | eBio13A | Thermo | 61-7133-82 | AB_2574654 |
| IFNγ | APC | 1:150 | XMG1.2 | BioLegend | 505810 | AB_315404 |
| NK1.1 | AF647 | 1:200 | PK136 | BioLegend | 108720 | AB_2132713 |
| IL17a | AF700 | 1:200 | TC11-18H10.1 | BioLegend | 506914 | AB_536016 |
| Viability | ZombieNIR | 1:5000 | N/A | BioLegend | 423106 | N/A |
| Isotypes |  |  |  |  |  |  |
| Rat IgG1 | BV421 | 1:100 | RTK2071 | BioLegend | 400429 | AB_10900998 |
| Hu IgG1 | Vio B515 | 1:100 | REA293 | Miltenyi Biotec | 130-114-556 | AB_2751239 |
| Rat IgG1 | PE | 1:100 | RTK2071 | BioLegend | 400407 | AB_326513 |
| Rat IgG1 | PE-eFluor610 | 1:250 | eBRG1 | Thermo | 61-4301-82 | AB_2637348 |
| Rat IgG1 | APC | 1:150 | RTK2071 | BioLegend | 400411 | AB_326517 |
| Rat IgG1 | AF700 | 1:200 | RTK2071 | BioLegend | 400420 | AB_493781 |

Markers shown in blue are stained as intracellular

**Suppl Table 5:** Intracellular Cytokine Panel (Attune NxT)

| Marker | Fluorophore | Dilution | Clone | Vendor | Catalog | RRID |
| --- | --- | --- | --- | --- | --- | --- |
| FoxP3 | BV421 | 1:500 | MF-14 | BioLegend | 126419 | AB_2565933 |
| CD45 | V500 | 1:150 | 30-F11 | BD | 561487 | AB_10697046 |
| NK1.1 | BV605 | 1:300 | PK136 | BioLegend | 108740 | AB_2562274 |
| CD8a | BV711 | 1:300 | 53-6.7 | BioLegend | 100748 | AB_2562100 |
| CD3 | AF488 | 1:200 | 17A2 | BioLegend | 100210 | AB_389301 |
| CD19 | PerCP-Cy5.5 | 1:200 | 6D5 | BioLegend | 115534 | AB_2072925 |
| IL4 | PE | 1:200 | 11B11 | BioLegend | 504104 | AB_315318 |
| γδTCR | PE-CF594 | 1:300 | GL3 | BD | 563532 | AB_2661844 |
| CD4 | PE-Cy7 | 1:300 | GK1.5 | BioLegend | 100422 | AB_312707 |
| IFNγ | APC | 1:200 | XMG1.2 | BioLegend | 505810 | AB_315404 |
| IL17a | AF700 | 1:200 | TC11-18H10.1 | BioLegend | 506914 | AB_536016 |
| Viability | eFluor780 | 1:1000 | N/A | Thermo | 65-0865-18 | N/A |

Markers shown in blue are stained as intracellular

**Suppl Table 6:** FoxP3^eGFP/YFP^IL13^TdTomato^ Reporter Mouse Panel (Cytek Aurora)

| Marker | Fluorophore | Dilution | Clone | Vendor | Catalog | RRID |
| --- | --- | --- | --- | --- | --- | --- |
| CD45 | BV421 | 1:200 | 30-F11 | BioLegend | 103151 | AB_2565884 |
| LAG3 | BV875 | 1:100 | C9B7W | BioLegend | 125219 | AB_2566571 |
| FoxP3 | eGFP | N/A | N/A | N/A | N/A | N/A |
| FoxP3 | YFP | N/A | N/A | N/A | N/A | N/A |
| IL13 | tdTomato | N/A | N/A | N/A | N/A | N/A |
| CD3 | APC | 1:100 | 17A2 | BioLegend | 100236 | AB_2561456 |
| CD11b | AF700 | 1:400 | M1/70 | BioLegend | 101222 | AB_493705 |
| Viability | ZombieNIR | 1:5000 | N/A | BioLegend | 423106 | N/A |
| CD4 | APC-Fire810 | 1:100 | GK1.5 | BioLegend | 100480 | AB_2860583 |

**Suppl Table 7:** T Cell Sorting Panel for scRNASeq (BD FACSAria)

| Marker | Fluorophore | Dilution | Clone | Vendor | Catalog | RRID |
| --- | --- | --- | --- | --- | --- | --- |
| CD45 | BV605 | 1:150 | 30-F11 | BioLegend | 103140 | AB_2562342 |
| CD3 | AF488 | 1:200 | 17A2 | BioLegend | 100210 | AB_389301 |
| Viability | ZombieNIR | 1:2000 | N/A | BioLegend | 423106 | N/A |
| Marker | Hashing Oligo | Dilution | Clone | Vendor | Catalog | RRID |
| MHCI; CD45 | TotalSeq-C0301 | 1:100 | M1/42; 30-F11 | BioLegend | 155861 | AB_2800693 |
| MHCI; CD45 | TotalSeq-C0302 | 1:100 | M1/42; 30-F11 | BioLegend | 155863 | AB_2800694 |
| MHCI; CD45 | TotalSeq-C0303 | 1:100 | M1/42; 30-F11 | BioLegend | 155865 | AB_2800695 |
| MHCI; CD45 | TotalSeq-C0304 | 1:100 | M1/42; 30-F11 | BioLegend | 155867 | AB_2800696 |
| MHCI; CD45 | TotalSeq-C0305 | 1:100 | M1/42; 30-F11 | BioLegend | 155869 | AB_2800697 |

**Suppl Table 8:** T Cell Sorting Panel for NanoString (BD FACSAria)

| Marker | Fluorophore | Dilution | Clone | Vendor | Catalog | RRID |
| --- | --- | --- | --- | --- | --- | --- |
| CD31 (Lin-) | BV421 | 1:100 | 390 | BioLegend | 102424 | AB_2650892 |
| CD19 (Lin-) | BV421 | 1:400 | 6D5 | BioLegend | 115537 | AB_10895761 |
| Viability | Aqua | 1:1000 | N/A | Thermo | L34957 | N/A |
| CD45 | BV605 | 1:150 | 30-F11 | BioLegend | 103140 | AB_2562342 |
| CD11b | BV711 | 1:300 | M1/70 | BioLegend | 101241 | AB_11218791 |
| CD3 | AF488 | 1:200 | 17A2 | BioLegend | 100210 | AB_389301 |
| CD34 | PerCP-Cy5.5 | 1:100 | HM34 | BioLegend | 128607 | AB_1279222 |
| F4/80 | PE-Cy7 | 1:250 | BM8 | BioLegend | 123114 | AB_893478 |
| CD29 | APC-Cy7 | 1:100 | HMB1-1 | BioLegend | 102226 | AB_2128076 |

## Notes

### Competing Interest Statement

The authors have declared no competing interest.

