## Supplemental Data for "Immune checkpoint inhibitors amplify type 2 immune mediated repair by pro-regenerative scaffolds"

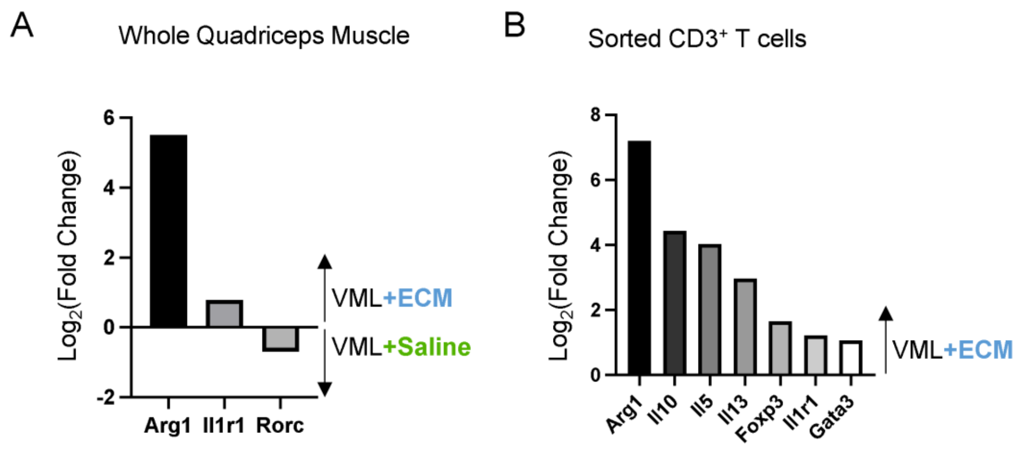


**<u>Fig S1</u>. Type 2 immune-related transcriptional signature in 1-week ECM-treated wounds using NanoString PanCancer Immune Profiling panel.**

**(A)** Expression of type 2 immune-related genes (*Arg1, Il1r1*) in bulk muscle tissue of ECM-treated VML injuries relative to saline-treated controls.

**(B)** Expression of $T_H2$-related (*Arg1, Il5, Il13, Il1r1, Gata3*) and Treg-related (*Il10, Foxp3*) in sorted $CD3^+$ T cells from ECM-treated VML injuries relative to saline-treated controls.

Data presented as $\log_2$(fold change) and only depicting significant differentially expressed genes (adj. $p<0.05$ and $|\log_2(FC)|>0.5$).

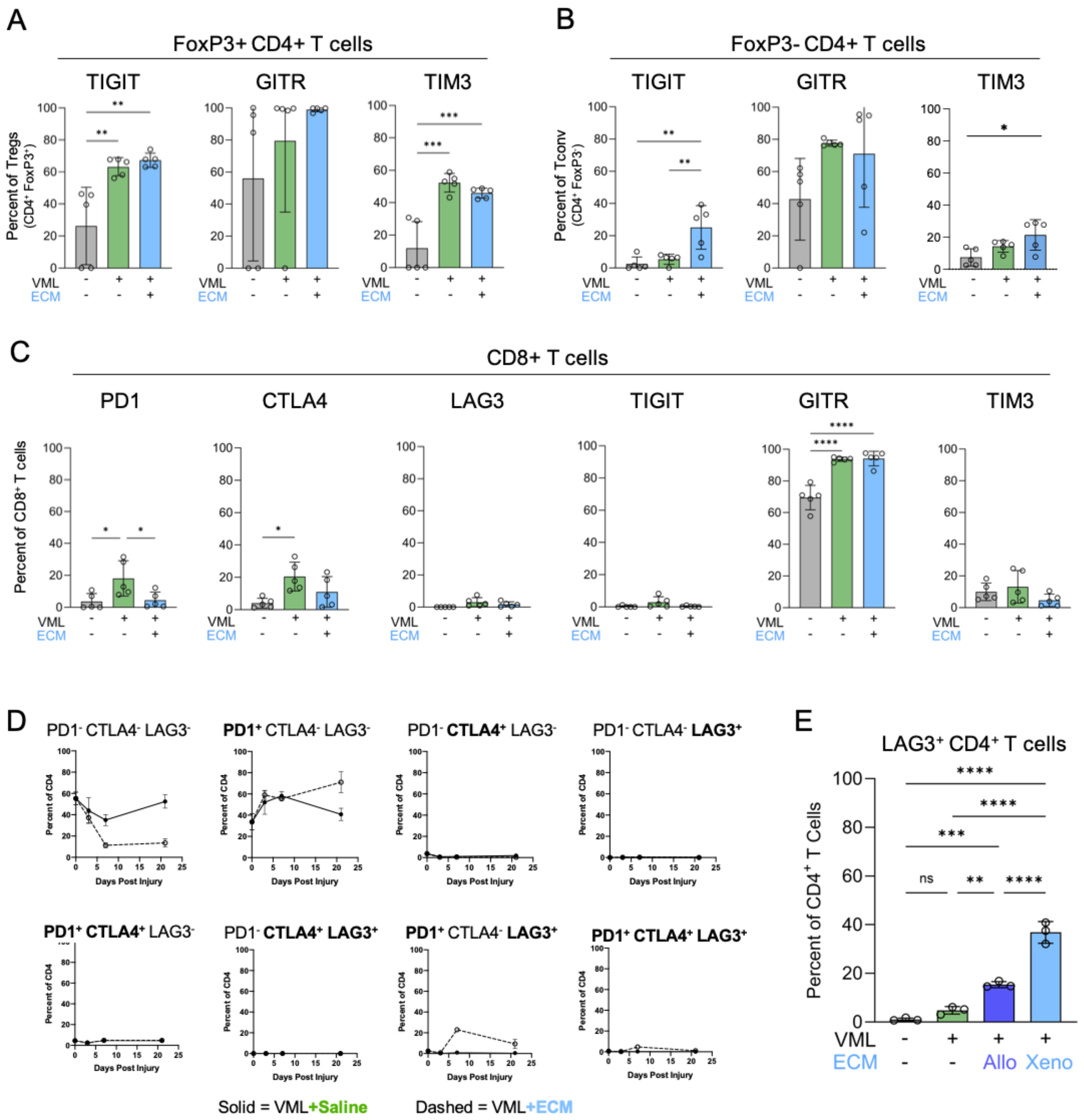


**Fig S2. CD4⁺ T cells expressed increased levels of immune checkpoints over a 3-week period following injury with both allogeneic and xenogeneic ECM implants.**

**(A)** Frequency of TIGIT, GITR, TIM3 protein expression in FoxP3⁺ Tregs in uninjured, saline-treated injuries, or ECM-treated injuries at 1-week.

**(B)** Frequency of TIGIT, GITR, TIM3 protein expression in Foxp3⁻ CD4⁺ T cells in uninjured, saline-treated injuries, or ECM-treated injuries at 1-week.

**(C)** Frequency of immune checkpoint protein expression in CD8⁺ T cells in uninjured, saline-treated injuries, or ECM-treated injuries at 1-week.

**(D)** Time course of PD1, CTLA4, and LAG3 expression in CD4⁺ T cells at 3-days, 1-week, and 3-weeks after saline-treated (solid) or ECM-treated (dashed) injury.

**(E)** Frequency of LAG3 expression in CD4⁺ T cells in xenogeneic vs allogeneic ECM-treated injuries at 1-week.

Data presented as mean±SD and analyzed using one-way ANOVA with Tukey's multiple comparisons test (A-C, E). NS $p>0.05$; * $p<0.05$; ** $p<0.01$; *** $p<0.001$; **** $p<0.0001$.

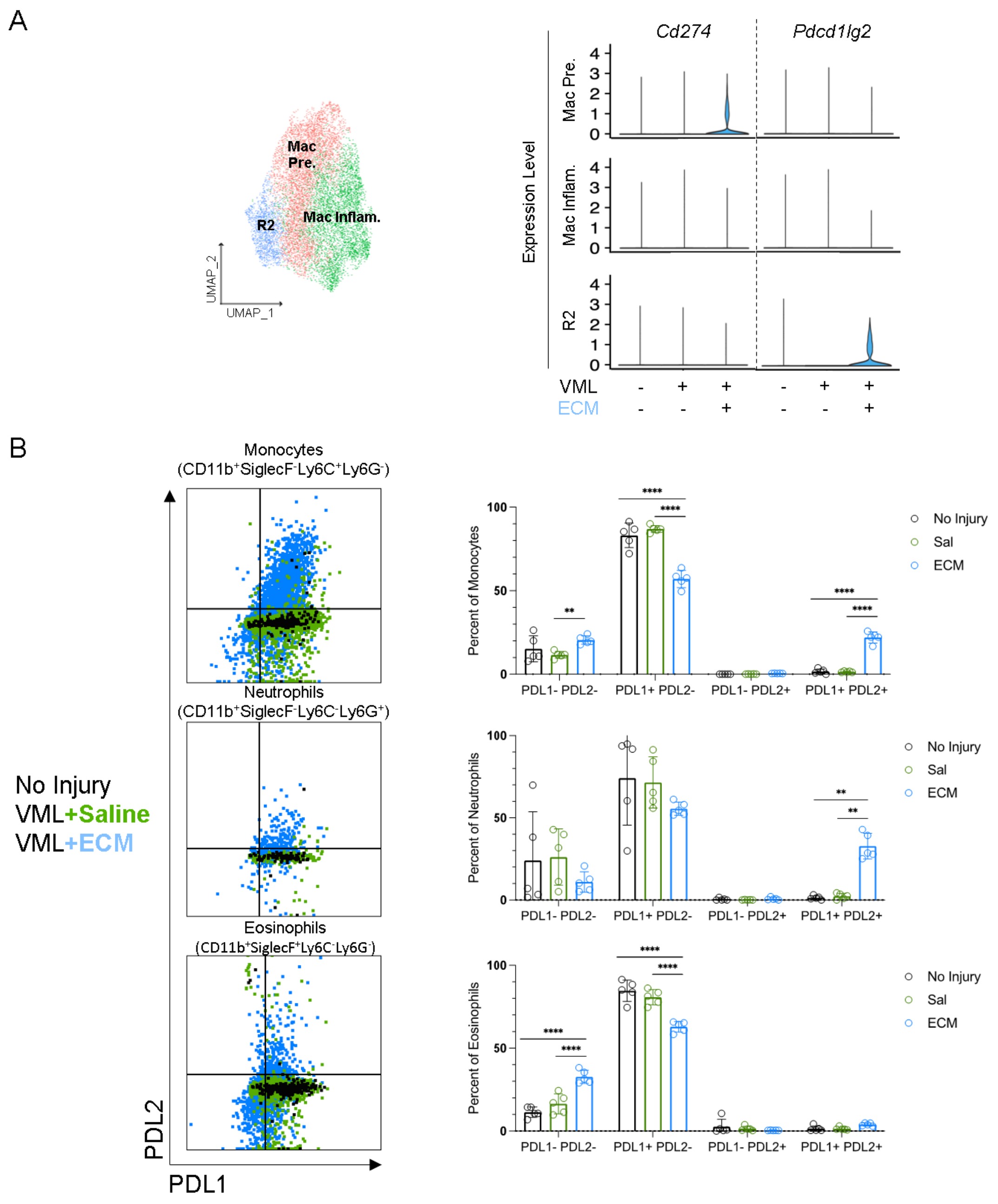


**<u>Fig S3.</u> PDL1 and PDL2 expression is present in ECM-treated wounds in some but not all myeloid cell types.**
**(A)** UMAP of macrophage subclusters from $CD45^+$ enriched scRNA-seq dataset collected from uninjured, saline-treated VML, and ECM-treated VML at 1-week. Violin plots for relative expression of *Cd274* (*Pdcd1lg1)* and *Pdcd1lg2*.
**(B)** PDL1 and PDL2 protein expression on monocytes, neutrophils, and eosinophils in saline-treated and ECM-treated wounds 1-week after injury.
Data presented as mean±SD and analyzed using two-way ANOVA with Tukey's multiple comparisons test (B). NS $p>0.05$; * $p<0.05$; ** $p<0.01$; *** $p<0.001$; **** $p<0.0001$.

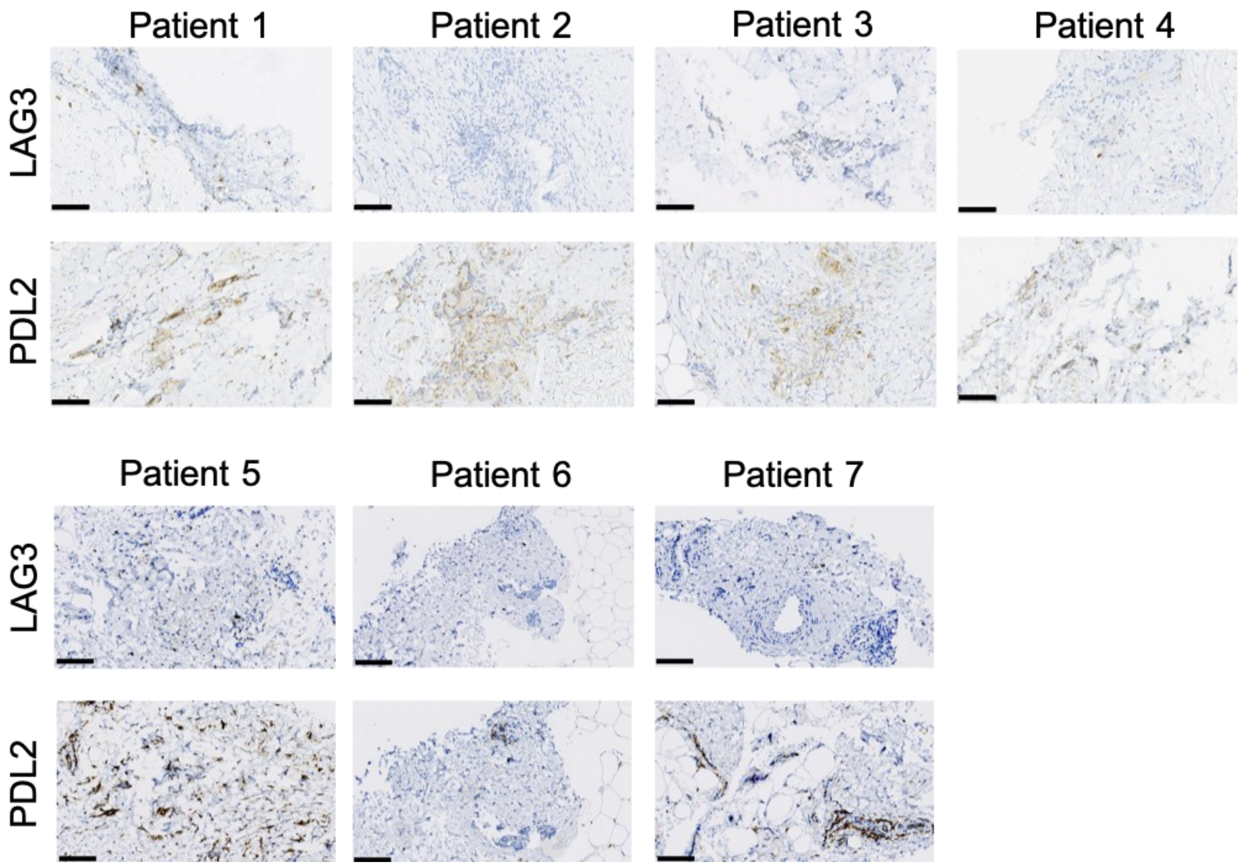


**<u>Fig S4</u>. Human ECM implants composed of acellular adipose tissue (AAT) demonstrate LAG3 and PDL2 staining on immunohistochemistry.**

**(A)** LAG3 and PDL2 immunohistochemistry in seven patients receiving acellular adipose tissue implants for soft tissue reconstruction. Scale bar: 100μm

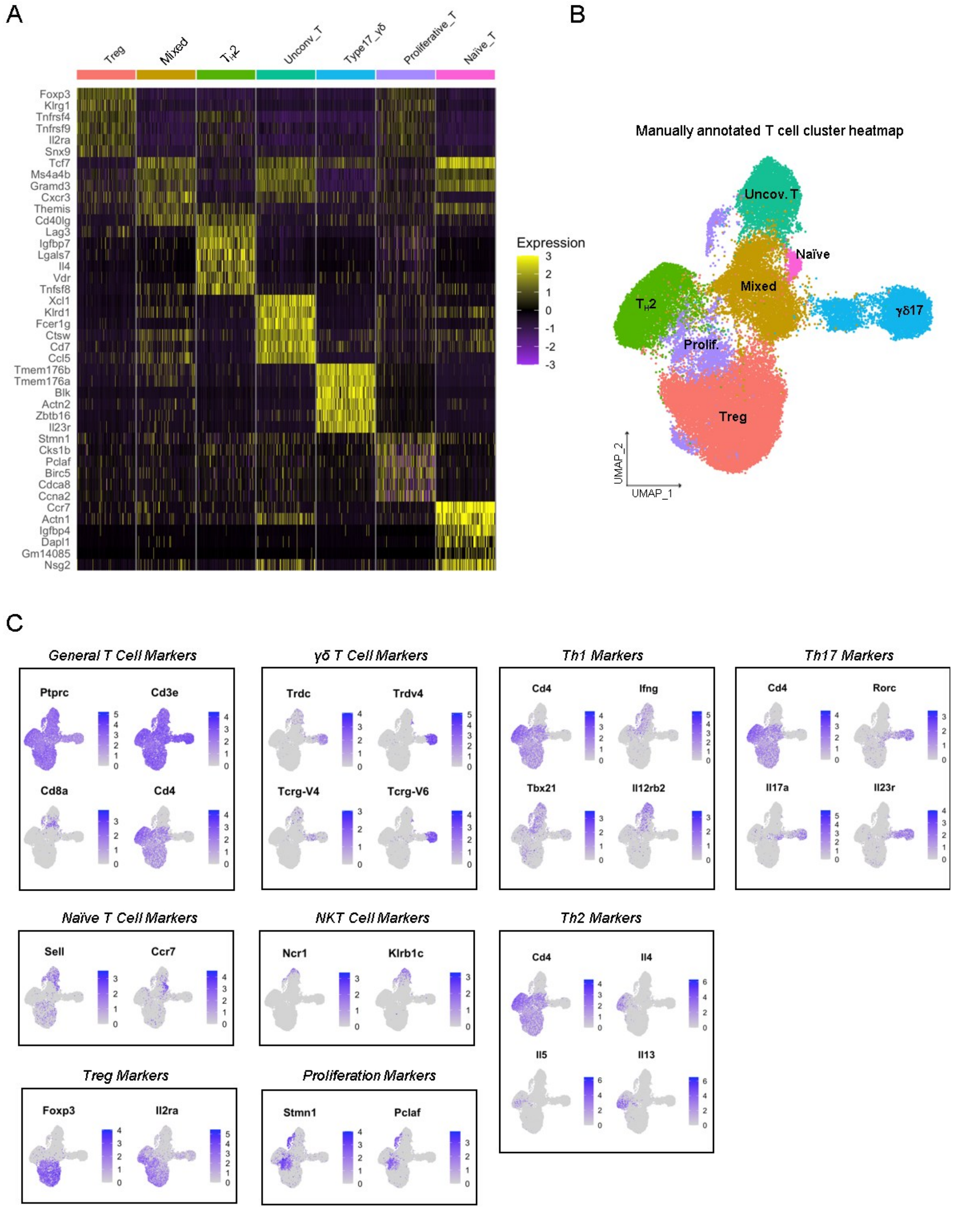


**<u>Fig S5</u>. Distinct T cell clusters identified in CD3$^+$ T cell scRNA-seq dataset of saline- and ECM-treated VML injuries at 1-week.**

**(A)** Heatmap of top differentially expressed genes across the seven T cell clusters.

**(B)** Representative UMAP projection of the seven T cell clusters identified in the dataset.

**(C)** Feature plots of canonical T cell subset markers to validate T cell clusters annotation.

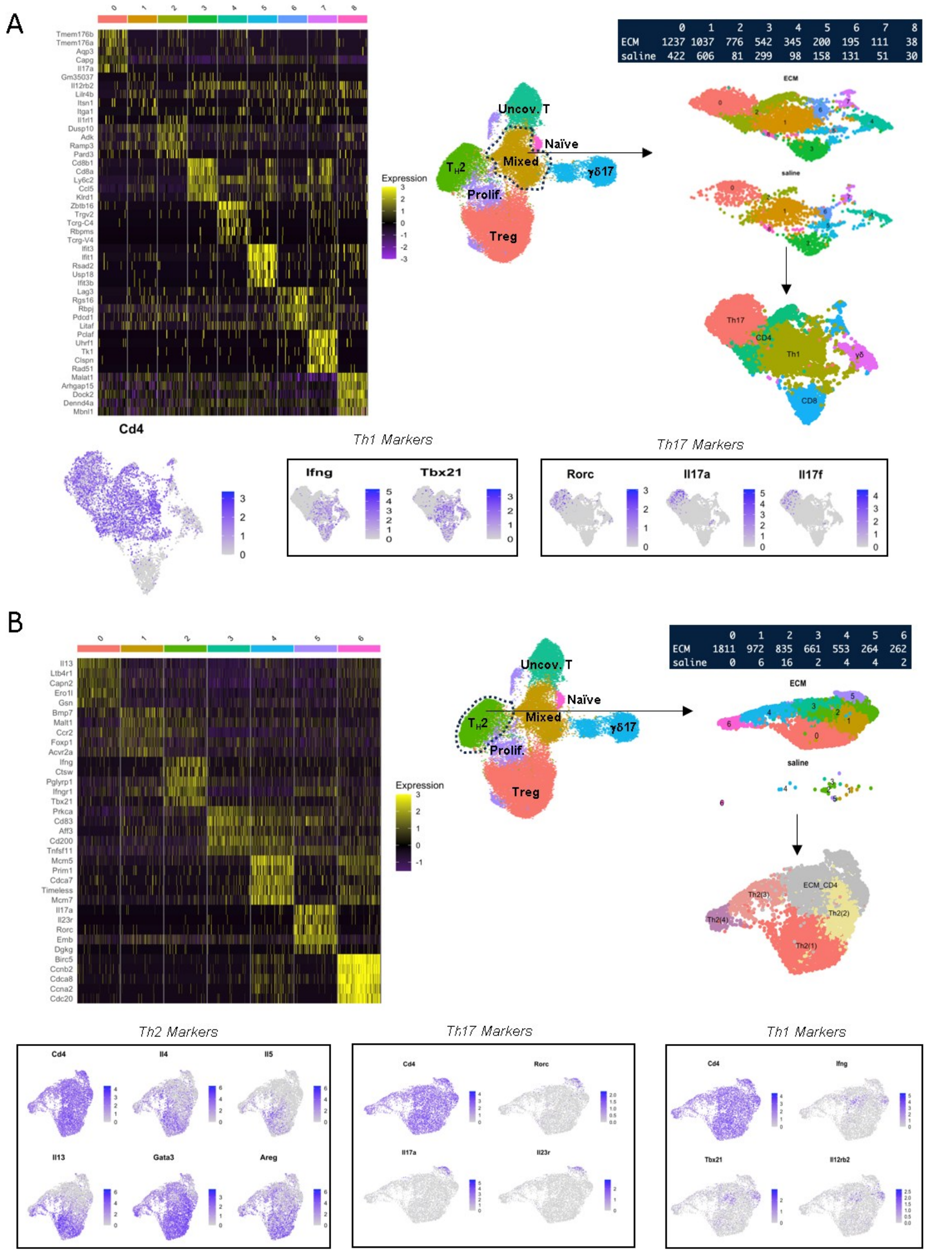


**<u>Fig S6</u>. Subcluster analysis shows distinct subclusters consistent with $T_H1$, $T_H2$, and $T_H17$ phenotypes.**

**(A)** Subcluster analysis was performed on the Mixed T cell cluster. Heat map of top differentially expressed genes per sub-cluster, sub-cluster UMAP, and sub-cluster cell counts by saline and ECM treatment condition. Feature plots of *Cd4*, canonical $T_H1$ (*Ifng, Tbx21*), and canonical $T_H17$ (*Rorc, Il17a, Il17f*) gene expression.

**(B)** Subcluster analysis was performed on the $T_H2$ cluster. Heat map of top differentially expressed genes per sub-cluster, sub-cluster UMAP, and sub-cluster cell counts by saline and ECM treatment condition. Feature plots for canonical CD4 helper subset markers clarify subcluster identity including $T_H2$ (*Il4, Il5, Il13, Gata3, Areg*), $T_H17$ (*Rorc, Il17a, Il23r*), and $T_H1$ (*Ifng, Tbx21, Il12rb2*) gene expression. For subsequent data analysis, only sub-clusters that strongly expressed $T_H2$ gene signature (0, 1, 4, and 6) were retained and relabeled Th2(1-4). Sub-clusters 2, 3, and 5 were excluded given their relatively high expression of $T_H17$ and $T_H1$ markers.

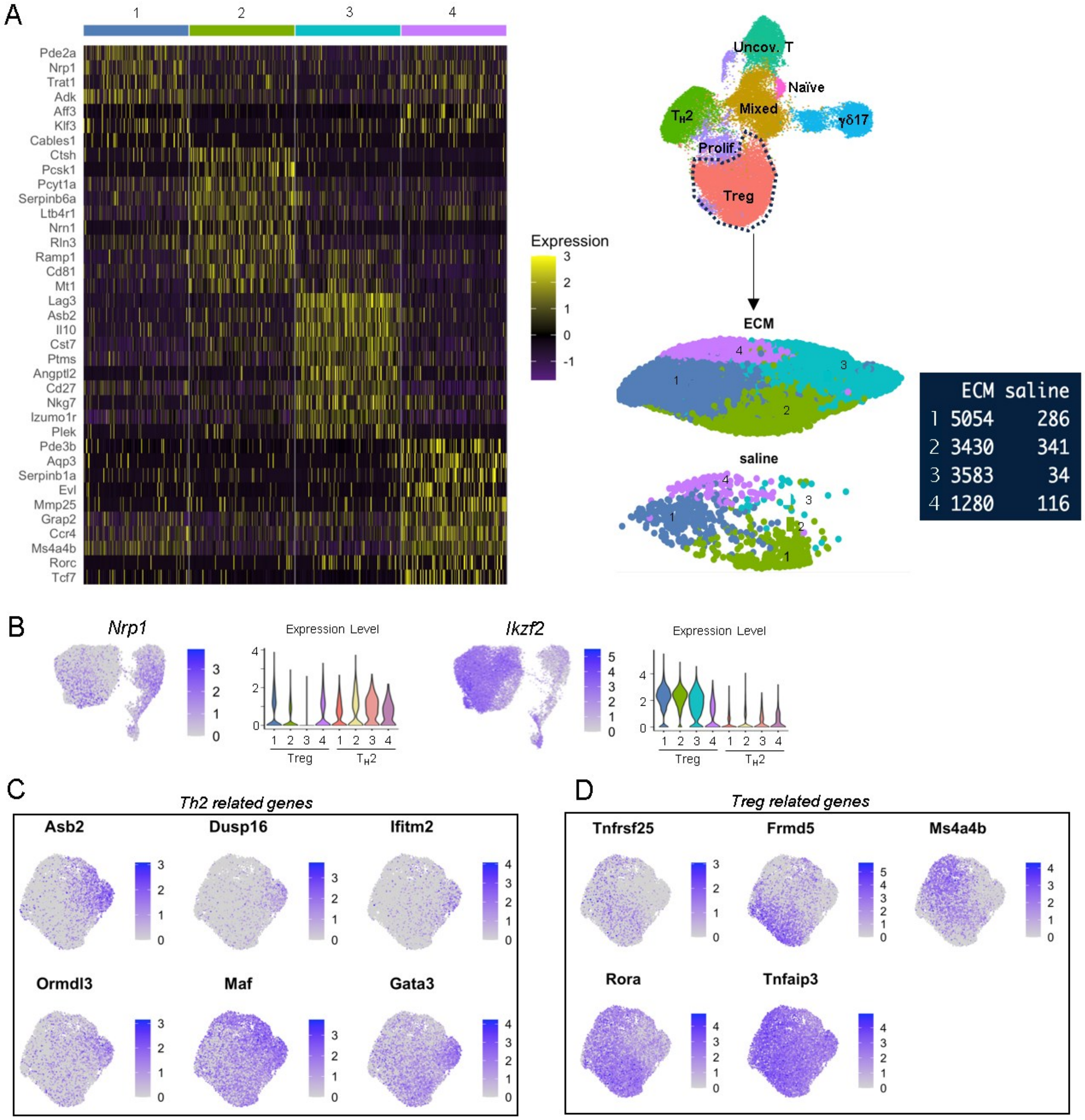


**Fig S7. Subcluster analysis shows distinct Treg subclusters with Treg_3 being most present in ECM-treated wounds.**

**(A)** Subcluster analysis was performed on the Treg cluster. Heat map of top differentially expressed genes per sub-cluster, sub-cluster UMAP, and sub-cluster cell counts by saline and ECM treatment condition.

**(B)** Feature plots for *Nrp1* and *Ikzf2* on UMAP from combined $T_H2$ and Treg populations. Diminished *Nrp1* expression and increased *Ikzf2* expression suggest Treg_3 to be peripherally derived Tregs.

**(C)** $T_H2$-related gene feature plots demonstrate overlap in expression on Treg_3 subcluster.

**(D)** Treg-related gene feature plots demonstrate overlap in expression on non-Treg_3 subclusters.

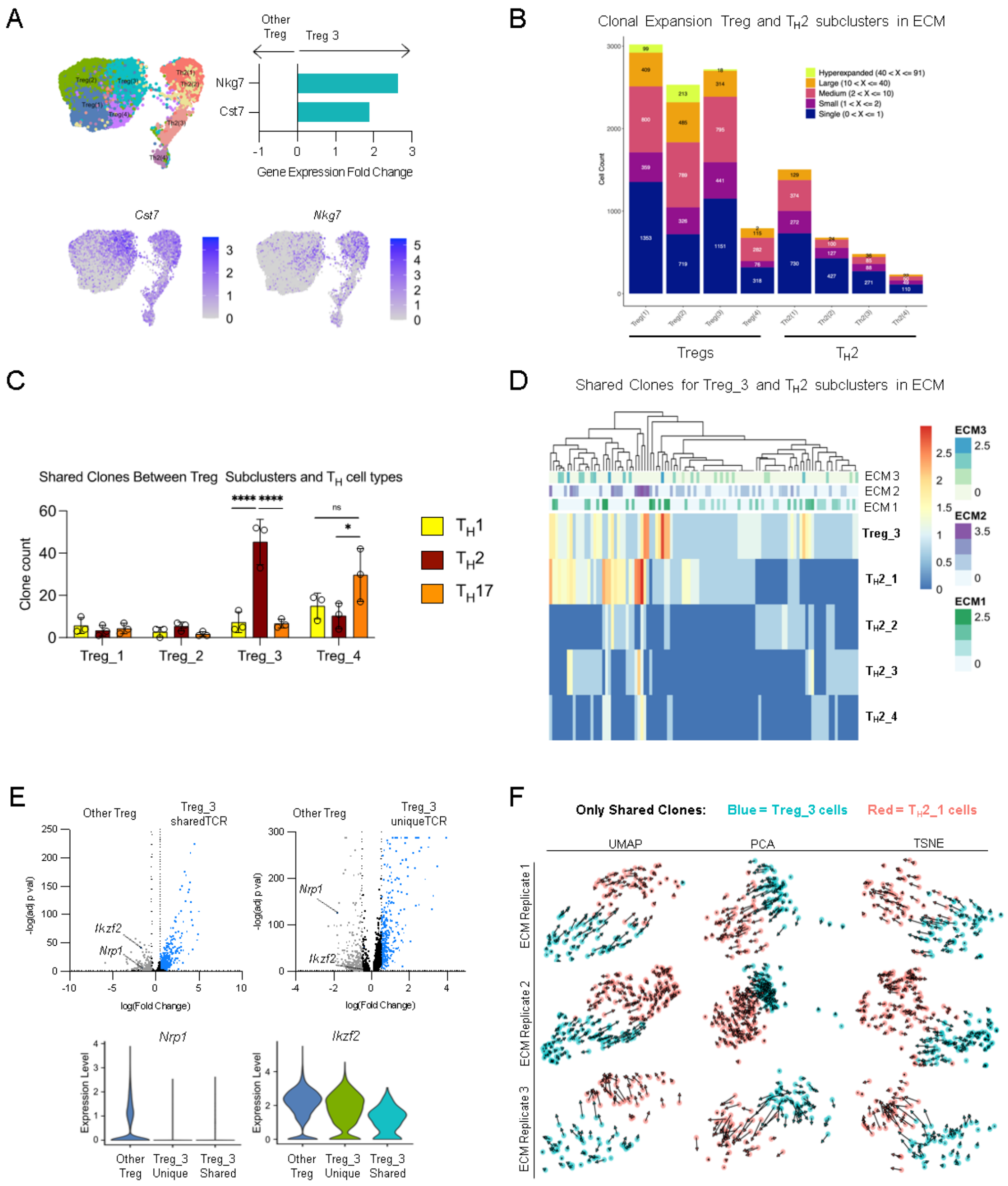


**Fig S8. Treg_3 cells share clones (TCR sequences) with T$_H$2 cells.**
**(A)** Expression of *Cst7* and *Nkg7* in Treg_3 relative to other Treg subclusters and T$_H$2 cells.
**(B)** Clonal expansion of Treg and T$_H$2 subclusters in ECM-treated wounds.
**(C)** Number of shared TCR clones between each Treg subcluster and T$_H$1, T$_H$2, and T$_H$17 cells. Treg_3 shares the most clones with T$_H$2 cells and Treg_4 shares the most clones with T$_H$17 cells.
**(D)** Heatmap of shared clones between Treg_3 and any T$_H$2 subcluster.
**(E)** Relative gene expression of Treg_3 cells with shared TCR sequences with T$_H$2 cells (Treg_3_sharedTCR) compared to other Treg subclusters. Relative gene expression of Treg_3 cells without shared TCR sequences with T$_H$2 cells (Treg_3_uniqueTCR) compared to other Treg subclusters. Violin plots for *Ikzf2* and *Nrp1.*
**(F)** RNA velocity diagrams of Treg_3 cells and T$_H$2_1 cells with shared TCR sequences by ECM replicate using various visual projection methods (UMAP, PCA, and TSNE)
Data presented as mean±SD and analyzed using two-way ANOVA with Tukey's multiple comparisons test (C). NS p>0.05; * p<0.05; ** p<0.01; *** p<0.001; **** p<0.0001.

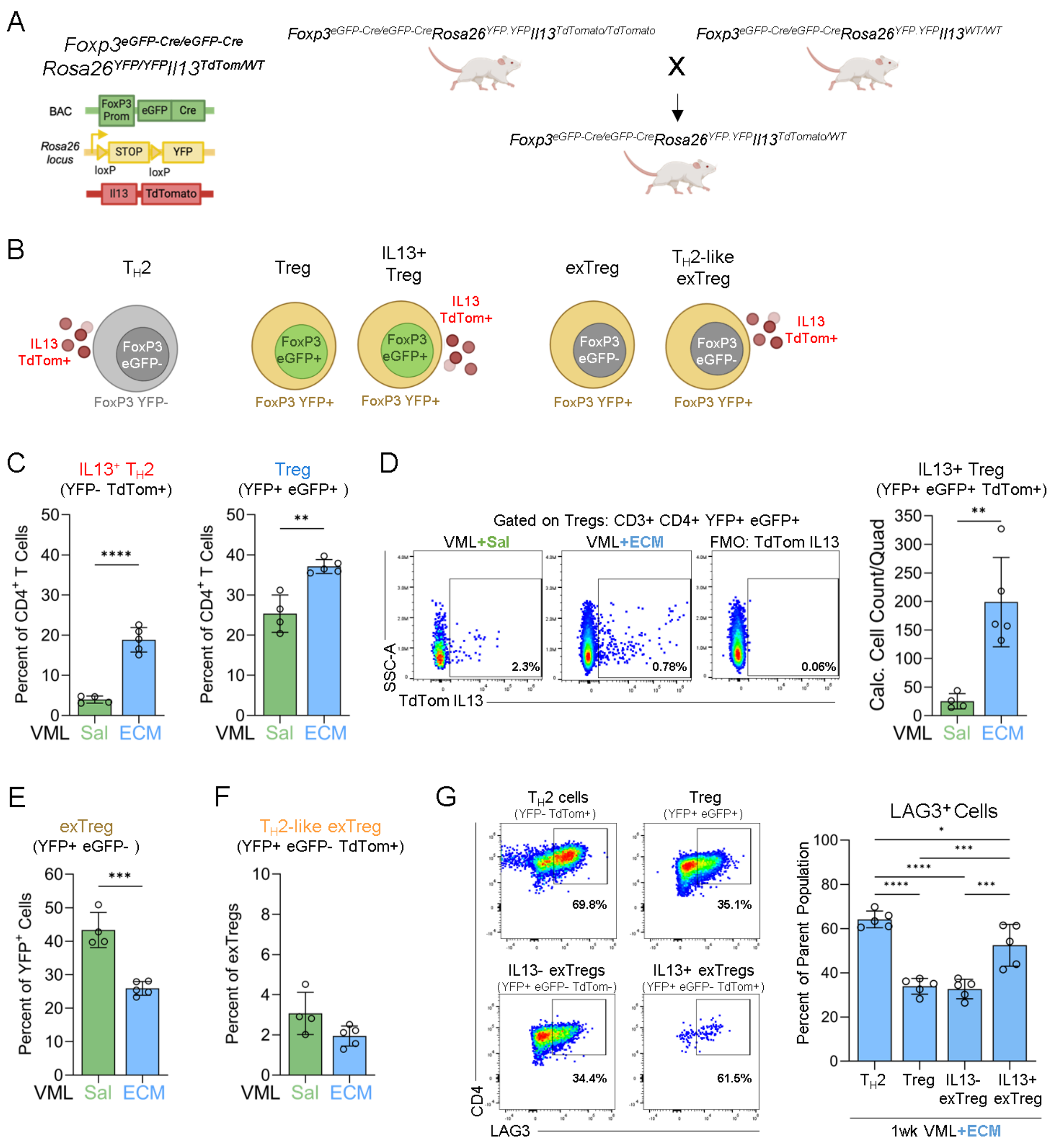


**Fig S9. FoxP3$^{eGFP/YFP}$IL13$^{TdTomato}$ reporter mice demonstrate a dynamic response between Tregs and T$_H$2 cells in ECM-treated wounds at 1-week.**

**(A)** Breeding scheme to generate FoxP3$^{eGFP/YFP}$IL13$^{TdTomato}$ reporter mice from available parent strains.

**(B)** Graphical representation of main CD4$^+$ T cell subsets that can be identified with this reporter strain, including IL13$^+$ T$_H$2 (YFP$^-$ TdTom$^+$), Tregs (YFP$^+$ eGFP$^+$), exTregs (YFP$^+$ eGFP$^-$), and T$_H$2-like exTregs (YFP$^+$ eGFP$^-$ TdTom$^+$).

**(C)** Frequency of T$_H$2 cells and Tregs out of CD4$^+$ T cells in saline- and ECM-treated injuries.

**(D)** Representative flow plots and quantification of IL13$^+$ Treg counts/quad in saline- and ECM-treated injuries.

**(E)** Frequency of exTregs out of CD4$^+$YFP$^+$ T cells in saline- and ECM-treated injuries.

**(F)** Frequency of T$_H$2-like exTregs out of exTregs in saline- and ECM-treated injuries.

**(G)** Representative flow plots and quantification of LAG3$^+$ expression in various T cell subsets (T$_H$2, Tregs, IL13$^-$ exTregs, and IL13$^+$ exTregs) in saline- and ECM-treated injuries.

Data presented as mean±SD and analyzed using unpaired two-tailed T-test (C-F) or one-way ANOVA with Tukey's multiple comparisons test (G). NS p>0.05; * p<0.05; ** p<0.01; *** p<0.001; **** p<0.0001.

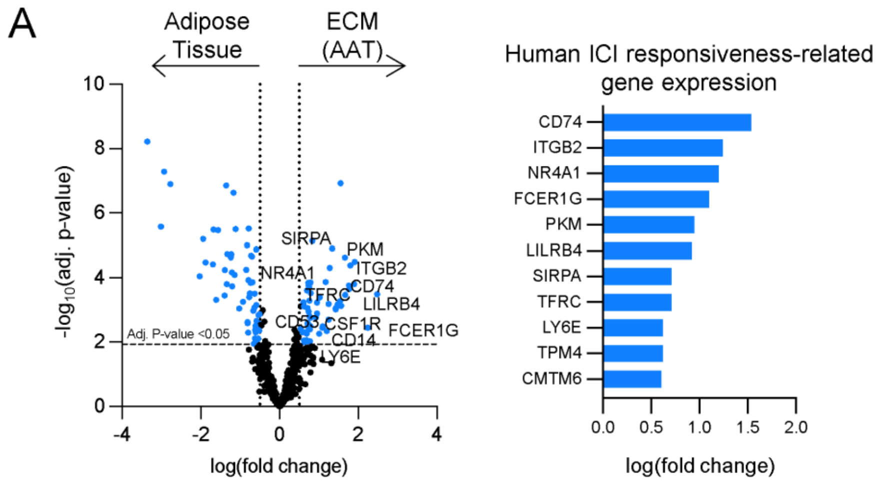


**Fig S10. Spatial transcriptomics comparing ICI responsiveness-related genes expression between ECM implants and native adipose tissue in clinical human samples.**

**(A)** Spatial transcriptomic analysis of differentially expressed genes between ECM implant regions and native adipose tissue in human biopsy samples. Listed genes have been implicated as markers of ICI responsiveness in clinical trials establishing the efficacy of ICI in different tumor settings.

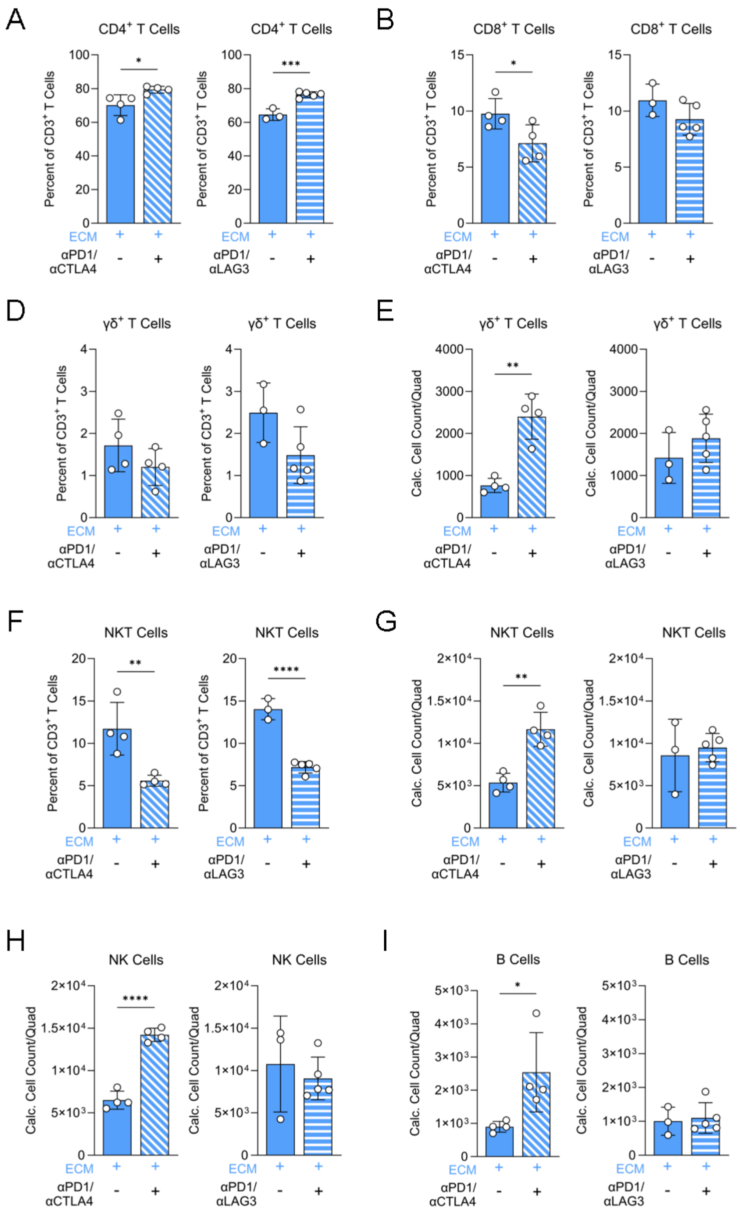


**Fig S11. Impact of combination anti-PD1/anti-CTLA4 or anti-PD1/anti-LAG3 treatment on various lymphocyte subsets in ECM-treated injuries at 3-weeks.**

**(A)** Frequency of $CD4^+$ T cells out of $CD3^+$ T cells with combination immune checkpoint inhibitors (ICI) in ECM-treated injuries.

**(B)** Frequency of $CD8^+$ T cells out of $CD3^+$ T cells with ICI in ECM-treated injuries.

**(C)** Quantification of $CD8^+$ T cell counts/quads with ICI in ECM-treated injuries.

**(D)** Frequency of γδ T cells out of $CD3^+$ T cells with ICI in ECM-treated injuries.

**(E)** Quantification of γδ T cell counts/quads with ICI in ECM-treated injuries.

**(F)** Frequency of NKT cells out of $CD3^+$ T cells with ICI in ECM-treated injuries.

**(G)** Quantification of NKT cell counts/quads with ICI in ECM-treated injuries.

**(H)** Quantification of NK cell counts/quads with ICI in ECM-treated injuries.

**(I)** Quantification of $B220^+$ B cell counts/quads with ICI in ECM-treated injuries.

Data presented as mean±SD and analyzed using unpaired two-tailed T-test (A-I). NS $p>0.05$; * $p<0.05$; ** $p<0.01$; *** $p<0.001$; **** $p<0.0001$.

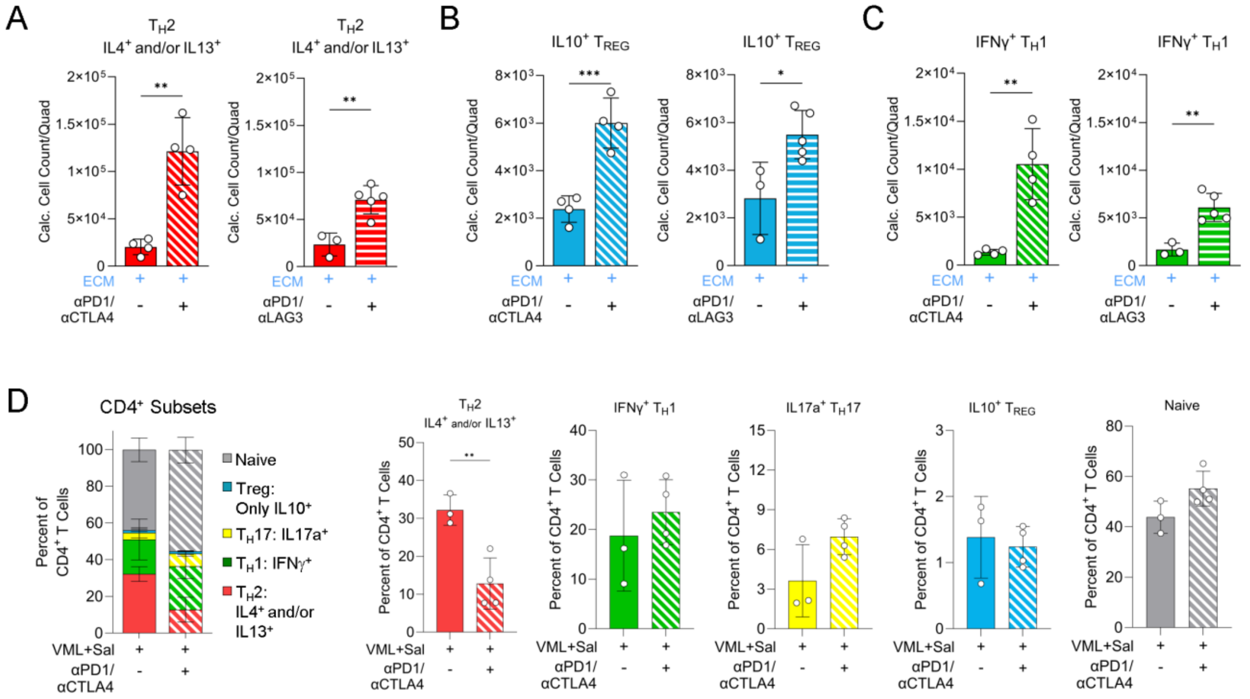


**Fig S12. Impact of combination anti-PD1/anti-CTLA4 or anti-PD1/anti-LAG3 treatment on CD4⁺ T cell cytokine expression in saline- or ECM-treated injuries at 3-weeks.**

**(A)** Quantification of $T_{H}2$ ($IL4^{+}$ and/or $IL13^{+}$) cell counts/quads with combination immune checkpoint inhibitors (ICI) in ECM-treated injuries.

**(B)** Quantification of $IL10^{+}$ "Treg" ($IL4^{-}$ $IL13^{-}$ $IFN\gamma^{-}$ $IL17a^{-}$ $IL10^{+}$) counts/quads with ICI in ECM-treated injuries.

**(C)** Quantification of $T_{H}1$ ($IFN\gamma^{+}$) cell counts/quads with ICI in ECM-treated injuries.

**(D)** Frequency of $CD4^{+}$ T cell subsets ($T_{H}1$, $T_{H}2$, $T_{H}17$, Treg, Naive) with anti-PD1/anti-CLTA4 in saline-treated injuries.

Data presented as mean±SD and analyzed using unpaired two-tailed T-test (A-D). NS $p>0.05$; * $p<0.05$; ** $p<0.01$; *** $p<0.001$; **** $p<0.0001$.

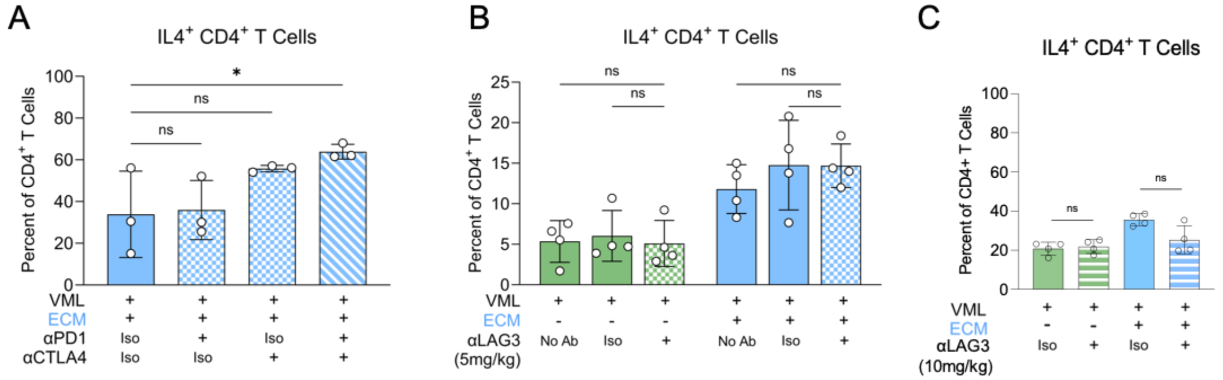


**Fig S13. Impact of monotherapy immune checkpoint inhibitors on IL4⁺ $T_H2$ responses in saline- or ECM-treated injuries at 3-weeks.**

**(A)** Frequency of $IL4^+$ $T_H2$ out of $CD4^+$ T cells with isotype, monotherapy anti-PD1, monotherapy anti-CTLA4, or combination anti-PD1/anti-CTLA4 in ECM-treated injuries.

**(B)** Frequency of $IL4^+$ $T_H2$ out of $CD4^+$ T cells with no treatment, isotype, or monotherapy anti-LAG3 (standard dose, 5 mg/kg) in saline- and ECM-treated injuries.

**(C)** Frequency of $IL4^+$ $T_H2$ out of $CD4^+$ T cells with isotype or monotherapy anti-LAG3 (increased dose, 10 mg/kg) in saline- and ECM-treated injuries.

Data presented as mean±SD and analyzed using one-way ANOVA with Dunnett's multiple comparison relative to isotype control (A) or two-way ANOVA with Tukey's multiple comparisons test (B-C). NS $p>0.05$; * $p<0.05$; ** $p<0.01$; *** $p<0.001$; **** $p<0.0001$.

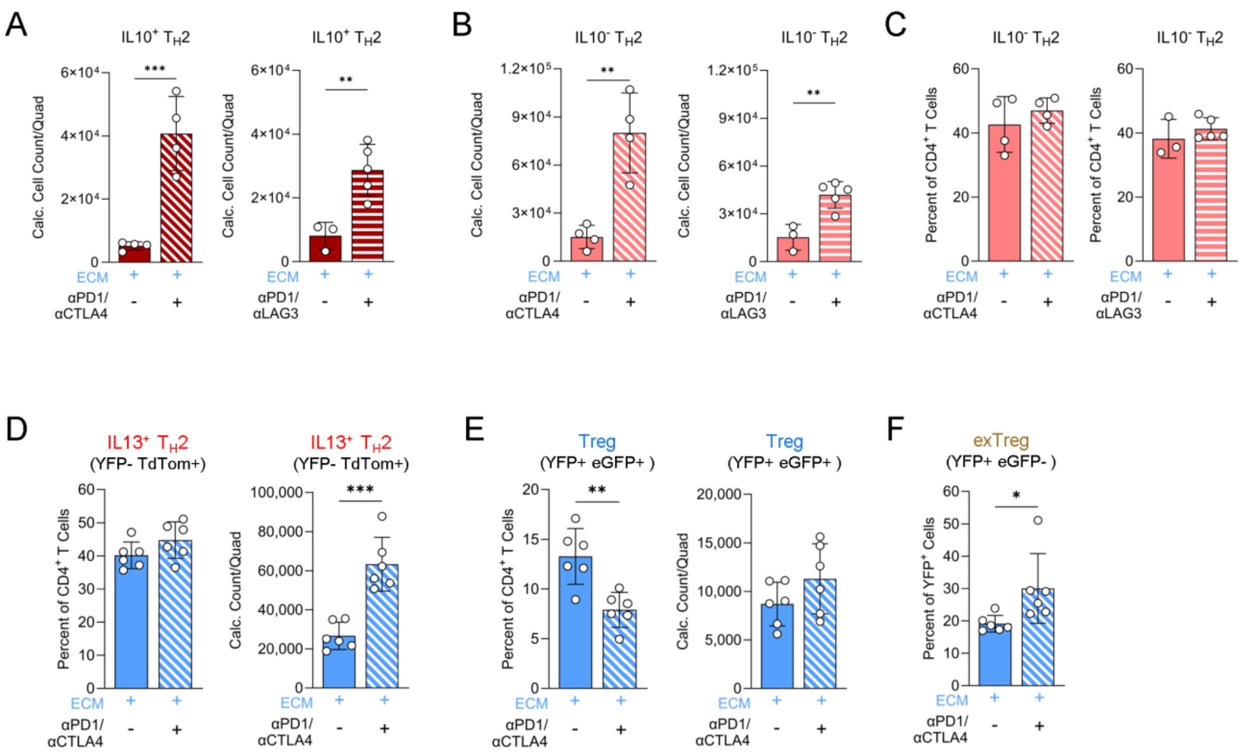


**<u>Fig S14</u>. Impact of combination anti-PD1/anti-CTLA4 or anti-PD1/anti-LAG3 treatment on IL10 production by T$_H$2 cells and Treg transition to T$_H$2-like exTregs in ECM-treated injuries at 3-weeks.**
**(A)** Quantification of IL10$^+$ T$_H$2 (IL4$^+$ and/or IL13$^+$) cell counts/quads with combination immune checkpoint inhibitors (ICI) in ECM-treated injuries.
**(B)** Quantification of IL10$^-$ T$_H$2 (IL4$^+$ and/or IL13$^+$) cell counts/quads with ICI in ECM-treated injuries.
**(C)** Frequency of IL10$^+$ T$_H$2 out of CD4$^+$ T cells with ICI in ECM-treated injuries.
**(D)** Frequency and quantification of IL13$^+$ T$_H$2 (YFP$^-$ TdTom$^+$) cell counts/quad with anti-PD1/anti-CTLA4 in ECM-treated injuries using FoxP3$^{eGFP/YFP}$IL13$^{TdTomato}$ reporter mice.
**(E)** Frequency and quantification of Treg (YFP$^+$ eGFP$^+$) cell counts/quad with anti-PD1/anti-CTLA4 in ECM-treated injuries using FoxP3$^{eGFP/YFP}$IL13$^{TdTomato}$ reporter mice.
**(F)** Frequency of exTregs (YFP$^+$ eGFP$^-$) out of YFP$^+$ cells with anti-PD1/anti-CTLA4 in ECM-treated injuries using FoxP3$^{eGFP/YFP}$IL13$^{TdTomato}$ reporter mice.
Data presented as mean±SD and analyzed using unpaired two-tailed T-test (A-F). NS $p>0.05$; * $p<0.05$; ** $p<0.01$; *** $p<0.001$; **** $p<0.0001$.

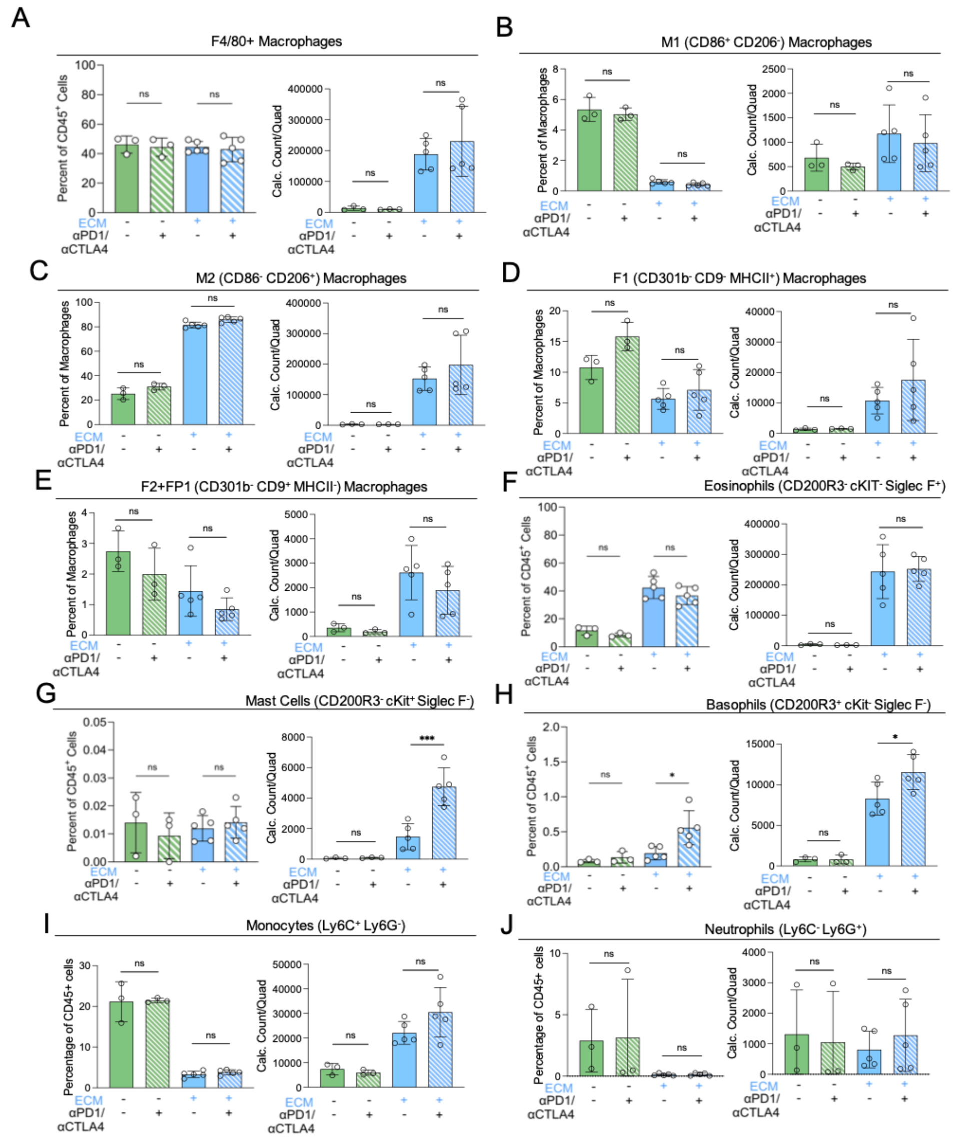


**<u>Fig S15</u>. Impact of combination anti-PD1/anti-CTLA4 on macrophage polarization and myeloid subsets in saline- or ECM-treated injuries at 3-weeks.**

**(A)** Frequency and count of F4/80$^+$ macrophages out of CD45$^+$ with anti-PD1/anti-CTLA4 (ICI) in saline- and ECM-treated injuries.

**(B)** Frequency and count of M1 (CD86$^+$CD206$^-$) macrophages with ICI in saline- and ECM-treated injuries.

**(C)** Frequency and count of M2 (CD86$^-$CD206$^+$) macrophages with ICI in saline- and ECM-treated injuries.

**(D)** Frequency and count of F1 (CD301b$^-$ CD9$^-$ MHCII$^+$) macrophages with ICI in saline- and ECM-treated injuries.

**(E)** Frequency and count of F2+FP1 (CD301b$^-$ CD9$^+$ MHCII$^-$) macrophages with ICI in saline- and ECM-treated injuries.

**(F)** Frequency and count of SiglecF$^+$ eosinophils with ICI in saline- and ECM-treated injuries.

**(G)** Frequency and count of CD200R3$^-$ cKit$^+$ mast cells with ICI in saline- and ECM-treated injuries.

**(H)** Frequency and count of CD200R3$^+$ cKit$^-$ basophils with ICI in saline- and ECM-treated injuries.

**(I)** Frequency and count of Ly6C$^+$ Ly6G$^-$ monocytes with ICI in saline- and ECM-treated injuries.

**(J)** Frequency and count of Ly6C$^-$ Ly6G$^+$ neutrophils with ICI in saline- and ECM-treated injuries.

Data presented as mean±SD and analyzed using two-way ANOVA with Tukey's multiple comparisons test (A-J). NS $p>0.05$; * $p<0.05$; ** $p<0.01$; *** $p<0.001$; **** $p<0.0001$.

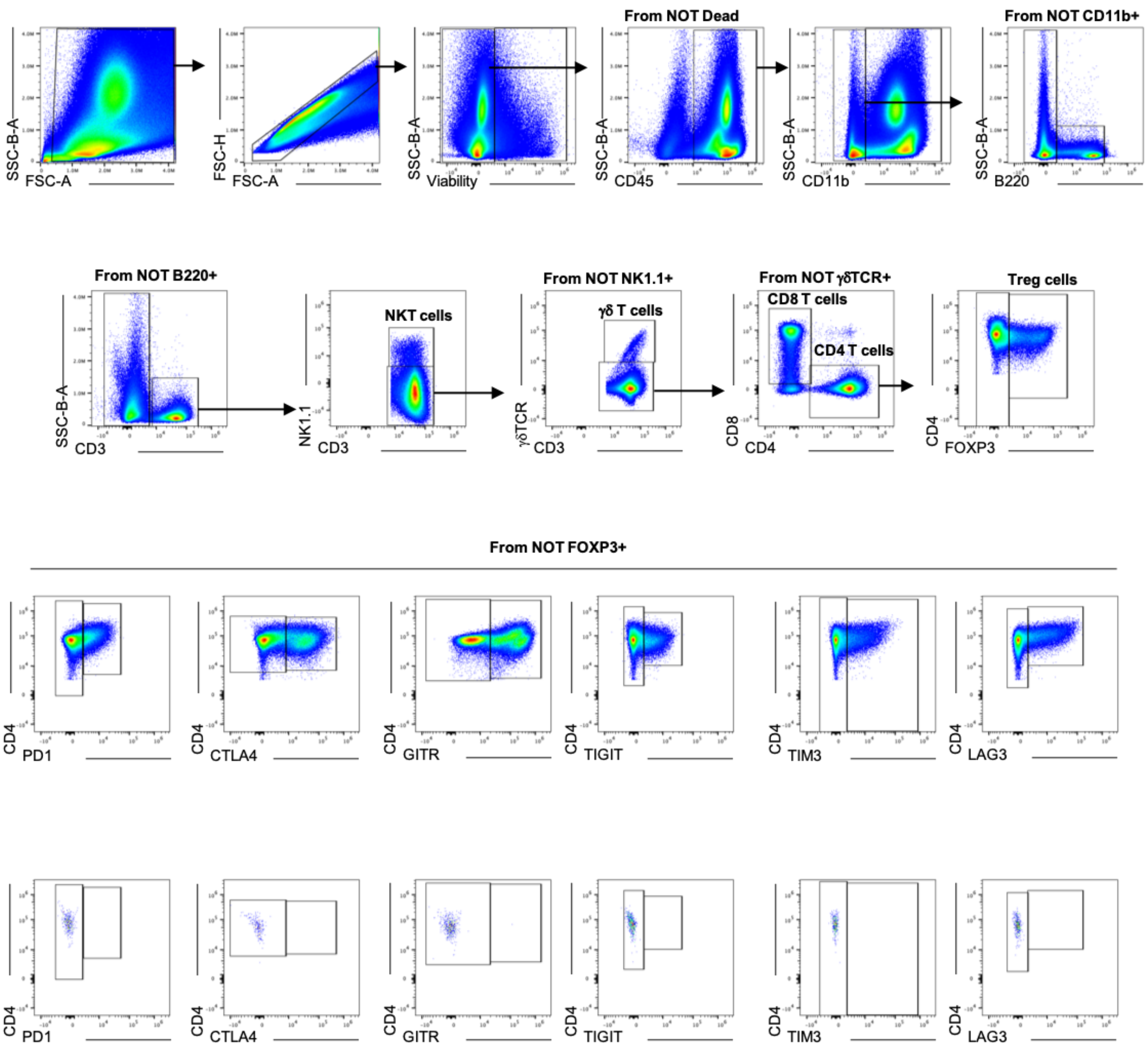


**<u>Fig S16</u>. Gating scheme for immune checkpoint expression panel with representative expression of immune checkpoints on conventional CD4$^+$ T cells and associated fluorophore-minus-one controls.**

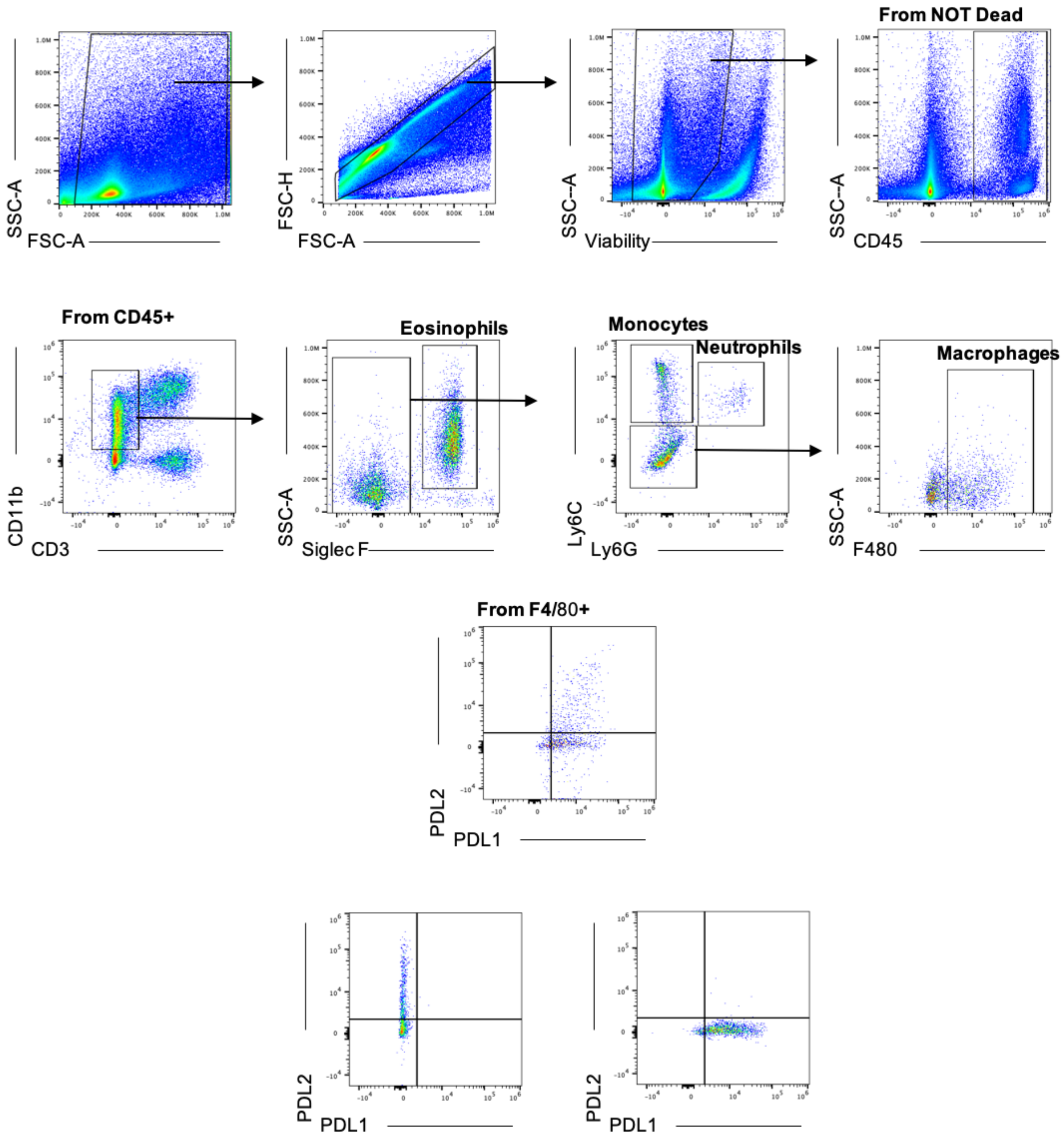


**<u>Fig S17</u>. Gating scheme for PDL1/PDL2 checkpoint expression panel with representative expression on conventional F4/80$^+$ macrophages and associated fluorophore-minus-one controls.**

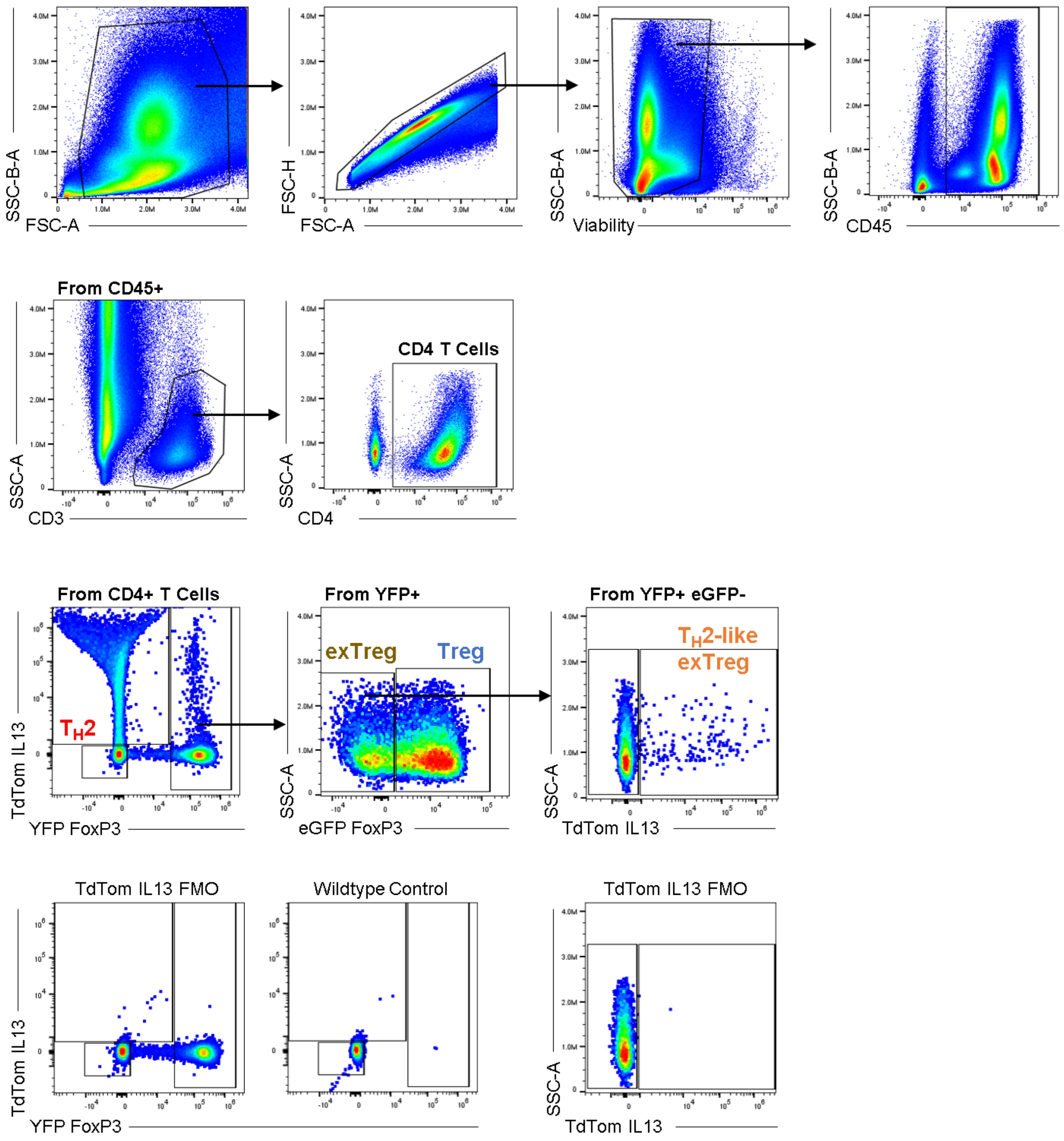


**<u>Fig S18</u>. Gating scheme for FoxP3$^{eGFP/YFP}$IL13$^{TdTomato}$ reporter strain panel to identify T$_H$2, Treg, exTreg, and T$_H$2-like exTreg populations and associated controls.**

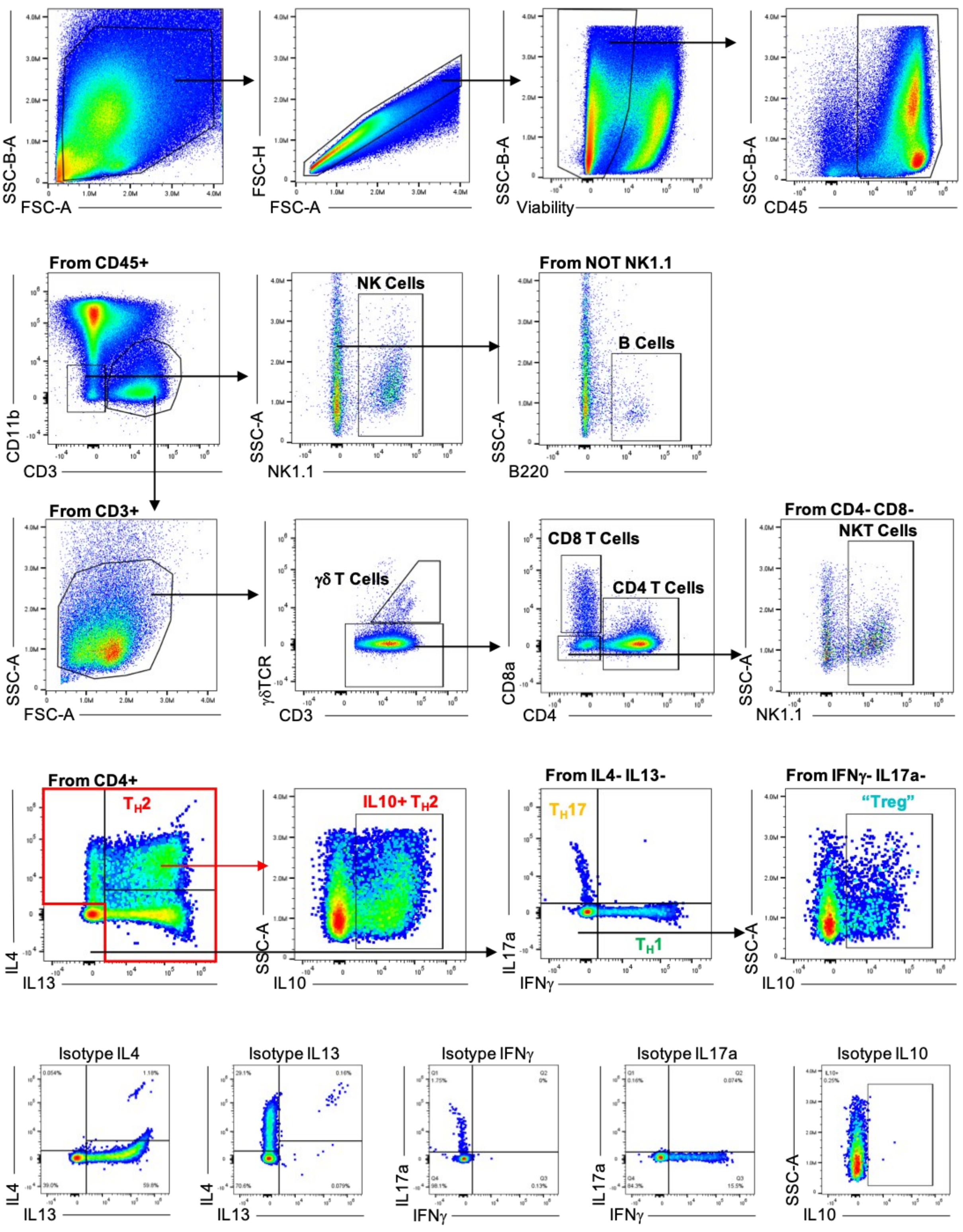


**Fig S19. Gating scheme for intracellular cytokine expression panel with representative expression of intracellular cytokines on conventional CD4$^+$ T cells and associated isotype controls.**

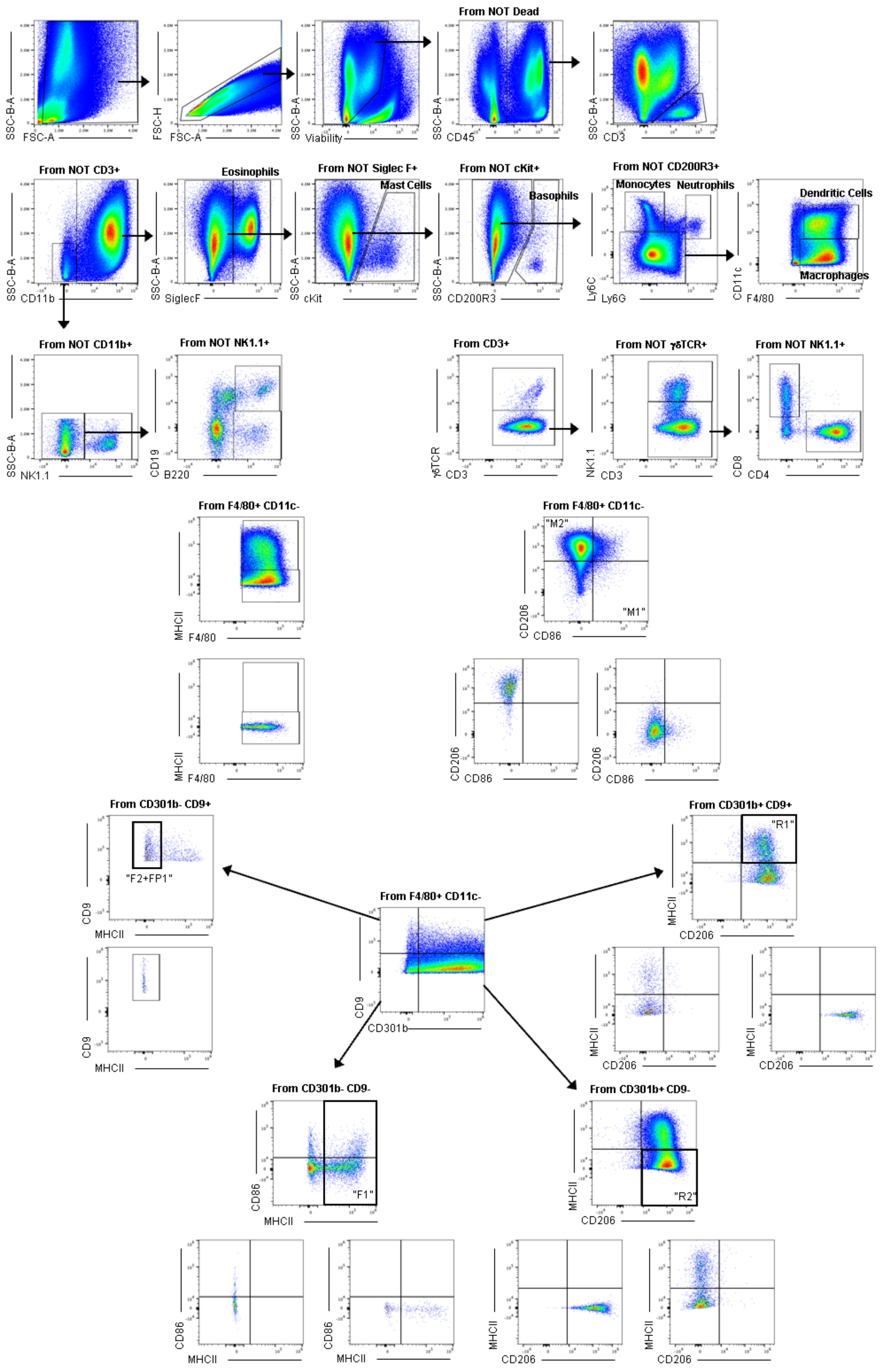


**Fig S20**. **Gating scheme for Pan-Immune panel with representative gating of macrophage subsets and associated fluorophore-minus-one controls.**
